# Protein entanglement misfolding determines divergent fates: proteasomal degradation or persistence in near-native misfolded states

**DOI:** 10.64898/2026.04.15.718748

**Authors:** Yang Jiang, Anushka Jain, Sina Ghaemmaghami, Edward P. O’Brien

## Abstract

A novel class of protein misfolding, involving changes in entanglement status, occurs across the cytosolic proteome of a bacterium and likely occurs in many other organisms. Here, we examine if this class of misfolding has measurable downstream consequences for protein homeostasis. Specifically, we test the hypothesis that proteins that misfold in this way are more likely to be degraded by the ubiquitin-proteasome system immediately after synthesis. We do this by cross-referencing protein structural information with ubiquitin mass spectrometry (Ubq-MS) data from human fibroblast cells. Ubq-MS identifies proteins that have been covalently modified with ubiquitin in a particular pattern and is a cellular signal for that protein to be degraded by the proteasome. We find that nascent proteins with native entanglements, which were previously shown to be twice as likely to misfold, are 93% (95% Confidence Interval: [44%, 160%]) more likely to be tagged with ubiquitin and targeted to the proteasome compared to proteins that do not contain such entanglements. Simulating the folding of these proteins using a coarse-grained model, we find that the ubiquitin-tagged proteins containing native entanglements are four times more likely to misfold than the non-ubiquitinated proteins that are devoid of entanglements. These results indicate that entanglement misfolding, primarily involving a failure to form native entanglements, leads to an increased likelihood that those proteins will be degraded in human cells. Finally, we estimate that approximately one-third of the globular proteome likely misfolds in this way but bypasses proteasomal degradation because their misfolded states are structurally similar to their native ensemble. These consequences for protein degradation are likely common across organisms as entanglement misfolding is inherent to the polymeric nature of proteins.

## Introduction

A series of studies from our lab (1–8) and others (9) have provided evidence that globular proteins can self-entangle their backbone segments into non-native geometries (Fig. 1A) creating off-pathway, kinetically trapped misfolded states. These states span a structural spectrum from grossly misfolded to highly native-like structures that are long lived (Fig. 1B), according to coarse-grained and all-atom simulations (1–9). The existence of these states provide an explanation for (*i*) how synonymous mutations can alter the long time scale structure and function of proteins (2, 5), (*ii*) the structural origin of stretched exponential folding kinetics and the kinetic partitioning mechanism (6), (*iii*) the nature of long-time scale intermediate states observed in limited proteolysis mass spectrometry experiments (1, 2, 6–8), and (*iv*) how some misfolded conformations of proteins can bypass the bacterial chaperone machinery GroEL, DnaK, DnaJ and HtpG (3, 8). Here, we explore whether the existence of these misfolded states, which are predicted to be widespread across the proteomes of organisms (1, 8), can affect the likelihood of degradation by the ubiquitin-proteasome system (UPS) in human fibroblast cells, for which high-quality data is available.

**Figure 1.**
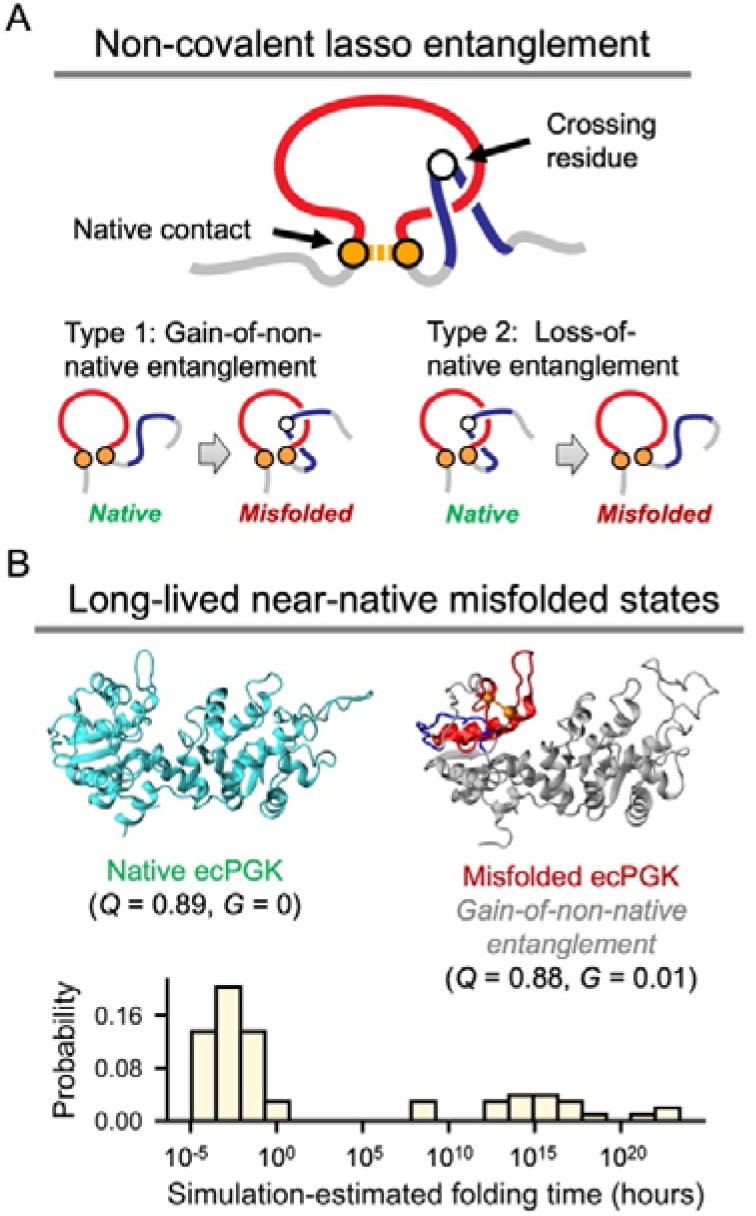
Protein misfolding involving changes in entanglement. A, Schematic of a non-covalent lasso entanglement (NCLE; top) and the two types of misfolding associated with entanglement changes (bottom). An NCLE is defined by a backbone loop (red) closed by one or more non-covalent contacts (orange), which is pierced one or more times by the N- or C-terminal backbone segments (blue). A protein may misfold either by gaining an NCLE absent from the native structure or by losing an NCLE present in the native structure. B, Near-native misfolded states involving entanglement changes often form long-lived kinetic traps, according to earlier studies (1–3, 6, 7). Shown is an example of a simulated misfolded structure of *E. coli* phosphoglycerate kinase (ecPGK) containing gain-of-non-native entanglements (top right), compared with its native structure (top left) (6). The corresponding *Q* and *G* values (see Methods) are shown below each structure. The probability distribution of simulation-estimated folding times from these misfolded states - obtained from our previous proteome-wide computational study for *E. coli* (1) - is shown at the bottom. These folding times were extrapolated by fitting the survival probability of unfolded proteins over time with a double-exponential function and we only reported the proteins (n = 73) whose fitting has a Pearson *R*^2^ > 0.9 (1). These misfolded states exhibit extremely slow folding kinetics, with folding times ranging from seconds to days or longer (1). (Longer time-scale estimates have much larger error bars.)

Non-covalent lasso entanglement (NCLE) misfolding may be a widespread class of monomeric protein misfolding that was only recently discovered (1). Non-covalent lassos are defined by two geometric components – a backbone loop closed by one or more non-covalent contacts between residues, and the threading of that loop by the N- or C-terminal backbone segments one or more times (Fig. 1A) (10). Simulations indicate two types of misfolding can occur involving these NCLEs (4, 6–8), either a non-native NCLE can form during the folding process, which is referred to as a ‘gain of entanglement’, or a native NCLE can fail to form during the folding process, referred to as ‘loss of entanglement’ or ‘failure to form mechanism’. Analysis of proteome-wide limited proteolysis mass spectrometry (LiP-MS) data of approximately 400 proteins from *E. coli* indicated the predominant misfolding mechanism involved the failure to form a native NCLE (8). This is a common misfolding mechanism because 70% of globular proteins in the human proteome contain one or more native NCLE (with a median number of 2) and therefore failing to form these should be more common than gaining a non-native NCLE (10). Further, there are more ways to misfold protein segments composing a native NCLE than segments not involving an NCLE. A native NCLE can misfold in two ways: it can gain an additional threading event leading to a gain of entanglement or it can fail to form. Other segments can only misfold one way: by gaining a non-native NCLE. The LiP-MS data indicate that proteins with native NCLE’s are 200% more likely to misfold than similar proteins that do not contain an NCLE (8). And that these natively entangled regions are 40% more likely to misfold compared to non-entangled regions (8).

In our previous simulations of protein folding we observed a distribution of misfolded entanglement states (1–8). Some were highly native like in structure, with most of their native contacts formed and the entanglement changes localized to a small portion of the primary structure (Fig. 1B). These states tend to have very long lifetimes, similar to and potentially longer than the native state (7). Others were less native like, having fewer native contacts formed, and greater structural deviations from the native ensemble.

A substantial fraction of newly synthesized proteins undergoes ubiquitination and proteasomal degradation during or shortly after translation (11–13). This process, mediated in part by the ribosome-associated quality control (RQC) pathway, serves as a key mechanism for removing translationally stalled or misfolded nascent proteins (14–16). We hypothesized that proteins containing native NCLEs, which are more likely to misfold (8), are also more likely to be targeted for UPS–mediated degradation immediately after synthesis. Advances in proteomics have enabled global identification and quantification of ubiquitinated proteins (Ubq-MS) (17, 18). A recent study combined Ubq-MS with time-resolved metabolic labeling to investigate the prevalence of ubiquitination of nascent proteins in human fibroblasts (19). Due to the pulsed nature of these experiments, protein ages were binned into 0 to 6 hours post-synthesis, 6 hours to 1.25 days, 1.25 to 4.25 days and 4.25 to 9.25 days. A key finding of that study was that most ubiquitinated proteins are substantially younger than the bulk proteome, with 90% of ubiquitination events occurring when the protein is less than a day old, indicating that a significant fraction of newly synthesized proteins is ubiquitinated and degraded. This proteome-wide dataset affords us the opportunity to test our hypothesis.

Here, we apply statistical analyses to detect if an association exists between the presence of native entanglements and ubiquitination status. We then use coarse-grained protein folding simulations as a test of their misfolding propensity. In the processes we discover a more complex and richer scenario exists in which entanglement misfolding can promote degradation of some proteins and apparently allow some misfolded proteins, which adopt conformations structurally similar to the native ensemble, to bypass degradation by the UPS.

## Results

### Properties of the datasets

In the mass spectrum data from Ref. (19) from human fibroblast cells there are 6,450 observable proteins common across all conditions (see Dataset S1), 2,280 have high quality AlphaFold structures (average pLDDT ≥ 85). Amongst high quality structures, 1,651 have one or more native NCLEs, 47 have covalent lassos and 15 have knots. Six-hours post labeling and with inhibition of the proteasome, 1,384 unique proteins were detected to be ubiquitinated, 1,299 were identified as being targeted for UPS degradation and 906 were identified as young-age ubiquitination – of which 507 have high quality AlphaFold structures. Amongst these high-quality structures, 432 have native NCLEs, 3 have covalent lassos and 1 has a knot.

### Natively entangled proteins are 93% more likely to be ubiquitinated

*E. coli* proteins containing non-covalent lasso entanglements are twice as likely to misfold as proteins that do not contain such entanglements (8). We therefore hypothesized that human proteins containing NCLEs in their native structures are more prone to misfolding. This hypothesis predicts that these proteins will be more likely to be targeted for UPS-mediated degradation following synthesis compared to proteins that lack such entanglements. To test this prediction, we use a Logistic regression (Eq. 1) to calculate the association between the presence of one or more native NCLEs and proteins that are ubiquitinated soon after synthesis (within 6 hours, denoted ‘YU’, standing for ‘young ubiquitinated’) as identified by human proteome birth dating data (see Methods). As a control group, we identified non-ubiquitinated (denoted ‘NU’) proteins as those that are mass-spec observable but exhibit no detectable ubiquitination across the proteome birth dating experiments. For the set of YU and NU proteins, we only included those that had AlphaFold structures (20, 21) with pLDDT ≥ 85 (see Dataset S2), and calculated the association using logistic regression (Eq. 1, contingency table shown in Table S1). Since protein length could be a confounding factor - longer proteins may inherently be more susceptible to misfolding (8) - we included the canonical protein length from UniProt (22) as another predictor in the regression model (Eq. 1), which allows us to separate out length and entanglement contributions.

Our YU and NU protein dataset consists of 1,490 unique proteins. Across all of these proteins 34% are both tagged with ubiquitin and of young age. 38% of proteins with one or more native NCLEs are ubiquitinated, compared to 20% among those without a native NCLE. We find that the presence of one or more native NCLEs is positively associated with young ubiquitination, with an odds ratio (OR) of 1.93 (95% CI: [1.44, 2.60], p-value < 0.0001, Wald test (23)). Protein length also has a positive association (OR = 1.60, 95% CI: [1.39, 1.84], p-value < 0.0001, Wald test), consistent with larger proteins being more likely to misfold regardless of whether they have a native entanglement or not (24, 25). We conclude that proteins with native entanglements are more likely to misfold following synthesis and be shunted to UPS-mediated degradation, increasing the odds of young-age ubiquitination by approximately 93% (=(1.93 − 1) × 100%; 95% CI: [44%,160%]).

As a sensitivity test, we repeated this analysis using the looser definition that YU proteins are those that are ubiquitinated within one day of synthesis (contingency table shown in Table S2). We again observe a positive association between NCLEs and young-age ubiquitination (OR = 1.77, 95% CI: [1.33, 2.36], p-value < 0.0001, Wald test, n = 1,528 unique proteins). Protein length still shows a positive association (OR = 1.66, 95% CI: [1.45, 1.91], p-value < 0.0001, Wald test, n = 1,528 unique proteins). Thus, our result of a positive association between native NCLEs and ubiquitination of nascent proteins is robust to reasonable variations in the definition of young proteins.

### Oligomeric status does not confound NCLE–ubiquitination association

We next tested whether oligomeric status (monomeric versus oligomeric, curated from Uniprot) influences our results (26) by including it as an additional confounder in the logistic regression model. We find this does not alter the association between entanglement and early ubiquitination (OR =1.94 versus 1.93 without this confounder), and monomeric status is not significantly associated (adjusted p = 0.61) with ubiquitination status. This indicates that the observed effect is not driven by oligomerization-dependent instability.

### Association is robust across independent ubiquitination datasets

To assess whether the observed association between native NCLEs and protein ubiquitination generalizes across independent human datasets, we analyzed three additional ubiquitination datasets, which we refer to as the Wagner dataset (27), and the Kim_btz and Kim_epox datasets (28). The latter two were generated in the same study using different proteasome inhibitors. Unlike the dataset from Ref. (19), which combines metabolic labeling with ubiquitin enrichment and therefore distinguishes young from old ubiquitinated proteins, these additional datasets quantify ubiquitinated proteins accumulated upon proteasome inhibition without age partitioning. However, the Meadow study showed that young ubiquitinated proteins are degraded substantially more rapidly by the proteasome than older ubiquitinated proteins (19). Consequently, proteasome inhibition preferentially enriches for ubiquitinated proteins from the young protein population, providing a rationale for comparing these datasets with the young-age ubiquitination analyses performed above. Using a consistent post-processing pipeline (see Methods), we identified ubiquitinated and non-ubiquitinated proteins in each data set that had corresponding high-quality AlphaFold structures and performed logistic regression analyses controlling for protein length.

We observed a significant positive association between native NCLEs and ubiquitination in both the Wagner dataset (OR = 3.11, 95% CI: [2.20, 4.39], p < 0.0001, Wald test; n = 821 unique proteins) and the Kim_btz dataset (OR = 1.85, 95% CI: [1.33, 2.59], p = 0.0003, Wald test; n = 839). The Kim_epox dataset showed a similar trend, but the association did not reach statistical significance (OR = 1.36, 95% CI: [0.92, 2.00], p = 0.1192, Wald test; n = 633). We attribute the lack of statistical significance in the Kim_epox dataset to its smaller sample size and consequently reduced statistical power. Consistent with this interpretation, further reducing the sample size using an alternative IPI-to-UniProt mapping (see Methods) resulted in weaker statistical significance in both the Kim_btz and Kim_epox datasets (see Table S3).

Overall, proteins containing native entanglements are estimated to be 85% to 211% more likely to be ubiquitinated across two of the three independent datasets, supporting the robustness of this association in the human proteome.

### Young ubiquitinated, entangled proteins are more prone to misfolding

To test our hypothesis that natively entangled proteins misfold more often – ultimately enhancing their susceptibility to UPS-mediated degradation - we performed coarse-grained (CG) molecular dynamics (MD) simulations (2) on a representative set of 20 human proteins (see Table 1) and calculated their propensity to misfold at the end of 2 μs of simulation time. To construct this set, we randomly selected from the experimental set of YU proteins 10 globular proteins containing native NCLEs (hereafter, denoted YU-E proteins) and randomly selected from the experimental set of NU proteins 10, length-matched globular proteins lacking native entanglements (denoted NU-NE proteins; see Methods). Our hypothesis predicts the YU-E set of proteins will have a higher probability of misfolding compared to the NU-NE set.

**Table 1.** Randomly selected globular proteins for simulation.

| YU-E |  | NU-NE |  | NU-E |  |
| --- | --- | --- | --- | --- | --- |
| Uniprot ID | Length (AA) | Uniprot ID | Length (AA) | Uniprot ID | Length (AA) |
| Q0PNE2 | 266 | Q8WV22 | 266 | P07738 | 259 |
| P16152 | 277 | A6NDU8 | 294 | Q9UIV1 | 285 |
| P29218 | 277 | P30711 | 240 | Q9UBP6 | 276 |
| P00491 | 289 | Q9BU89 | 302 | Q9Y316 | 297 |
| P19623 | 302 | Q13825 | 339 | Q63HM1 | 303 |
| A8MXV4 | 375 | Q6NVY1 | 386 | P39748 | 380 |
| P04350 | 444 | P16520 | 340 | Q6DKJ4 | 435 |
| P31150 | 447 | Q8IV38 | 441 | Q9HB40 | 452 |
| O95394 | 542 | P02774 | 474 | Q96C11 | 551 |
| P52888 | 689 | Q12996 | 717 | Q9H6R3 | 686 |

These proteins were thermally unfolded and then driven to refold via a temperature quench to 310 K in the simulations (see Methods). To calculate their misfolding propensity we first identified the metastable states they populated in *Q* – *G* space, in which *Q* is the fraction of native contacts (Eq. 2) and *G* is the fraction of native contacts that exhibit a change in entanglement (Eq. 3). In this space, conformations were assigned to macrostates (Fig. 2A, C and E, see Methods). For example, the YU-E protein Carbonyl Reductase [NADPH] 1 (Uniprot ID: P16152) (Fig. 2A) populated 11 metastable states in *Q*-*G* space. The native basin is State 11 (S11, Fig. 2A), centered at Q = 0.94 and G = 0. States with G values greater than 0.005 exhibit a change in entanglement misfolding. Therefore, this protein has 5 misfolded states involving a change in entanglement (see Fig. 2B). In contrast, the NU-NE protein RAB7A-interacting MON1-CCZ1 Complex Subunit 1 (Uniprot ID: A6NDU8) exhibits no misfolded state involving a change in entanglement (Fig. 2C and D). Metastable states for other YU-E and NU-NE proteins can be found in Supplementary Figs. 1 to 20. Partitioning the space in this way, we can directly sum the probabilities of these different metastable states being observed in the last 100 ns of the simulations and calculate the protein’s probability of misfolding (Eq. 4).

**Figure 2.**
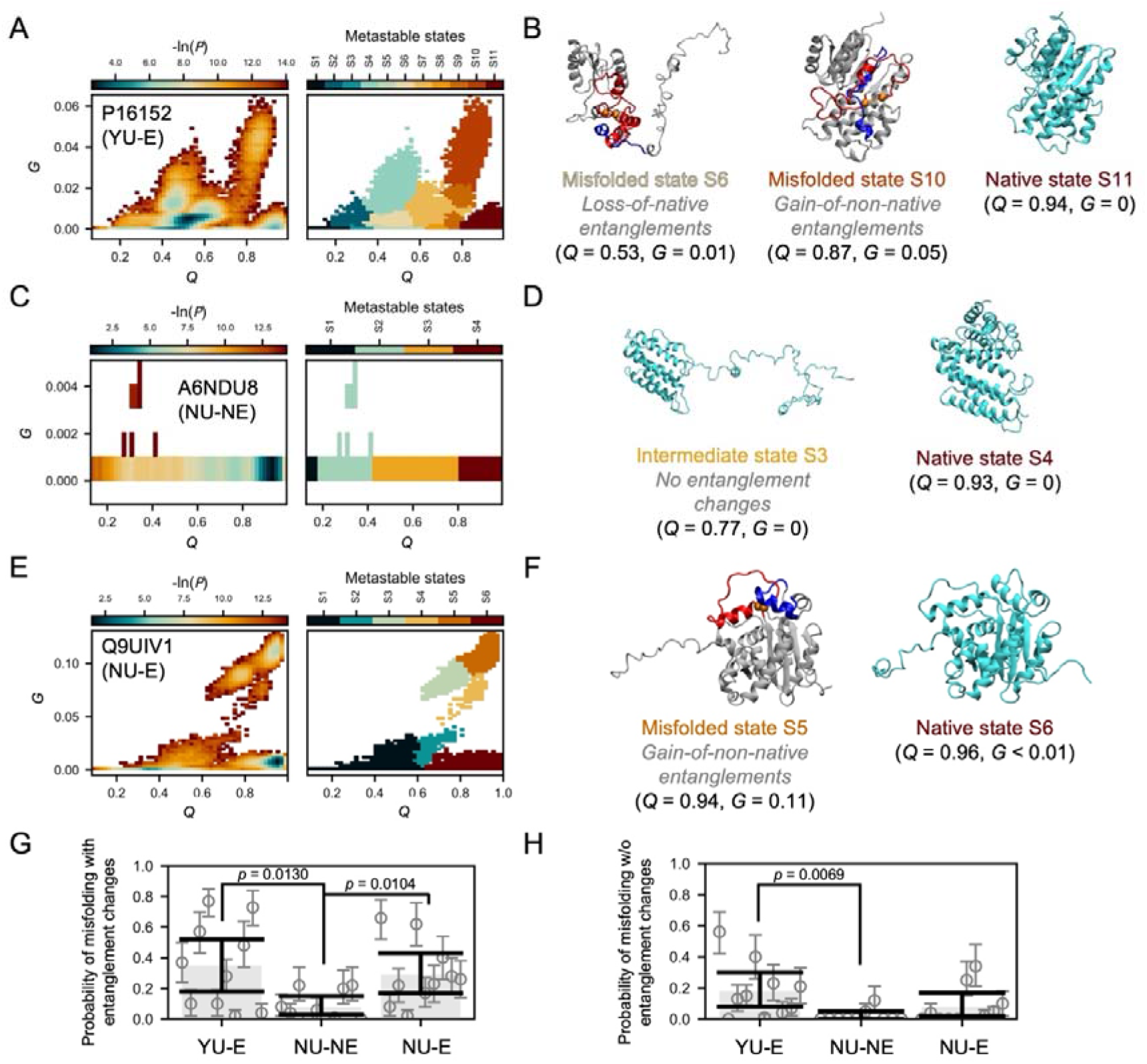
Misfolded structural ensembles of representative YU–E, NU–NE, and NU–E proteins from temperature-quench simulations. The probability distribution (-ln(*P*) ) in the Q – G space (left) and the metastable states assigned on the temperature quench refolding simulation structures (right) for the representative YU-E protein P16152 (A), NU-NE protein A6NDU8 (C) and NU-E protein Q9UIV1 (E). Representative misfolded and native structures are displayed in panels B, D, and F, respectively. In the misfolded states, the closed loop and threading segment are highlighted in red and blue, respectively, with loop-closing contacts shown as orange spheres. The Q and G values at the center of each metastable state are indicated below the corresponding structures. Probability of misfolding with (G) and without (H) entanglement changes (see Eq. 4). For each protein, the average misfolding probability across simulation trajectories is shown as gray circles, with 95% confidence intervals (CIs) indicated by error bars (bootstrap resampling, 10^6^ iterations). Group-level averages for each protein group are shown as gray bars, with 95% CIs estimated by hierarchical bootstrap resampling (10^6^ iterations, see Methods). P-values are shown only for comparisons with p < 0.05 (two-sided hierarchical bootstrap test, 10^6^ iterations, see Methods).

We find the YU-E proteins, on average, misfold with a change of entanglement 4.4-times that of NU-NE proteins (Probability of misfolding = 0.35, 95% CI = [0.18, 0.52] versus 0.08, 95% CI = [0.03, 0.15]; p-value = 0.0130, two-sided hierarchical permutation test for 10^6^ times, see Fig. 2G). As a comparison, we also examined the probability of adopting non-native states that do not involve a change in entanglement status. We found the YU-E proteins also exhibited a probability 9-fold that of NU-NE proteins (Probability of misfolding = 0.18, 95% CI = [0.08, 0.30] versus 0.02, 95% CI = [0.00, 0.05]; p-value = 0.0069, two-sided hierarchical permutation test for 10^6^ times, see Fig. 2H). This is not surprising, as misfolded states involving entanglement changes often require partial or full unfolding before the protein can refold into its native state (2, 4). During this back-tracking process, intermediate states without entanglement changes can be generated along the folding pathway.

To test whether these misfolded entanglement states are long-lived, we extended the simulation time to 20 μs and calculated the misfolding propensity at 1-μs intervals. The resulting time courses show that YU-E proteins consistently exhibit a higher propensity to misfold than NU-NE proteins throughout the extended simulations, with differences remaining statistically significant at most time points examined (p-value < 0.05; Fig. S21A and B).

These results indicate that native entanglements dramatically increase the likelihood of misfolding involving entanglement changes. This observation offers a mechanistic basis for the increased propensity of entangled proteins to be targeted to the UPS following translation.

### Non-ubiquitinated, entangled proteins are more likely to misfold into native-like states

Our previous publications demonstrated that some misfolded entangled states are structurally similar to the native ensemble and can act as long-lived kinetic traps, persisting on timescales similar to the native state (1, 2). For this reason, such states can evade recognition by chaperones (3), which is part of the cell’s protein quality control machinery. We hypothesized that the same phenomenon may influence the targeting of proteins to the UPS. That is, some natively entangled proteins misfold and do not get ubiquitinated for proteasome degradation because they are structurally similar to the native state and hence are treated by E3 ligases similar to the native state. This hypothesis predicts that in the birth dating mass spectrum data that those proteins that were identified as not being ubiquitinated but contain a native entanglement are (*i*) equally likely to misfold as the young ubiquitinated proteins, and (*ii*) are more likely to misfold into conformational ensembles that more closely resemble the native state.

To test the first prediction, we randomly selected 10 NU-E proteins, size-matched to the YU-E proteins (Table 1), and simulated their refolding (see Methods). For example, the NU-E protein CCR4-NOT transcription complex subunit 7 (Uniprot ID: Q9UIV1) (Fig. 2E) populated 6 metastable states in *Q*-*G* space. The native basin is State S6 in Fig. 2E, centered at *Q* = 0.96 and *G*≈0. This protein has 4 misfolded states involving a change in entanglement. Metastable states for other NU-E proteins can be found in supplementary Figs. 22 to 31. We again quantified the probability of forming misfolded entanglement states (Eq. 4) and compared it with the misfolding probability of YU-E proteins. We find that the two groups do not exhibit statistically different misfolding propensities (Probability of misfolding = 0.35, 95% CI = [0.18, 0.52] versus 0.29, 95% CI = [0.17, 0.43]; p-value = 0.6467, two-sided hierarchical permutation test for 10^6^ times, see Fig. 2G). We conclude that entanglement misfolding is equally common amongst those natively entangled proteins that were ubiquitinated and those that were not. The same conclusion was obtained when the simulations were extended to 20 μs (Fig. S21A and B).

To test the second prediction, we quantified the propensity for near-native misfolding in both groups by measuring the average fraction of native contacts in misfolded entanglement states normalized by the native state average, *Q*_norm_ (Eq. 5; per-protein values are shown in Table S4). We find that the YU-E proteins misfolded with an average *Q*_norm_ of 0.80 (95% CI: [0.71, 0.88], hierarchical bootstrapping for 10^5^ times, see Methods), while the NU-E proteins misfolded with an average *Q*_norm_ of 0.93 (95% CI: [0.87, 0.97], hierarchical bootstrapping for 10^5^ times, see Methods). The difference between these is significant (p-value = 0.0248, two-sided hierarchical permutation test for 10^6^ times, see Methods). For example, the NU-E protein Q9UIV1 populated a near-native misfolded state S5 in Fig. 2F, centered at *Q* = 0.94 and *G* = 0.11. This state formed 94% of its native contacts, which is very close to the level of the native ensemble at 96% at 310 K. Thus, NU-E misfolded proteins more closely resemble the native state than YU-E proteins. The difference in *Q*_norm_ between YU-E and NU-E misfolded proteins tended to decrease when the simulations were extended to 20 μs, as some proteins progressively folded toward the native state. Nevertheless, the same overall trend persisted throughout the extended simulations (Fig. S21C and D).

In addition, we examined the ratio of the solvent-accessible surface area (rSASA, see Eq. 6; per-protein values are shown in Table S5) of hydrophobic residues in misfolded-state ensembles relative to the corresponding native-state ensembles and compared this metric between YU-E and NU-E misfolded proteins (see Methods). We found that misfolded states in YU-E proteins displayed greater exposure of hydrophobic surface than misfolded NU-E proteins (⟨ *rSASA* ⟩ = 1.45, 95% CI: [1.21, 1.70] vs. 1.13, 95% CI: [1.04, 1.25], p-value = 0.0378, two-sided hierarchical permutation test for 10^6^ times, see Methods). The more native-like hydrophobic surface area of misfolded NU-E proteins - consistent with their more near-native conformations - may contribute to their reduced recognition by E3 ubiquitin ligases.

These three results are consistent with our hypothesis, and indicate misfolding is occurring in the NU-E proteins to a similar degree as the YU-E proteins, but it seems likely they are not being recognized by E3 ubiquitin ligases and tagged for degradation.

### Co-translational folding on the ribosome exhibits similar misfolding behaviors

A criticism of our simulations is that temperature-quench refolding is not the same process as co-translational folding, which can occur in fibroblast cells. To probe whether the misfolded states are similar on- and off- the ribosome, and whether YU-E proteins misfold more frequently than NU-NE proteins during co-translational folding we performed protein synthesis simulations on the ribosome, followed by post-translational folding (see Methods). We selected two size-matched proteins - protein P16152 (from the YU-E group) and protein A6NDU8 (from the NU-NE group) - that showed the largest difference in misfolding probability in our temperature quench refolding simulations.

Consistent with our refolding simulations in the absence of the ribosome, protein P16152 exhibited higher misfolding probability than protein A6NDU8 on and off the ribosome. On the ribosome we identified one early misfolded state (state C8, see Fig. 3C) in protein P16152 (indicated by the non-zero G values found during synthesis, see Fig. 3E), while no misfolding was found in protein A6NDU8 (G value remains zero during synthesis, see Fig. 3F). Post-translationally, we identified 5 misfolded states in protein P16152 and again no misfolding was observed in protein A6NDU8 (see Fig. 3A and B). In protein P16152, the early misfolded state C8 involving loss of entanglement (failure-to-form) leads to two kinetically trapped, misfolded states (P2 and P8) during post-translational folding, resulting in only 8% of simulation trajectories reaching the native state P9 (see Fig. 3C). In contrast, protein A6NDU8, which did not misfold co-translationally, folded rapidly to its native state (P3) after synthesis in all simulation trajectories (see Fig. 3D).

**Figure 3.**
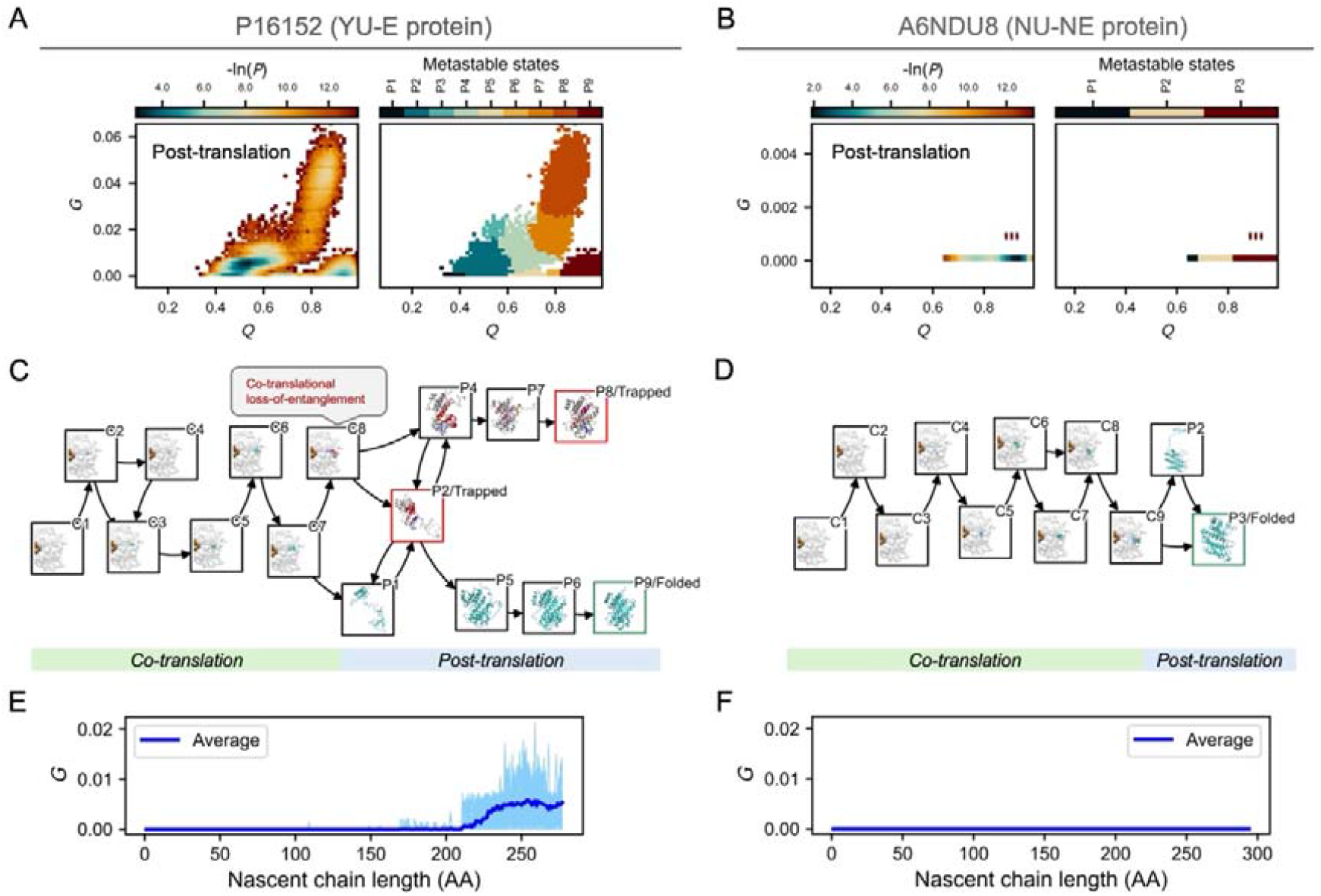
Similar misfolded states are populated by the nascent chain when co- and post-translational folding are modeled. A, B, The log probability surfaces (–ln P; left) in the Q–G space and the corresponding metastable states populated during the post-translational phase of folding simulations (right) for the YU–E protein P16152 (A) and the NU–NE protein A6NDU8 (B). The axes are identical to those in Fig. 2A and C. C, D, The most probable co- and post-translational folding pathways (top 85% probability) depicted as network diagrams for P16152 (C) and A6NDU8 (D), where nodes represent metastable states (labeled “C” for co-translational and “P” for post-translational), and edges denote transitions between states, with arrows indicating directionality. Representative structures for each metastable state are displayed on the corresponding nodes; co-translational states show the ribosome (white) and tRNAs (orange). Both nascent chains (co-translational) and full-length proteins (post-translational) are illustrated in the same manner as in Fig. 2B, D, and F. Nodes corresponding to the native state and kinetically trapped states (misfolded states involving entanglement changes where trajectories terminated) are outlined in green and red, respectively. E, F, The G values for each co-translational trajectory (light blue) and their trajectory-averaged values (dark blue) as a function of nascent-chain length.

In the temperature-quench refolding simulations we identify 5 misfolded states in protein P16152 and none in protein A6NDU8 (compare Fig. 2A and C). For protein P16152, 4 out of the 5 misfolded states are common between refolding and synthesis (compare Fig. 2A and Fig. 3A). Thus, the results from our synthesis simulations are qualitatively consistent with conclusions drawn from the temperature-quench simulations, indicating that YU-E proteins are inherently more prone to misfolding with entanglement changes and such misfolding events can occur during synthesis on the ribosome.

## Discussion

Our findings demonstrate that proteins containing native entanglements are nearly twice as likely to be ubiquitin-tagged for degradation during or shortly after synthesis and are four times more likely to misfold compared to similar proteins that lack native NCLEs. These results indicate that entanglement misfolding promotes ubiquitination leading to proteosomal degradation in a subset of proteins in human fibroblast cells. Another subset of proteins, however, contain native NCLEs that are likely misfold to a similar extent but are not ubiquitinated. This subset misfolds into conformational ensembles that more closely resemble the native state which presumably allows them to evade detection by E3 ligases to a similar degree as folded proteins. We therefore conclude that entanglement misfolding can lead to divergent fates for a nascent protein – proteins that misfold soon after synthesis into less structured states are degraded, but those that misfold into more ordered states persist, likely remaining soluble and nonfunctional inside cells (1–3) (Fig. 4).

**Figure 4.**
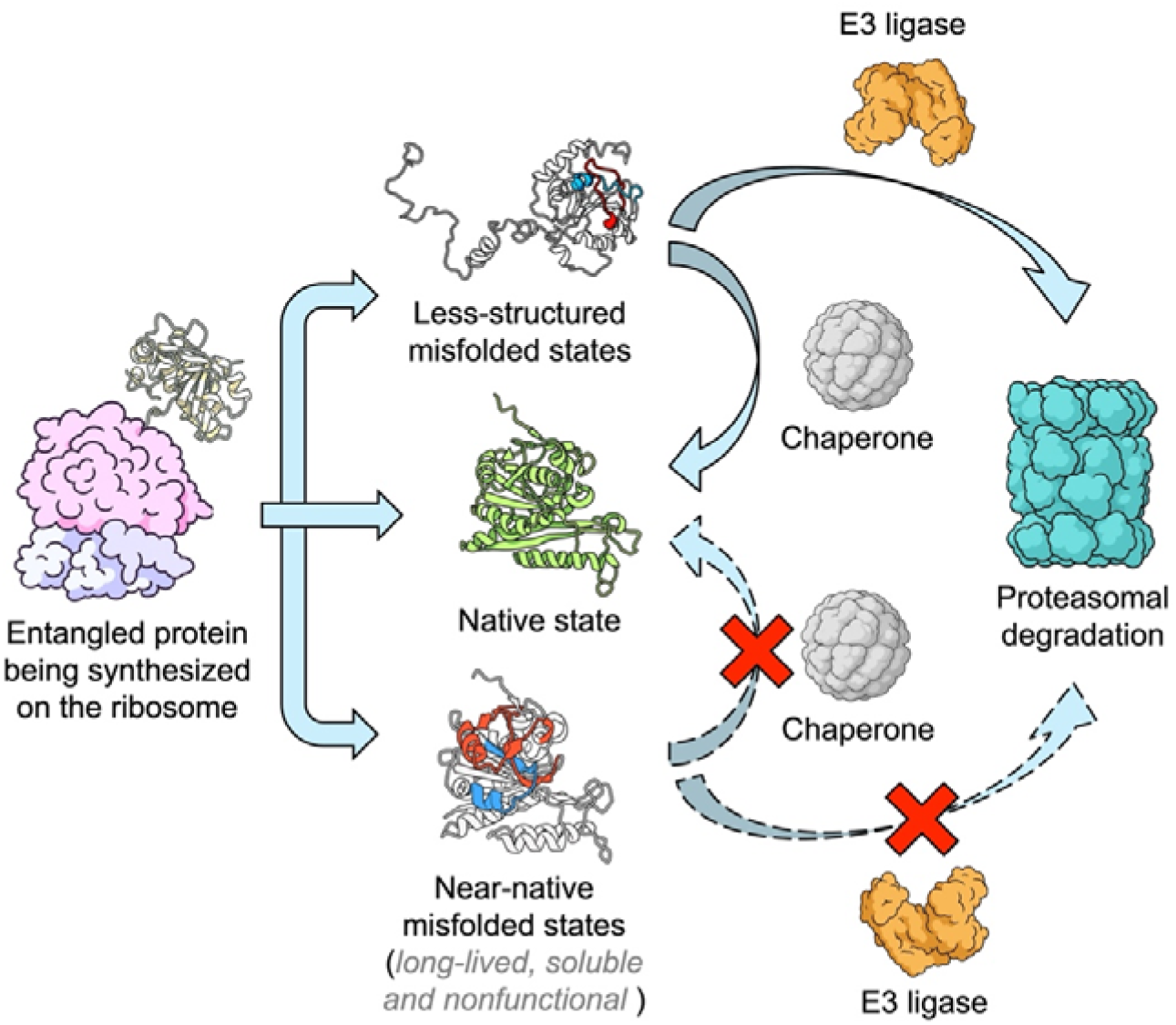
Entanglement misfolding leads to divergent protein fates in the cell. Proteins containing NCLEs can misfold during or after synthesis by the ribosome into either less structured misfolded states or near-native misfolded states. Misfolding is driven by changes in entanglement status, with non-native entanglements highlighted in red (loop) and blue (thread). Less structured misfolded states undergo substantial conformational changes and are readily recognized by chaperones for refolding or by E3 ligases for ubiquitination and subsequent proteasomal degradation. In contrast, near-native misfolded states can evade chaperone and E3 ligase surveillance, remaining soluble but partially or fully nonfunctional, thereby posing a challenge to maintaining functional protein homeostasis.

Our simulation results provide a plausible mechanism by which proteins containing native NCLEs are more likely to misfold than proteins that do not contain them. Misfolding involving changes of entanglement can occur via a gain of a non-native entanglement, or the failure to form a native entanglement. Proteins containing native NCLEs can misfold via both mechanisms, while proteins without them can only exhibit a gain of entanglement. Hence, proteins with native NCLEs are more likely to misfold and more likely to be ubiquitinated. This conclusion is consistent with a recent study that found proteins with native NCLEs are more likely to misfold than proteins without them (8).

From our analyses, we can estimate key numbers related to entanglement misfolding and degradation. 74% (1,135 out of 1,541) of the observable globular proteins in our dataset (excluding membrane proteins and IDPs) contain one or more native NCLEs. Of these entangled globular proteins, 49% (557 out of 1,135) get ubiquitinated for proteasome degradation (proteins with accumulated Kε-GG peptides upon proteasome inhibition) and 44% (497 out of 1,135) do not get ubiquitinated. The remaining 7% are tagged with ubiquitin but are not targeted for proteasome degradation. (These likely reflect non-degradative functions of ubiquitin, such as regulatory or signaling roles.) Using the observation from our simulations that 90% (9 out of 10) of the non-ubiquitinated, entangled proteins exhibit misfolding (i.e., proteins whose estimated misfolding probabilities have 95% CIs that do not overlap zero, see Fig. 2G), we estimate that 40% (= 44% × 90%) of natively entangled proteins misfold but do not get ubiquitinated. Similarly, using the observation that 80% (8 out of 10) of the ubiquitinated, entangled proteins exhibit misfolding (Fig. 2G), we estimate that 39% (= 49% × 80%) of natively entangled proteins misfold and get tagged for degradation. We therefore estimate that in vivo while approximately half (39%/(40% + 39%) × 100% = 49%) of misfolded entangled proteins get tagged for degradation, the other half of misfolded entangled proteins do not. Thus, although native NCLEs significantly increase the chances of proteins to be ubiquitinated in comparison to non-entangled proteins, we estimate that nearly half of misfolded entangled proteins nonetheless manage to evade UPS-mediated degradation. These results are consistent with an earlier prediction that entanglement misfolding was likely widespread and that a large proportion of proteins would exhibit near-native like misfolded states that could bypass degradation, aggregation, and chaperone refolding activities (1). According to these results and earlier reports (1–3), the protein homeostasis model should be updated to include an additional state that many proteins appear to populate: widespread entanglement-misfolded states that are soluble and less-functional or even non-functional, and persist inside cells for long time periods as they are not successfully acted upon by degradation and chaperone pathways (Fig. 4). The biological significance of soluble, less-functional misfolded states, we speculate, could have a variety of detrimental effects for an organism over its lifetime. The accumulation of these states will almost certainly decrease the efficiency of subcellular process since these proteins have lost their native function. Therefore, we hypothesize they could contribute to aging – which is the progressive and systematic decline of function across all scales, from the molecular to an entire organism (29). Similarly, these misfolded states could be the origin of many loss-of-function diseases whose causes are unknown (30–33).

Ubiquitination does not just arise from misfolding, it can also arise from several non-mutually exclusive mechanisms. Previous studies have shown that newly synthesized proteins may be degraded due to failure to assemble with binding partners (26) (known as orphan proteins), while ribosome pausing or collisions can trigger ribosome-associated quality control pathways (34), including disome-associated surveillance mechanisms (35). These processes can expose hydrophobic regions or degrons that are subsequently recognized by molecular chaperones and E3 ligases. Our findings suggest that misfolding driven by NCLEs represents an additional, complementary mechanism that contributes to early ubiquitination, rather than being mutually exclusive with these established pathways. When we controlled for oligomeric status, the association between entanglement and early ubiquitination was unaltered, indicating the findings in this study are not driven by oligomerization-dependent instability.

NCLEs likely reflect an evolutionary trade-off between structural and functional advantages, on the one hand, and the ability to reliably fold on the other. Our previous studies demonstrated that NCLEs are widespread in the native structure of proteins and are enriched in proteins with catalytic and small-molecule/ion-binding functions, suggesting that these geometric features help support structural architectures required for biochemical activity (10, 36). At the same time, NCLEs introduce kinetic constraints during folding, making proteins more prone to misfolding due to premature loop closure (8). Our previous work further suggests that cellular proteostasis mechanisms can partially mitigate this cost (8). Although near-native misfolded states often evade both chaperone refolding and degradation pathways and can therefore persist in cells, the misfolding of essential proteins, which maintain core cellular functions, are more effectively avoided by chaperone systems – at least in *E. coli* (8). Together, these findings suggest that one reason NCLE-containing proteins may persist through evolution is because their functional advantages can outweigh their misfolding risks, particularly when critical proteins receive greater protection from proteostasis mechanisms.

Like all models, our simulations have certain limitations. The simulation framework used in this study is designed to model the folding of globular monomeric proteins and therefore does not explicitly represent transmembrane proteins or proteins that are stabilized as subunits within larger complexes present in the Ubq-MS dataset. Nevertheless, it seems plausible that similar misfolded entanglement states may also occur in these protein classes, as they do contain NCLEs (e.g., one-fifth transmembrane protein domains contain NCLEs (37)) that could potentially misfold through mechanisms analogous to those observed in globular proteins (5). Further, protein folding in vivo occurs in the presence of molecular chaperones such as the Hsp70/Hsp90 systems and the TRiC/CCT chaperonin, whereas these chaperones are absent from our simulations. And our simulations do not model differences in substrate recognition by the diverse range of E3 ubiquitin ligases. Taken together, what this suggests is that while the statistical associations we observe in this study are likely arising from entanglement misfolding the effect size, or magnitude, of misfolding is likely smaller in the presence of these other cellular factors but not abolished entirely.

We selected the proteins for simulation based on the ubiquitination data from Ref. (19). To the best of our knowledge, this dataset is unique in that it simultaneously profiles the identity, proteasomal flux, and molecular age of ubiquitinated proteins, enabling identification of the subset of the proteome that is rapidly targeted for proteasomal degradation soon after synthesis. The overall accuracy of the measurements was verified by a battery of control experiments. Nonetheless, several limitations warrant consideration. Because nascent ubiquitinated proteins are inherently transient, their detection required proteasome inhibition, raising the possibility that MG132-induced proteotoxic stress may itself elevate ubiquitination of newly synthesized proteins, complicating the distinction between constitutive ubiquitination and stress-induced ubiquitination. In addition, these experiments were performed in a single quiescent human fibroblast cell line, and whether the observed properties of the ubiquitinome extend to other cell types remains to be determined.

The conclusions of this study are restricted to globular proteins, as our structural analysis necessitated focusing on high-confidence AlphaFold models (average pLDDT ≥ 85), which represent ∼35% of the proteins detected in the Ubq-MS dataset. The excluded proteins are overrepresented in classes such as transcription factors, DNA-binding proteins, zinc-finger proteins, and coiled-coil proteins - categories that often include long intrinsically disordered segments or are conformationally flexible and therefore less amenable to high-confidence structure prediction. IDPs and regulatory proteins, such as transcription factors, have been found to populate NCLE states (38). So it may be possible these states are also relevant to degradation in these systems.

Our study implies but does not establish a causal role of NCLEs in driving young-age ubiquitination. To test this mechanistic link several experimental approaches could be used. Limited proteolysis mass spectrometry (LiP–MS) (39) performed under proteasome inhibition conditions could reveal proteome-wide conformational changes associated with the accumulation of misfolded states, enabling direct comparison between structural perturbations and ubiquitination profiles. Structural methods such as cryo-electron microscopy (cryo-EM) (40) may enable direct characterization of misfolded ensembles involving entanglement changes for carefully selected targets, although such experiments will require proteins with sufficiently stable and less heterogeneous conformational states.

Moreover, it would be interesting, in future studies, to explore how broad these conclusions apply to various cell types and organisms. And testing other downstream biological consequences of entanglement misfolding such as potential associations with proteins that are known to aggregate, and its contribution to dysregulation in aging and disease.

## Materials and Methods

### Identification of young-ubiquitinated proteins using human proteome birthdating data

Human proteome birthdating data were obtained from a previous study by Meadow et al. (19) that quantified the age distribution of ubiquitinated proteins based on trypsin-digested peptides containing di-glycine residues on lysine side chains (Kε-GG peptides) in the presence and absence of proteasome inhibition. In the same study, the baseline age distribution of the proteome was also established by profiling unmodified peptides (i.e., trypsin-digested peptides lacking the Kε-GG motif) under conditions without proteasome inhibition. In total 6,450 proteins were detected and quantified by this experiment.

Proteins were classified as ubiquitinated if at least one Kε-GG peptide mapped to that protein was identified, and as non-ubiquitinated if no such peptides were detected. Upon proteasome inhibition, ubiquitinated proteins targeted by the UPS accumulate, resulting in increased abundance of their Kε-GG peptides compared to those produced without inhibition. Conversely, Kε-GG peptides that do not increase upon proteasome inhibition likely originate from ubiquitinated proteins that are not acted on by the proteasome (19). To focus specifically on UPS-mediated degradation, we calculated the maximum fold change in Kε-GG peptide abundance (with vs. without proteasome inhibition) for each protein and excluded proteins whose maximum fold change did not exceed 6.75—a threshold that approximately separates the bimodal distribution into two distinct subpopulations (see Fig. S32).

To identify YU proteins, we calculated the fold change between the age of the youngest Kε-GG peptide and the baseline protein age (defined as the median age of unmodified peptides for each protein). Ubiquitinated proteins were classified as YU only if this fold change was less than 0.2772, a threshold that also separates the age distribution into two subpopulations (see Fig. S33). Additionally, to ensure these proteins were recently synthesized, we further filtered for proteins with median Kε-GG peptide ages younger than 6 hours. In total 906 proteins were identified to be YU proteins.

As a control set, we selected all NU proteins that had no identified Kε-GG peptides (n = 3,213).

### Identification of entangled proteins in AlphaFold-predicted structures

To identify non-covalent lasso entanglements, we first detect all loops that are pierced by either the N-terminal or C-terminal segment of the protein, such that at least one crossing occurs. Candidate loops are initially identified based on potential crossing events using Gauss linking numbers (8, 10), and these are subsequently refined through minimal surface analysis implemented in Topoly (41). A loop is defined as a protein segment closed by a native contact, where two residues have any heavy atoms within 4.5 Å of each other. We exclude covalent lassos where the loop-closing residues are cysteines (42, 43), as well as loops known or predicted to form knots or slipknots (44–46).

AlphaFold-predicted structures of the human proteome (n = 20,588) were obtained from the AlphaFold F1 model database at version 4 (20, 21). The proteins with covalent lassos, knots or slipknots identified were first excluded. Non-covalent lassos were further filtered by excluding those in which the loop-closing native contact had a pLDDT score (20) < 70. Additionally, if any crossing residue had a pLDDT score < 70, that crossing and all subsequent crossings within the same terminus were discarded. To minimize false positives and exclude intrinsically disordered proteins (IDPs), only structures with an average predicted local distance difference test (pLDDT) score ≥ 85 were included in the analysis (n = 5,641).

Proteins were classified as *entangled* if they contained at least one non-covalent lasso entanglement (n = 3,776). Proteins lacking such entanglements were classified as *non-entangled* (n = 1,865).

### Logistic regression analysis

We used logistic regression models (47) to test associations between predictors (e.g., the presence of NCLEs in a protein) and outcomes (e.g., young-age ubiquitination), while controlling for potential confounding factors. The general form of the logistic regression model is:

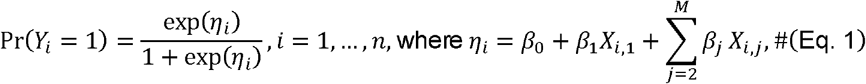

where *Y*_*i*_ is the binary outcome for protein (or residue) *i, x*_*i*,1,_ is the predictor of interest, *x*_*i,j* >1_ are the *M* − 1 confounding factors, and *β*_*k*>O_ are the regression coefficients representing the log-odds ratios of the predictors.

To test the association between young-age ubiquitination and the presence of NCLEs, *Y*_*i*_ was set to 1 if the protein is a YU protein and 0 if it is a NU protein. x_*i*,1_, was defined as a binary indicator of NCLE presence (set to 1 if it has NCLEs otherwise 0), and the confounding factor x_*i*,2_was the standardized protein length. For each predictor, the O.R. was calculated as exp(*β*_*k*>0_), and 95% confidence intervals (CIs) for the O.R. were estimated using the Wald method. All logistic regressions were performed using the Python statsmodels package (48).

### Processing other independent human ubiquitination datasets

We curated three additional human ubiquitination datasets from two independent publications, each reporting changes in Kε-GG peptide abundance following proteasome inhibition. One dataset was obtained from the study by Wagner et al. (27), whereas two additional datasets were obtained from Kim et al. (28), in which different proteasome inhibitors were used (Kim_btz and Kim_epox datasets). For each dataset, we identified a fold-change threshold corresponding to the separation between the two major modes of the observed distribution. Kε-GG peptides with fold changes above this threshold were classified as enriched upon proteasome inhibition. Proteins containing at least one enriched Kε-GG peptide were classified as ubiquitinated proteins (i.e., candidate substrates targeted for proteasomal degradation), whereas proteins with at least one quantified Kε-GG peptide but no enriched peptides were classified as non-ubiquitinated proteins. Proteins for which all peptides failed the original study quality-control criteria and therefore lacked quantitative fold-change values (reported as NaN) were excluded from the analysis. Within the ubiquitinated and non-ubiquitinated protein sets, we identified NCLEs and estimated O.R.s and 95% CIs using the same approaches described above.

In the Kim_btz and Kim_epox datasets, protein identities were reported as IPI accession numbers rather than UniProt identifiers. We therefore evaluated two strategies for mapping IPI entries to UniProt IDs. The first approach used the official IPI cross-reference mapping database. The second approach inferred UniProt IDs using annotated gene names together with the ubiquitination-site motif sequences. The first approach recovered 16% fewer UniProt proteins than the second approach and resulted in statistically less significant associations. Therefore, we used the second approach in the Results Section, save for where we mention using an ‘alternative’ IPI-to-UniProt, which used the first approach.

### Selection of candidate proteins for temperature quench simulations

Because our CG model is not designed to simulate membrane protein folding, we first excluded from the dataset the transmembrane proteins (i.e., UniProt (22) entries annotated with transmembrane domains). We also excluded proteins that form large complexes, as their folding processes may depend on interactions with partner proteins, which are not feasible to model. These proteins typically adopt elongated shapes. To remove such cases, we calculated asphericity for each protein using the same set of AlphaFold structures and the HullRad script (49), and excluded all proteins with asphericity values > 0.1.

After these filters, the candidate pool contained 94 YU-E proteins, 33 NU-NE proteins, and 172 NU-E proteins. From the YU&E group, we randomly selected 10 proteins. To control for protein length, we also selected 10 proteins from each of the other groups using a size-matching procedure. Specifically, for a YU-E protein of length *L*, we allowed selection of NU-NE or NU-E proteins with lengths within [*L* - *L*_*b*_,*L* + *L*_*b*_ ], where *L*_*b*_ = 10 residues. Selected proteins were then removed from the candidate list to avoid duplication. If no match was available within this range, we iteratively expanded the buffer (2×, 3×, etc.) until a protein was selected.

Finally, we visually inspected all selected proteins to confirm a globular fold. Any elongated proteins that passed the automated filters were manually excluded and replaced by a randomly selected alternative protein. The selected proteins for simulation are listed in Table 1.

### Temperature quenching simulations

To simulate the protein folding process, we performed temperature-quenching simulations using a Gō-based coarse-grained (CG) model (1, 2, 50–55). In this model, each amino acid residue is represented as a single interaction site centered on the Cα atom, and interactions are governed by a structure-based potential energy function (2). The force field parameters were tuned to reproduce the structural stability of each protein (2). Specifically, structural stability was iteratively evaluated by running ten parallel 1-μs MD simulations starting from the AlphaFold structures at 310 K, with different parameter sets obtained from prior work (2). The minimum parameter values capable of maintaining ≥68% of native contacts for more than 98% of the simulation time were selected for protein parameterization (2). The final force field parameters used for each protein are listed in Tables S6-S8.

For the temperature-quenching simulations, AF2 structures were first thermally unfolded at 800 K for 60 ns, followed by refolding at 310 K for 2 μs. For each protein, we performed 50 independent simulations with different random seeds. All simulations were carried out using Langevin dynamics with a collision frequency of 0.05 ps^−1^ and a time step of 15 fs, implemented in OpenMM (56).

### Metastable states clustering

To characterize misfolded states associated with changes in entanglement, we used two previously established order parameters, *Q* and *G*, to describe the folding status of proteins (2).

The parameter *Q*, representing the fraction of native contacts, was calculated as:

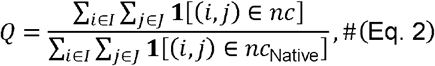

where *i* and *j* are the residue indices and satisfy j >*i*+ 3; *I* and *J* are both the set of residues within secondary structure elements (α-helical or β-strands); 1 [·] is the indicator function which returns 1 when condition is satisfied and 0 otherwise; *nc* and *nc*_*Native*_ are the set of residue pairs forming native contacts in the structure being evaluated and the native structure, respectively. Native contacts are considered formed when the distance between the Cα atoms of residues *i* and *j* does not exceed 1.2 times their native distance and the native distance does not exceed 8 Å.

The parameter *G* quantified deviations in entanglement relative to the native structure:

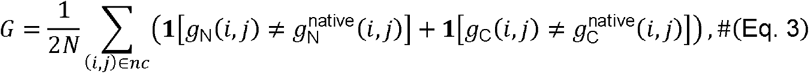

where (*i, j*) is a pair of residues forming a native contact (*nc*) in the structure being evaluated; *g*(*i,j*) and *g*^native^ (*i,j*) represent the linking numbers of contact (*i, j*) in the testing and native structures, respectively, estimated using a discrete version of Gauss double integration (2). The integral values were rounded based on a 0.6 threshold (e.g., 0.5 → 0, -1.7 → -2) (8). The subscripts N and C refer to the N- and C-terminal threads, respectively. *N* is the total number of native contacts in the native structure.

*Q* and *G* were computed for each frame of the refolding trajectories. To avoid artifacts of the coarse-grained force field, we also examined secondary-structure packing chirality (6) and removed trajectories that populated mirror-image states (6, 57, 58). Trajectory data for each protein were projected onto the *Q*–*G* space, and conformations were grouped using the *k*-means algorithm (59) into 100–400 microstates, depending on the complexity of the free-energy landscape. A Markov state model (MSM) was then constructed, and microstates were coarse-grained into 4–15 metastable states using the PCCA+ algorithm (60). All clustering and MSM construction were performed using the PyEMMA package (61).

### Estimation of protein misfolding propensity

The misfolding propensity of a protein 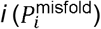 was estimated as the fraction of misfolded frames (obtained from the MSM) found in the final 100 ns of simulation trajectories, averaged across all independent trajectories. It can be computed as:

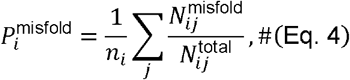

where 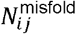 and 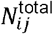 are the number of misfolded frames and total number of frames in the final 100 ns of simulation trajectory *j*, respectively. *n*_*i*_is the number of trajectories of protein *i*. Two categories of misfolded states were considered: non-native states involving entanglement changes, defined as states with *G* at the cluster center ≥ 0.005, and non-native states without entanglement changes, defined as states with *G* at the cluster center < 0.005.

For each protein, 95% confidence intervals (CIs) of misfolding propensities were estimated by bootstrapping per-trajectory data 10^5^ times. An overall misfolding propensity was then calculated for the YU-E and NU-NE protein groups by averaging the per-protein data, respectively.

### Calculation of *Q*_norm_

To evaluate how closely misfolded states resemble their native conformations, we computed a normalized fraction of native contacts, denoted as *Q*_norm_, by scaling the fraction of native contacts in misfolded states relative to the average native-state value. For a trajectory *j* of protein *i*, we compute *Q*_*norm*_ (*i,j*) as:

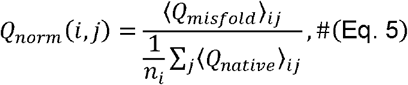

where ⟨ *Q*_*native*_ ⟩_*ij*_ and ⟨ *Q*_*misfold*_ ⟩_*ij*_ represent the average fraction of native contacts for the native and misfolded entanglement states (*G* at the cluster center ≥ 0.005), respectively, computed over the final 100-ns frames in trajectory *j* of protein *i*, and *n*_*i*_ is the number of trajectories of protein *I* that contained native state in the final 100-ns frames.

For each protein, per-trajectory values were averaged to yield ⟨ *Q*_*norm*_ ⟩_*i*_. The 95% CI was estimated by bootstrapping both ⟨ *Q*_*native*_ ⟩_*ij*_ and ⟨ *Q*_*misfold*_ ⟩_*ij*_ values 10^5^ times, recalculating ⟨ *Q*_*norm*_ ⟩_*i*_at each iteration to generate the empirical distribution. Group-level averages ⟨ *Q*_*norm*_ ⟩ were then calculated for the YU-E and NU-NE protein groups.

### Calculation of hydrophobic SASA ratio

In addition to *Q*_norm_, we computed hydrophobic SASA ratio (rSASA) of misfolded states with relevant to the native state. For a trajectory *j* of protein *i*, we compute *rSASA*(*i,j*) as:

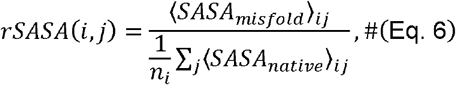

where ⟨ *SASA*_*native*_ ⟩_*ij*_ and ⟨ *SASA*_*misfold*_ ⟩_*ij*_ represent the average hydrophobic SASA for the native and misfolded entanglement states (*G* at the cluster center ≥ 0.005), respectively, computed over the final 100-ns frames in trajectory *j* of protein *i*, and *n*_*i*_ is the number of trajectories of protein *i* that contained native state in the final 100-ns frames. The hydrophobic SASA for a protein conformation was computed by summing over the SASA of hydrophobic residues (i.e., Ile, Val, Leu, Phe, Cys, Met, Ala, Gly and Trp) estimated using the freesasa package (62), following by back-mapping the CG structure to all-atom.

For each protein, per-trajectory values were averaged to yield *rSASA*_*i*_ . The 95% CI was estimated by bootstrapping both ⟨ *SASA*_*native*_ ⟩_*ij*_ and ⟨ *SASA*_*misfold*_ ⟩_*ij*_ values 10^5^ times, recalculating *rSASA*_*i*_ at each iteration to generate the empirical distribution. Group-level averages ⟨ *rSASA* ⟩were then calculated for the YU-E and NU-NE protein groups.

### Hierarchical resampling for group-level 95% CIs and permutation test

The misfolding propensity and *Q*_norm_ values were first computed per simulation trajectory, then averaged per protein and grouped into the YU–E and NU–NE categories. To properly account for intra-protein variability (i.e., deviations among independent trajectories), we applied hierarchical resampling (63, 64) for both bootstrapping and permutation tests when estimating group-level statistics.

For hierarchical bootstrapping, protein labels within each group were resampled with replacement to generate a new list of proteins. For each resampled protein, trajectory-level data were again resampled with replacement, followed by recalculation of cluster and group means. This procedure was repeated 10^5^ times to generate the empirical distribution of group means. The 95% CIs were determined from the 2.5th and 97.5th percentiles of this distribution.

For the hierarchical permutation test, the observed test statistic was defined as the absolute difference between group means. We first randomly resampled the trajectory means for each protein in both groups with replacement for 1,000 times and computed the new cluster means, producing 1,000 new lists of cluster means for each group. Next, for each new list, we combined the two groups, permuted the protein labels for another 1,000 times, divided proteins into two groups and recomputed the test statistic. This generated in total 10^6^ samples for the null distribution. The two-tailed p-value was computed as the fraction of null samples equal to or greater than the observed test statistic.

### Simulation of co-translational folding of nascent proteins

Co-translational folding of nascent protein chains was simulated using our previously established continuous synthesis protocol (2). Briefly, translation on the ribosome was modeled as a three-step process for each elongation cycle: (1) aminoacyl-tRNA binding to the A-site, (2) peptidyl transfer from the P-site to the A-site extending the nascent chain by one amino acid, and (3) translocation of the A-site nascent chain to the P-site. The elongation rate was set to an average of 4.2 codons/s, representing the mean of experimentally measured rates (3.5–4.9 codons/s) in human cells (65). The dwell time values of the A-site tRNA binding (0.0017 s) and peptidyl transfer (0.021 s) were obtained by scaling the *E. coli* values (2) by the ratio of codon elongation rates (20/4.2 ≈ 4.8). For each elongation step, the simulation time was sampled from an exponential distribution with the corresponding mean dwell time, then rescaled by a factor of 4,331,293 (2). At the stop codon, the full-length nascent chain nascent chain was released from the exit tunnel and dissociated from the ribosome surface. The resulting structure was used as the starting conformation for post-translational folding simulations.

The CG structure and force field of the human ribosome were generated following our previous protocol (2), using the high-resolution cryo-EM structure PDB 8G61 as the starting model. The 60S subunit, as well as A- and P-site tRNAs, was coarse-grained using a three-/four-point RNA model (50) for ribosomal RNA and a Cα model (50) for ribosomal proteins. The model was truncated to retain only interaction sites within 30 Å of the exit tunnel centerline, 20 Å of the peptidyl transferase center (A4548 in 28S rRNA), and regions near the tunnel exit on the ribosome surface, resulting in 5,706 interaction sites in total.

Ribosomal interaction sites were fixed during the simulations, and their interactions with the nascent chain included only excluding volume and electrostatic forces (2). The van der Waals radii were taken from the *E. coli* ribosome model (2).

### Folding pathway analysis

To analyze co-translational folding pathways, protein synthesis simulation trajectories were divided into three segments based on nascent chain length (2): residues 1–100, 101–200, and 201 to full length. Within each segment, conformations were clustered into 200 groups using the *k*-means algorithm (59) based on the order parameters *Q* and *G*. Up to three metastable states were then identified in each segment using the PCCA+ algorithm (60). The resulting discrete co-translational trajectories were constructed from the sequence of metastable states across all segments. Post-translational metastable states were identified in the same manner as in the temperature-quench simulations. Combined discrete trajectories representing both co- and post-translational folding were generated by appending the post-translational discrete trajectories to the corresponding co-translational ones.

Folding pathways were extracted as follows (2): (1) record the starting state of the first frame; (2) traverse the trajectory, adding each newly visited state to the pathway; if a state is revisited, truncate the pathway at its first occurrence and continue forward; (3) repeat until the trajectory ends. This procedure yields loop-free pathways that include only on-pathway metastable states for each discrete trajectory. Distinct pathways and their distribution probabilities were then computed across all trajectories.

## Acknowledgments

Y.J. and E.P.O. thank Prof. Antonio Trovato for valuable discussions. E.P.O. gratefully acknowledges support from the National Institutes of Health (R35-GM124818) and this work was supported by the National Science Foundation (NSF) through the NSF National Synthesis Center for Emergence in the Molecular and Cellular Sciences (NCEMS) under Grant NSF MCB-2335029. Computational support for simulations and data analyses was provided by Penn State Institute for Computational and Data Sciences (RRID:SCR_025154)

## Author Contributions

Conceptualization: E.P.O.

Methodology: Y.J. and E.P.O.

Investigation: Y.J., A.J., S.G., and E.P.O.

Resources: Y.J., S.G., and E.P.O.

Data curation: Y.J., A.J., S.G., and E.P.O.

Validation: Y.J., S.G., and E.P.O.

Formal analysis: Y.J. and E.P.O.

Software: Y.J.

Visualization: Y.J.

Supervision: E.P.O.

Writing—original draft: Y.J. and E.P.O.

Writing—review and editing: Y.J., A.J., S.G., and E.P.O.

Funding acquisition: E.P.O.

Project administration: Y.J., S.G., and E.P.O.

## Data Availability

Source data for Figs. 1 to 3 and Fig. S21 are provided. All data needed to evaluate the conclusions in the paper are present in the paper and/or the Supplementary Materials. All simulation trajectories are publicly available on CyVerse at https://data.cyverse.org/dav-anon/iplant/projects/NCEMS/working-groups/protein-misfolding-aging/data/Protein-entanglement-misfolding-determines-divergent-fates/. The datasets used in the logistic regression, including the NCLE dataset of the human proteome, are available in the Github repository https://github.com/obrien-lab/Protein-entanglement-misfolding-determines-divergent-fates.

## Code Availability

Codes used for running the simulations and analyzing the trajectories are available in the Github repository https://github.com/obrien-lab/cg_simtk_protein_folding/. Codes used for data analyses and input files used for both simulations and analyses are available in the Github repository https://github.com/obrien-lab/Protein-entanglement-misfolding-determines-divergent-fates.

