## Supporting Information for "Protein entanglement misfolding determines divergent fates: proteasomal degradation or persistence in near-native misfolded states"

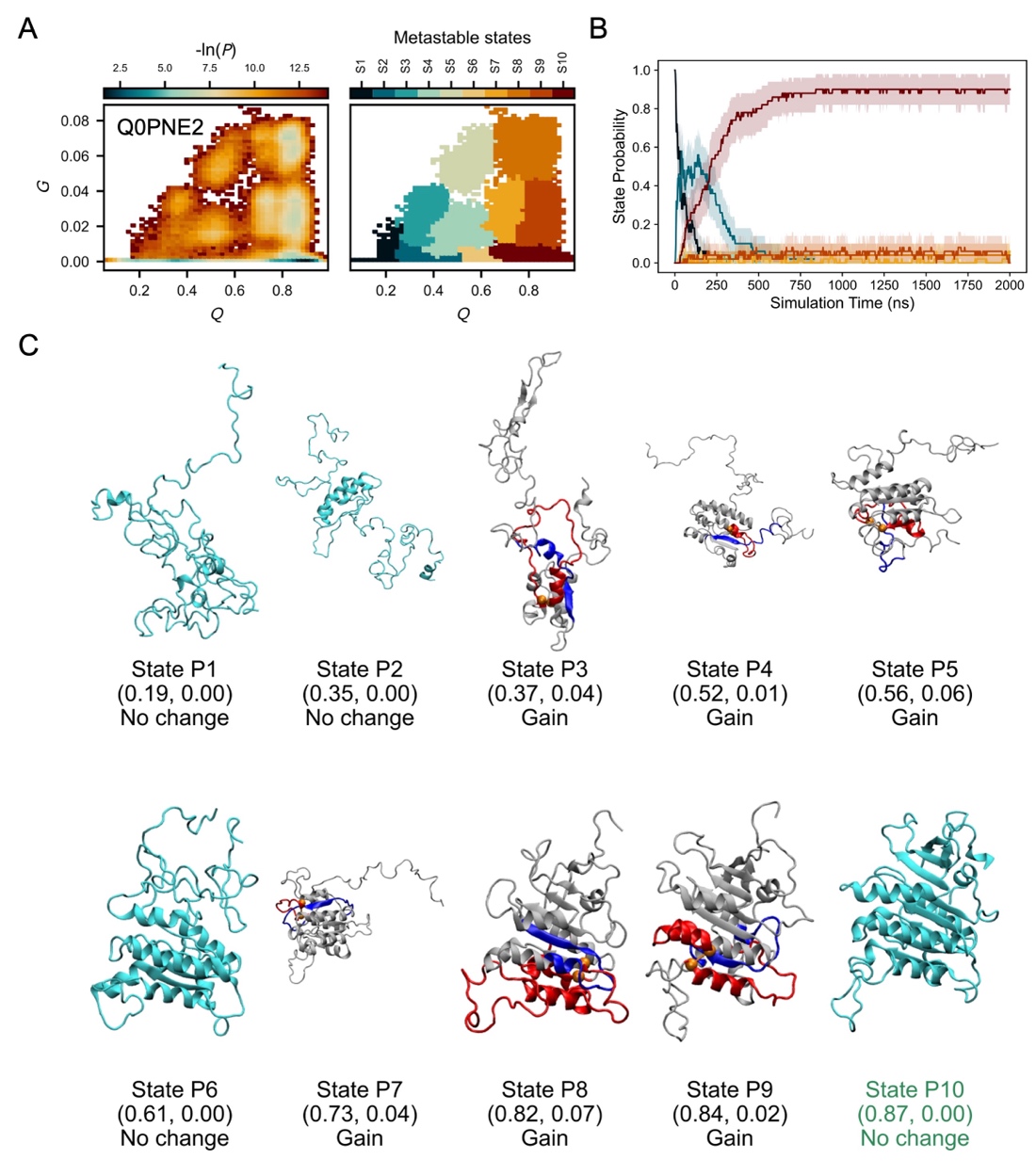

**Figure S1. Simulation structural ensemble of the YU-E protein Q0PNE2.** A. The probability distribution (-ln*P*) in the *Q* – *G* space (left) and the corresponding metastable states mapped onto structures from temperature-quench refolding simulations (right). B. Time evolution of metastable-state probabilities. Colors match the state assignments shown in panel A. Shaded regions denote 95% confidence intervals, estimated using bootstrap resampling (10^5^ iterations). C. Representative structures of each metastable state predicted from simulations. *Q* and *G* values at the cluster center, as well as the type of entanglement change (gain, loss or no change), are shown below each structure image. Color scheme follows that used in Fig. 2B, 2D, and 2F.

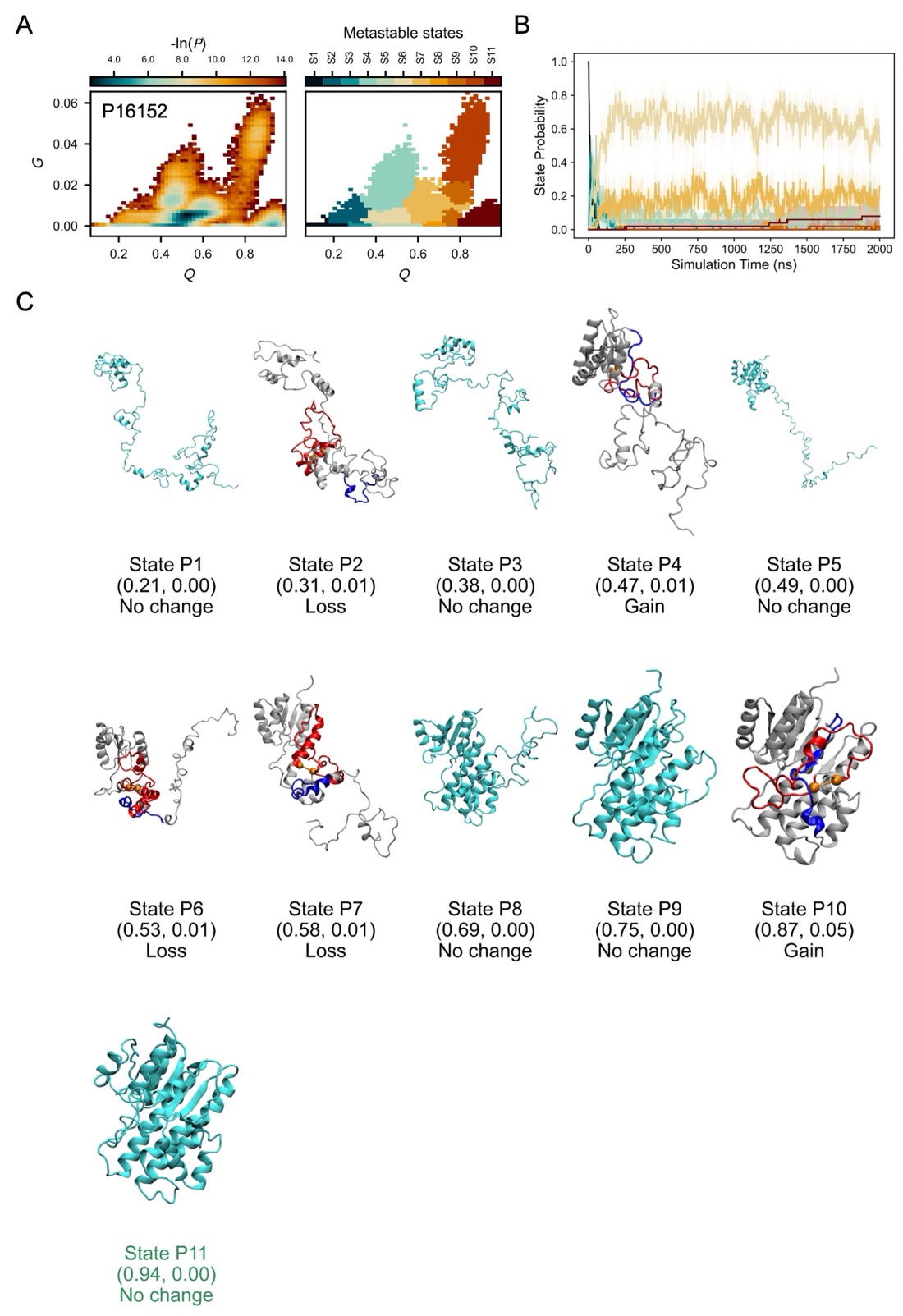

**Figure S2. Simulation structural ensemble of the YU-E protein P16152.** A. The probability distribution (-ln*P*) in the *Q* – *G* space (left) and the corresponding metastable states mapped onto structures from temperature-quench refolding simulations (right). B. Time evolution of metastable-state probabilities. Colors match the state assignments shown in panel A. Shaded regions denote 95% confidence intervals, estimated using bootstrap resampling (10^5^ iterations). C. Representative structures of each metastable state predicted from simulations. *Q* and *G* values at the cluster center, as well as the type of entanglement change (gain, loss or no change), are shown below each structure image. Color scheme follows that used in Fig. 2B, 2D, and 2F.

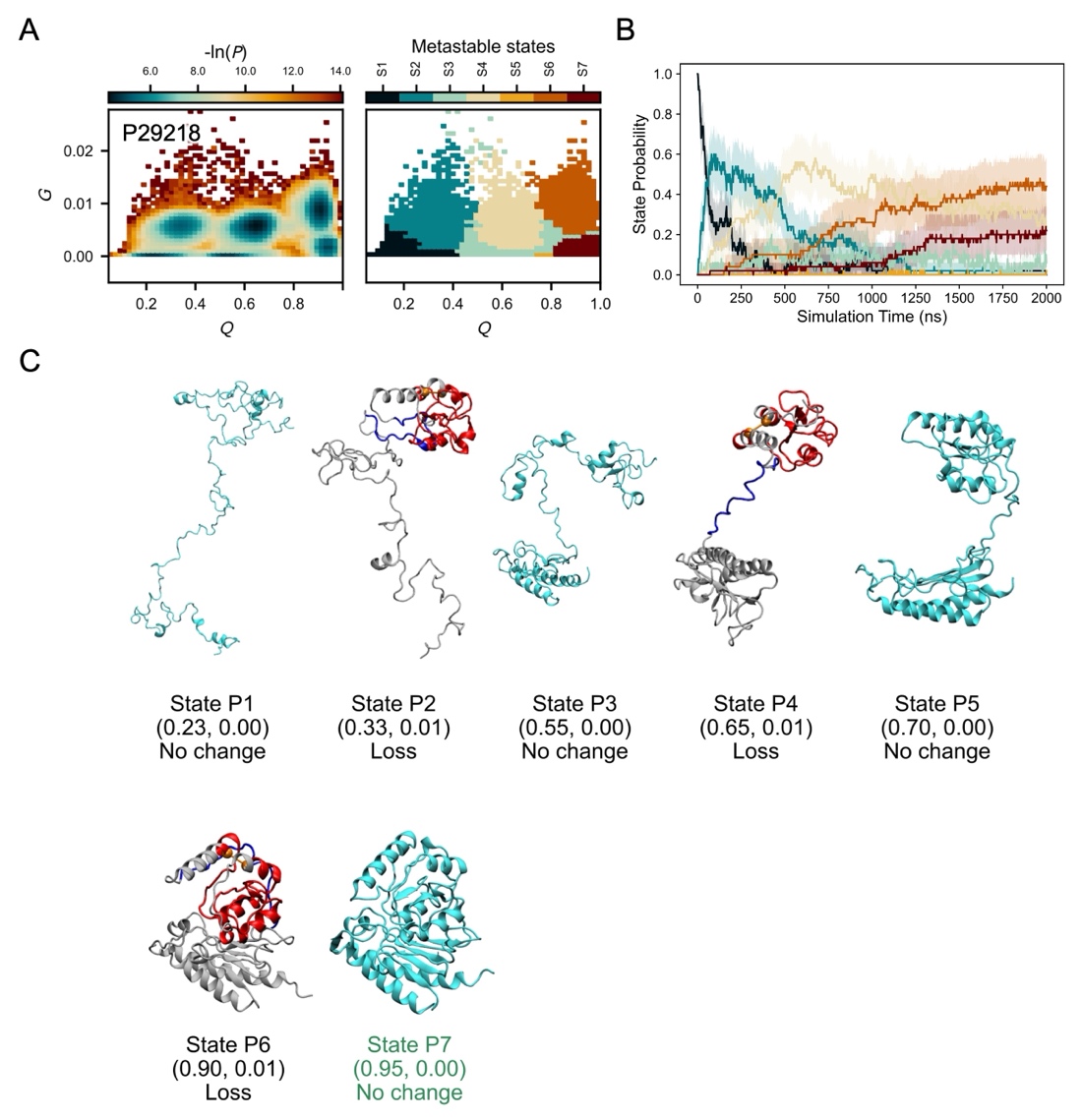

**Figure S3. Simulation structural ensemble of the YU-E protein P29218.** A. The probability distribution (-ln*P*) in the *Q* – *G* space (left) and the corresponding metastable states mapped onto structures from temperature-quench refolding simulations (right). B. Time evolution of metastable-state probabilities. Colors match the state assignments shown in panel A. Shaded regions denote 95% confidence intervals, estimated using bootstrap resampling (10^5^ iterations). C. Representative structures of each metastable state predicted from simulations. *Q* and *G* values at the cluster center, as well as the type of entanglement change (gain, loss or no change), are shown below each structure image. Color scheme follows that used in Fig. 2B, 2D, and 2F.

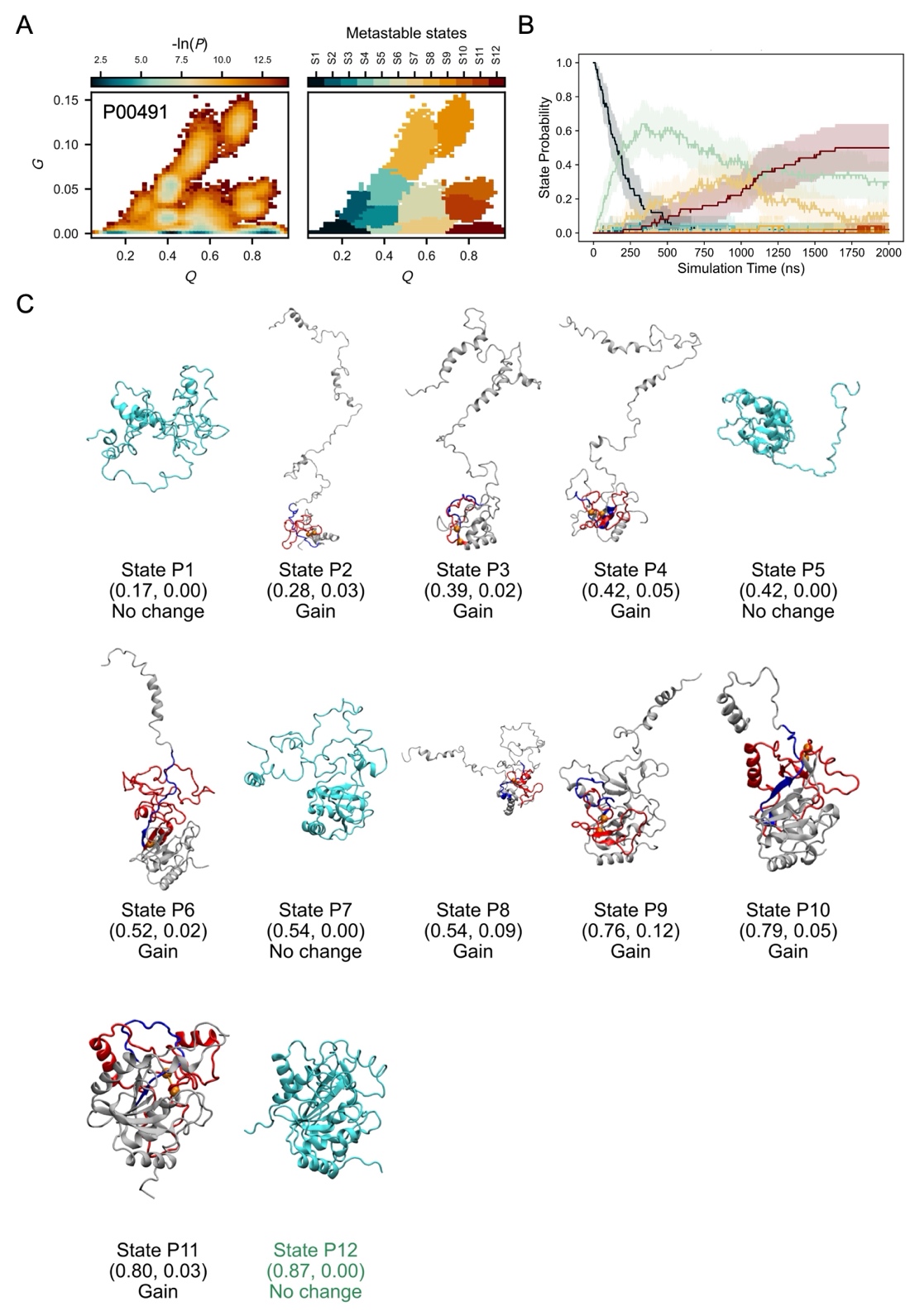

**Figure S4. Simulation structural ensemble of the YU-E protein P00491.** A. The probability distribution (-ln*P*) in the *Q* – *G* space (left) and the corresponding metastable states mapped onto structures from temperature-quench refolding simulations (right). B. Time evolution of metastable-state probabilities. Colors match the state assignments shown in panel A. Shaded regions denote 95% confidence intervals, estimated using bootstrap resampling (10^5^ iterations). C. Representative structures of each metastable state predicted from simulations. *Q* and *G* values at the cluster center, as well as the type of entanglement change (gain, loss or no change), are shown below each structure image. Color scheme follows that used in Fig. 2B, 2D, and 2F.

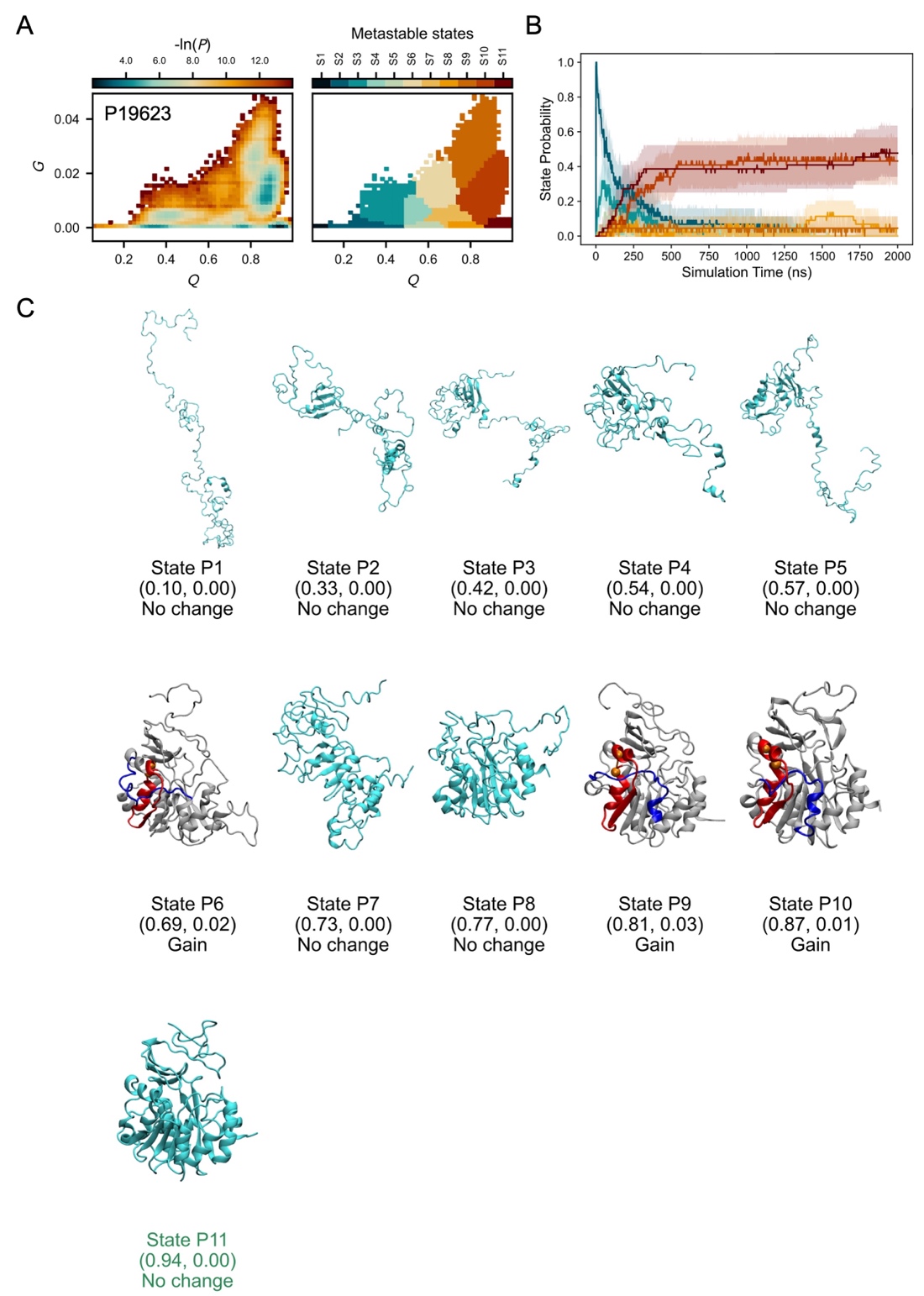

**Figure S5. Simulation structural ensemble of the YU-E protein P19623.** A. The probability distribution (-ln*P*) in the *Q* – *G* space (left) and the corresponding metastable states mapped onto structures from temperature-quench refolding simulations (right). B. Time evolution of metastable-state probabilities. Colors match the state assignments shown in panel A. Shaded regions denote 95% confidence intervals, estimated using bootstrap resampling (10^5^ iterations). C. Representative structures of each metastable state predicted from simulations. *Q* and *G* values at the cluster center, as well as the type of entanglement change (gain, loss or no change), are shown below each structure image. Color scheme follows that used in Fig. 2B, 2D, and 2F.

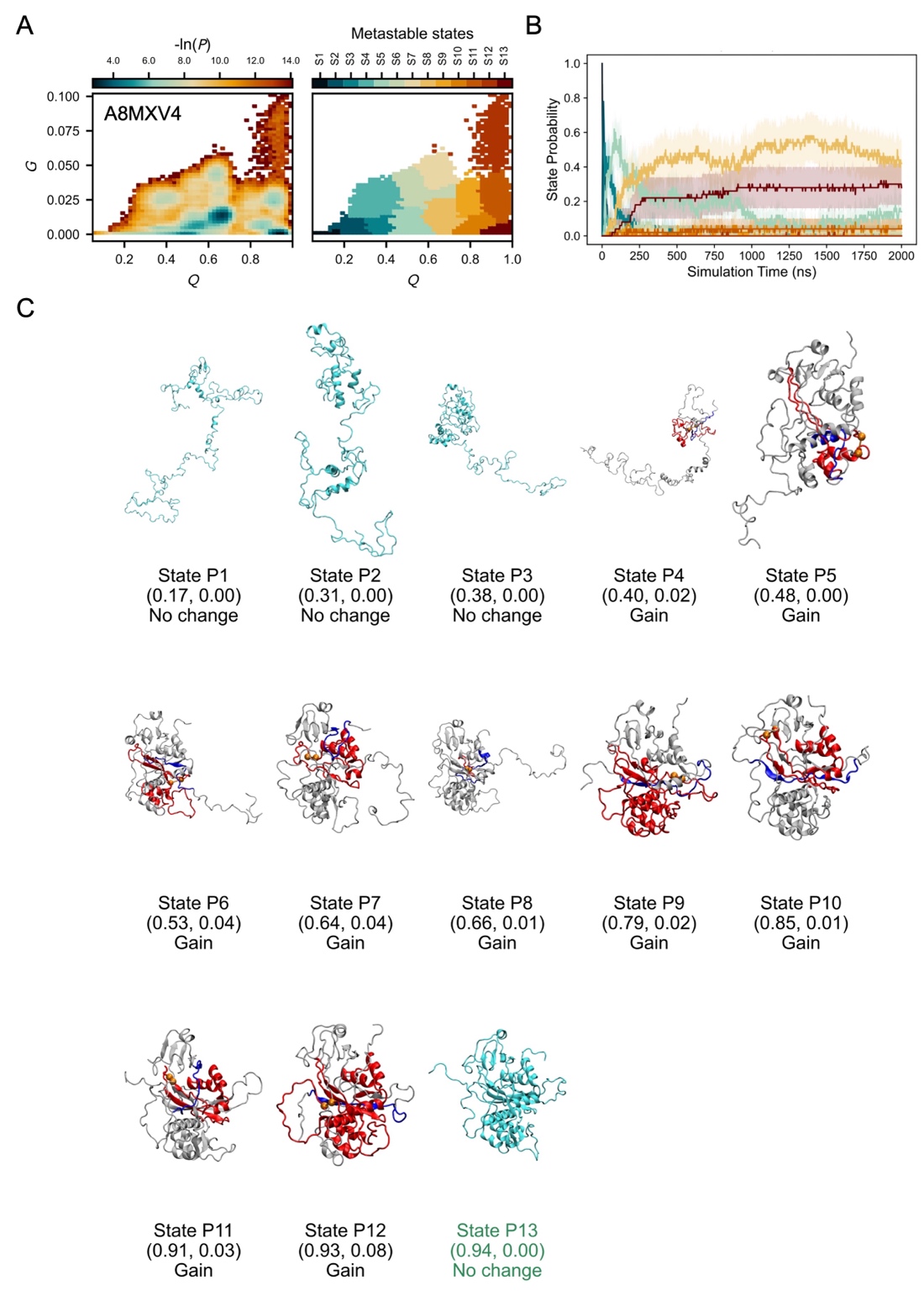

**Figure S6. Simulation structural ensemble of the YU-E protein A8MXV4.** A. The probability distribution (-ln*P*) in the *Q* – *G* space (left) and the corresponding metastable states mapped onto structures from temperature-quench refolding simulations (right). B. Time evolution of metastable-state probabilities. Colors match the state assignments shown in panel A. Shaded regions denote 95% confidence intervals, estimated using bootstrap resampling (10^5^ iterations). C. Representative structures of each metastable state predicted from simulations. *Q* and *G* values at the cluster center, as well as the type of entanglement change (gain, loss or no change), are shown below each structure image. Color scheme follows that used in Fig. 2B, 2D, and 2F.

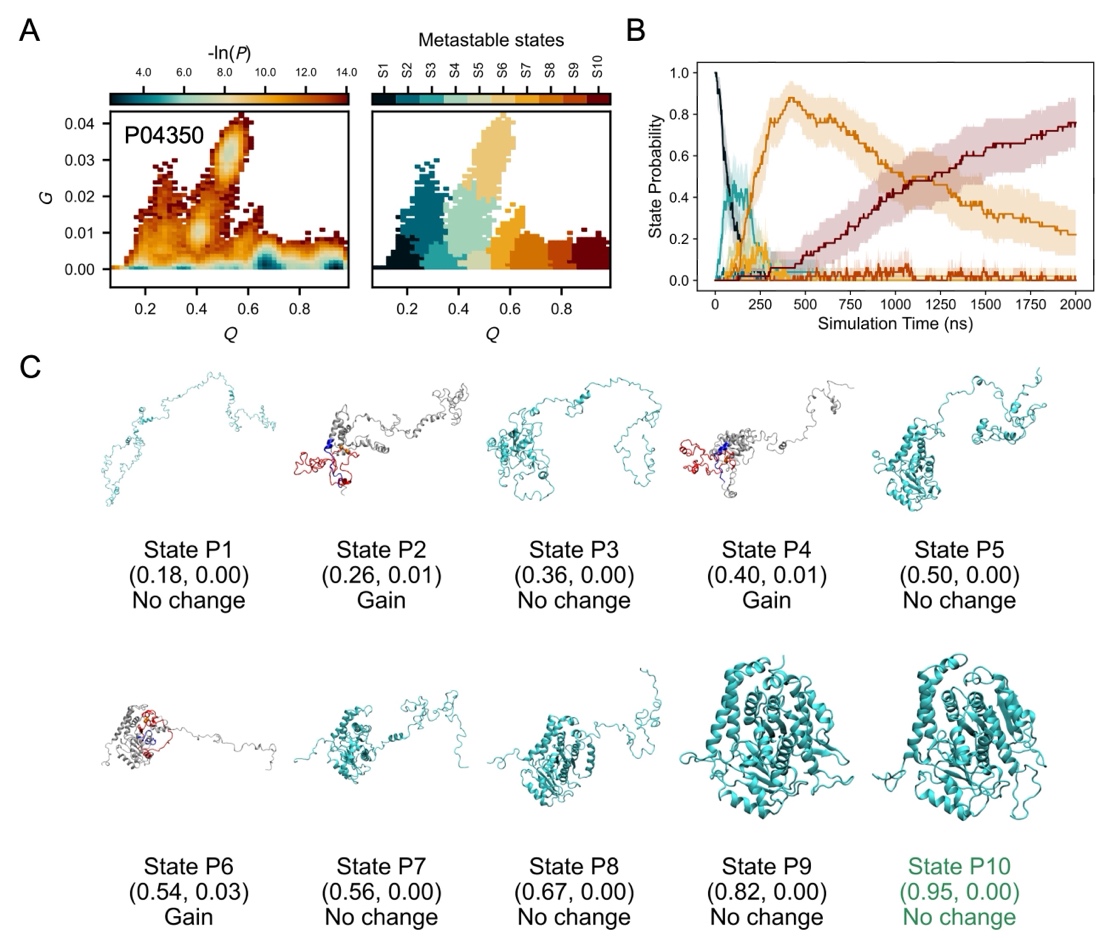

**Figure S7. Simulation structural ensemble of the YU-E protein P04350.** A. The probability distribution (-ln*P*) in the *Q* – *G* space (left) and the corresponding metastable states mapped onto structures from temperature-quench refolding simulations (right). B. Time evolution of metastable-state probabilities. Colors match the state assignments shown in panel A. Shaded regions denote 95% confidence intervals, estimated using bootstrap resampling (10^5^ iterations). C. Representative structures of each metastable state predicted from simulations. *Q* and *G* values at the cluster center, as well as the type of entanglement change (gain, loss or no change), are shown below each structure image. Color scheme follows that used in Fig. 2B, 2D, and 2F.

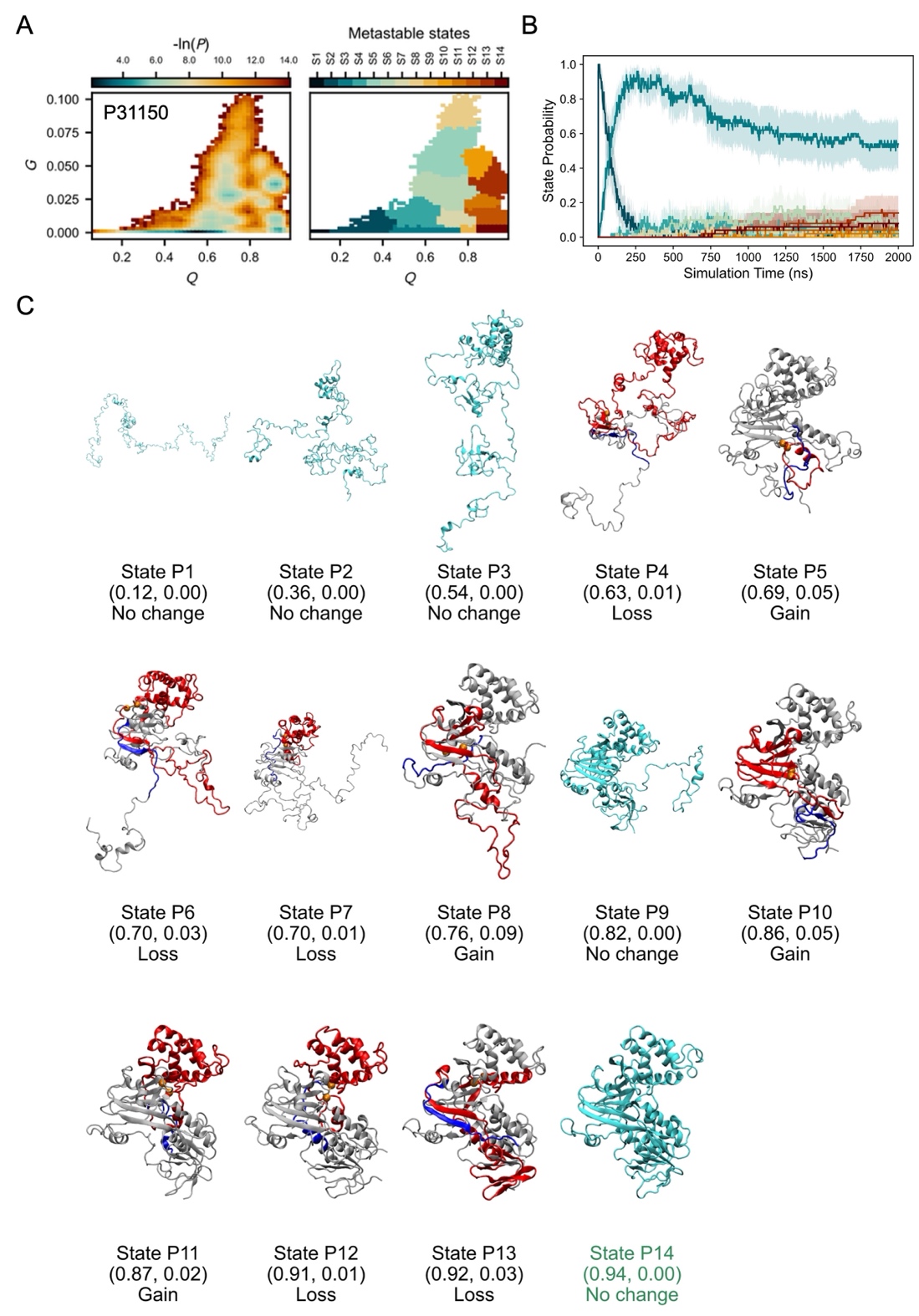

**Figure S8. Simulation structural ensemble of the YU-E protein P31150.** A. The probability distribution (-ln*P*) in the *Q* – *G* space (left) and the corresponding metastable states mapped onto structures from temperature-quench refolding simulations (right). B. Time evolution of metastable-state probabilities. Colors match the state assignments shown in panel A. Shaded regions denote 95% confidence intervals, estimated using bootstrap resampling (10^5^ iterations). C. Representative structures of each metastable state predicted from simulations. *Q* and *G* values at the cluster center, as well as the type of entanglement change (gain, loss or no change), are shown below each structure image. Color scheme follows that used in Fig. 2B, 2D, and 2F.

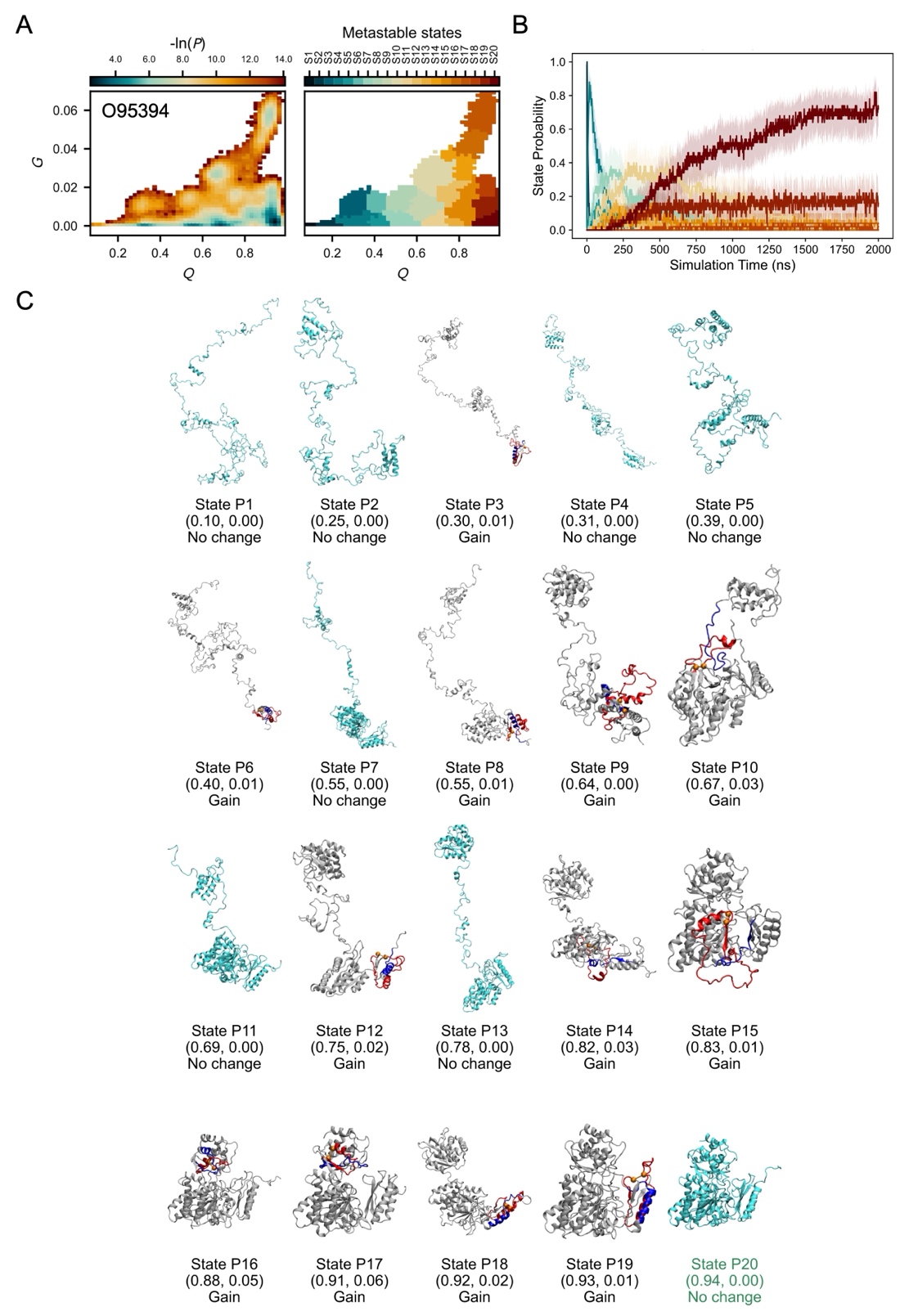

**Figure S9. Simulation structural ensemble of the YU-E protein O95394.** A. The probability distribution (-ln*P*) in the *Q* – *G* space (left) and the corresponding metastable states mapped onto structures from temperature-quench refolding simulations (right). B. Time evolution of metastable-state probabilities. Colors match the state assignments shown in panel A. Shaded regions denote 95% confidence intervals, estimated using bootstrap resampling (10^5^ iterations). C. Representative structures of each metastable state predicted from simulations. *Q* and *G* values at the cluster center, as well as the type of entanglement change (gain, loss or no change), are shown below each structure image. Color scheme follows that used in Fig. 2B, 2D, and 2F.

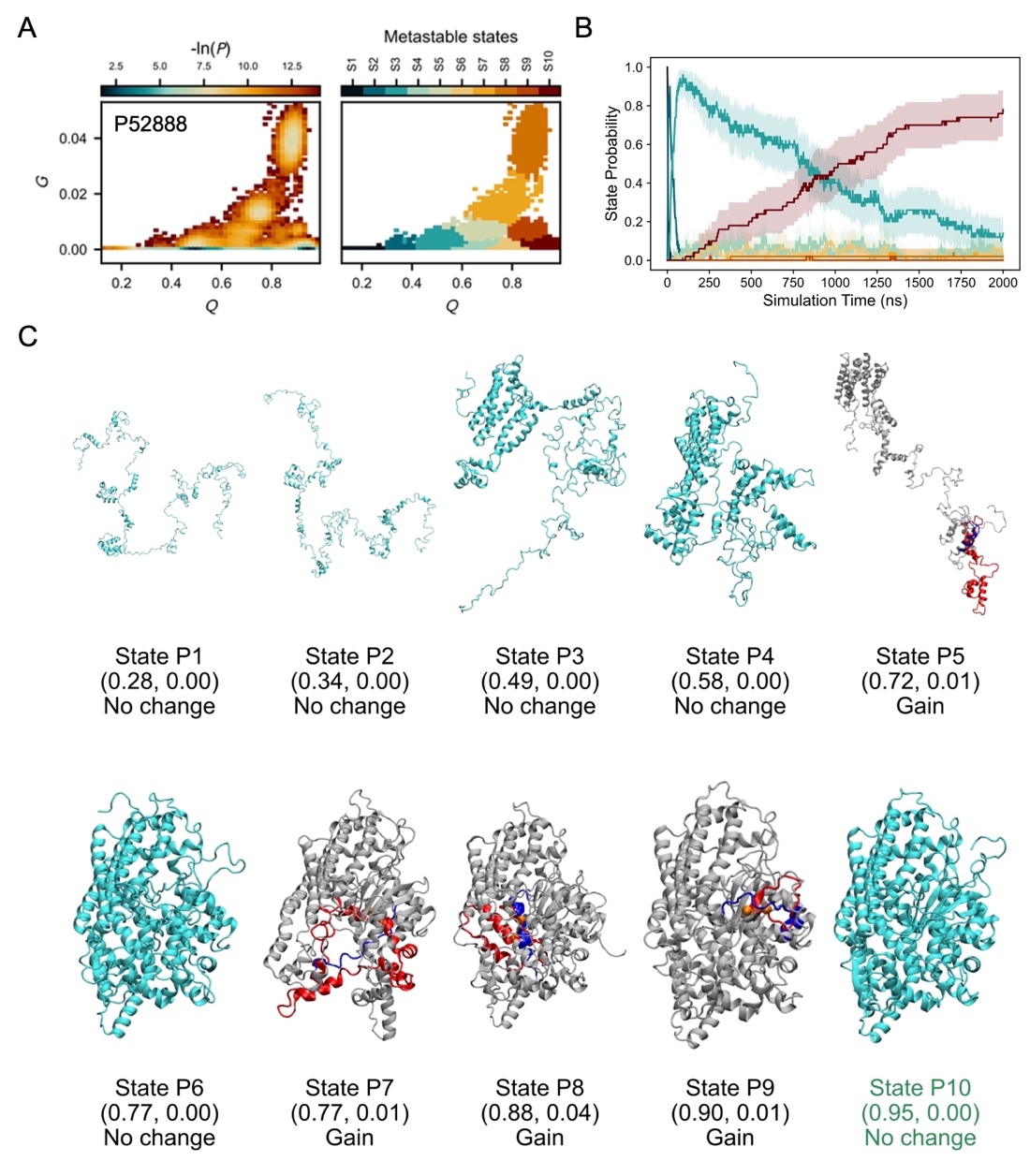

**Figure S10. Simulation structural ensemble of the YU-E protein P52888.** A. The probability distribution (-ln*P*) in the *Q* – *G* space (left) and the corresponding metastable states mapped onto structures from temperature-quench refolding simulations (right). B. Time evolution of metastable-state probabilities. Colors match the state assignments shown in panel A. Shaded regions denote 95% confidence intervals, estimated using bootstrap resampling (10^5^ iterations). C. Representative structures of each metastable state predicted from simulations. *Q* and *G* values at the cluster center, as well as the type of entanglement change (gain, loss or no change), are shown below each structure image. Color scheme follows that used in Fig. 2B, 2D, and 2F.

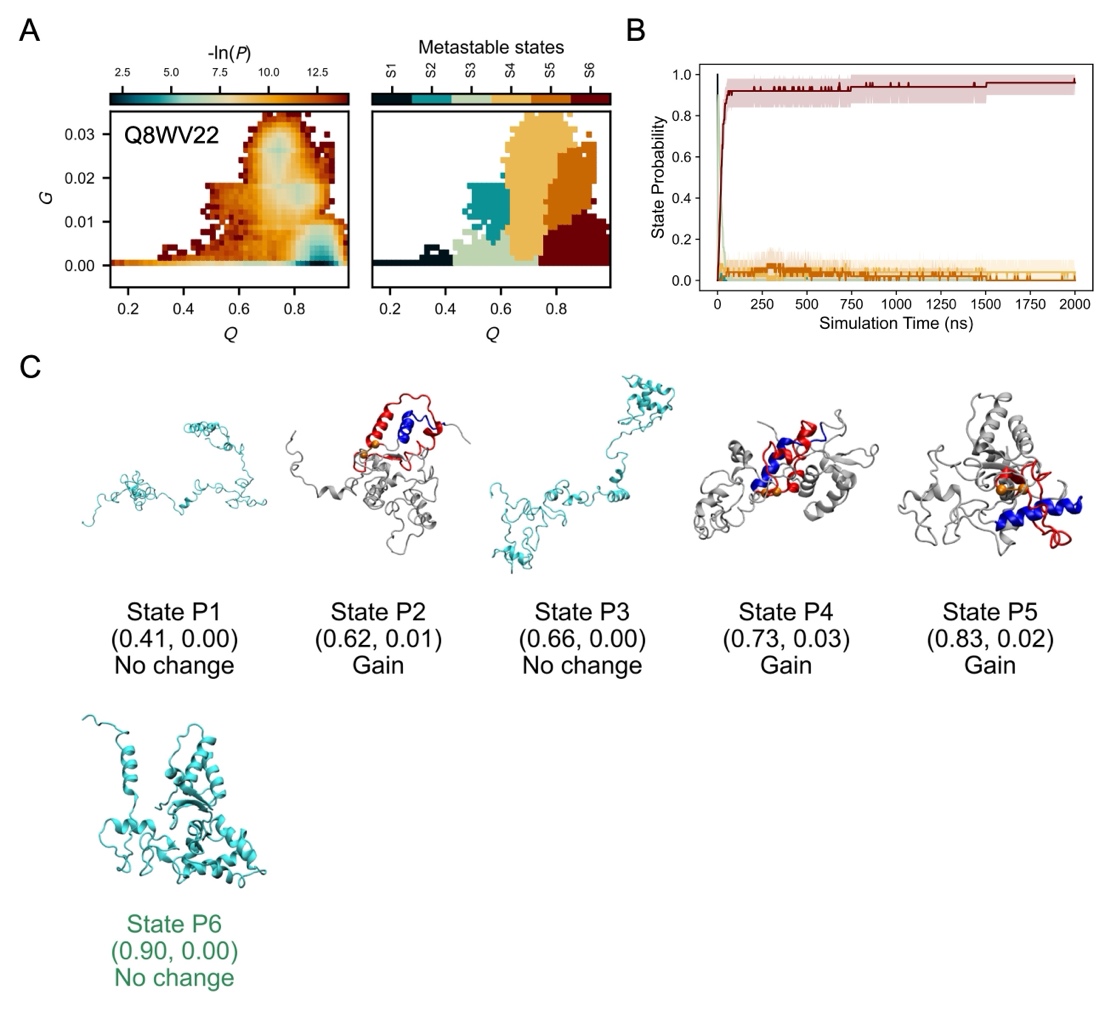

**Figure S11. Simulation structural ensemble of the NU-NE protein Q8WV22.** A. The probability distribution (-ln*P*) in the *Q* – *G* space (left) and the corresponding metastable states mapped onto structures from temperature-quench refolding simulations (right). B. Time evolution of metastable-state probabilities. Colors match the state assignments shown in panel A. Shaded regions denote 95% confidence intervals, estimated using bootstrap resampling (10^5^ iterations). C. Representative structures of each metastable state predicted from simulations. *Q* and *G* values at the cluster center, as well as the type of entanglement change (gain, loss or no change), are shown below each structure image. Color scheme follows that used in Fig. 2B, 2D, and 2F.

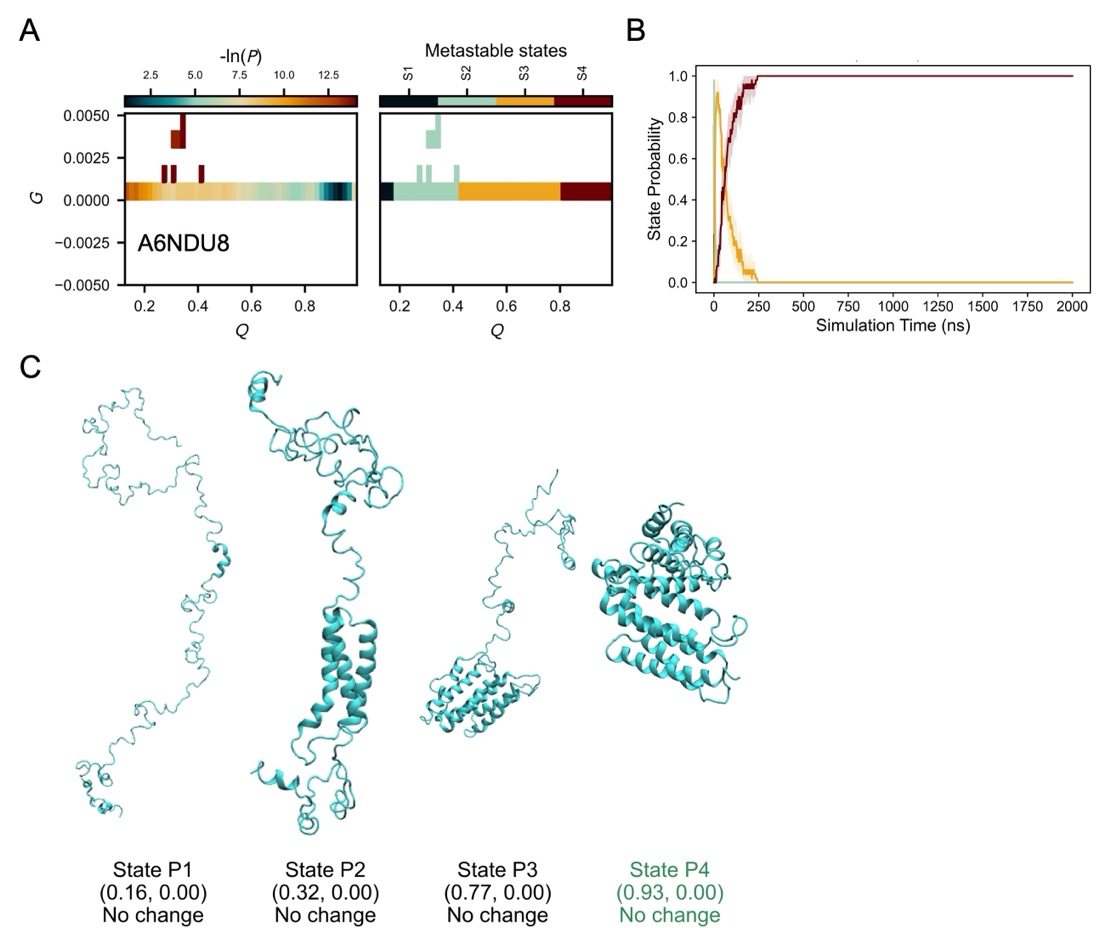

**Figure S12. Simulation structural ensemble of the NU-NE protein A6NDU8.** A. The probability distribution (-ln*P*) in the *Q* – *G* space (left) and the corresponding metastable states mapped onto structures from temperature-quench refolding simulations (right). B. Time evolution of metastable-state probabilities. Colors match the state assignments shown in panel A. Shaded regions denote 95% confidence intervals, estimated using bootstrap resampling (10^5^ iterations). C. Representative structures of each metastable state predicted from simulations. *Q* and *G* values at the cluster center, as well as the type of entanglement change (gain, loss or no change), are shown below each structure image. Color scheme follows that used in Fig. 2B, 2D, and 2F.

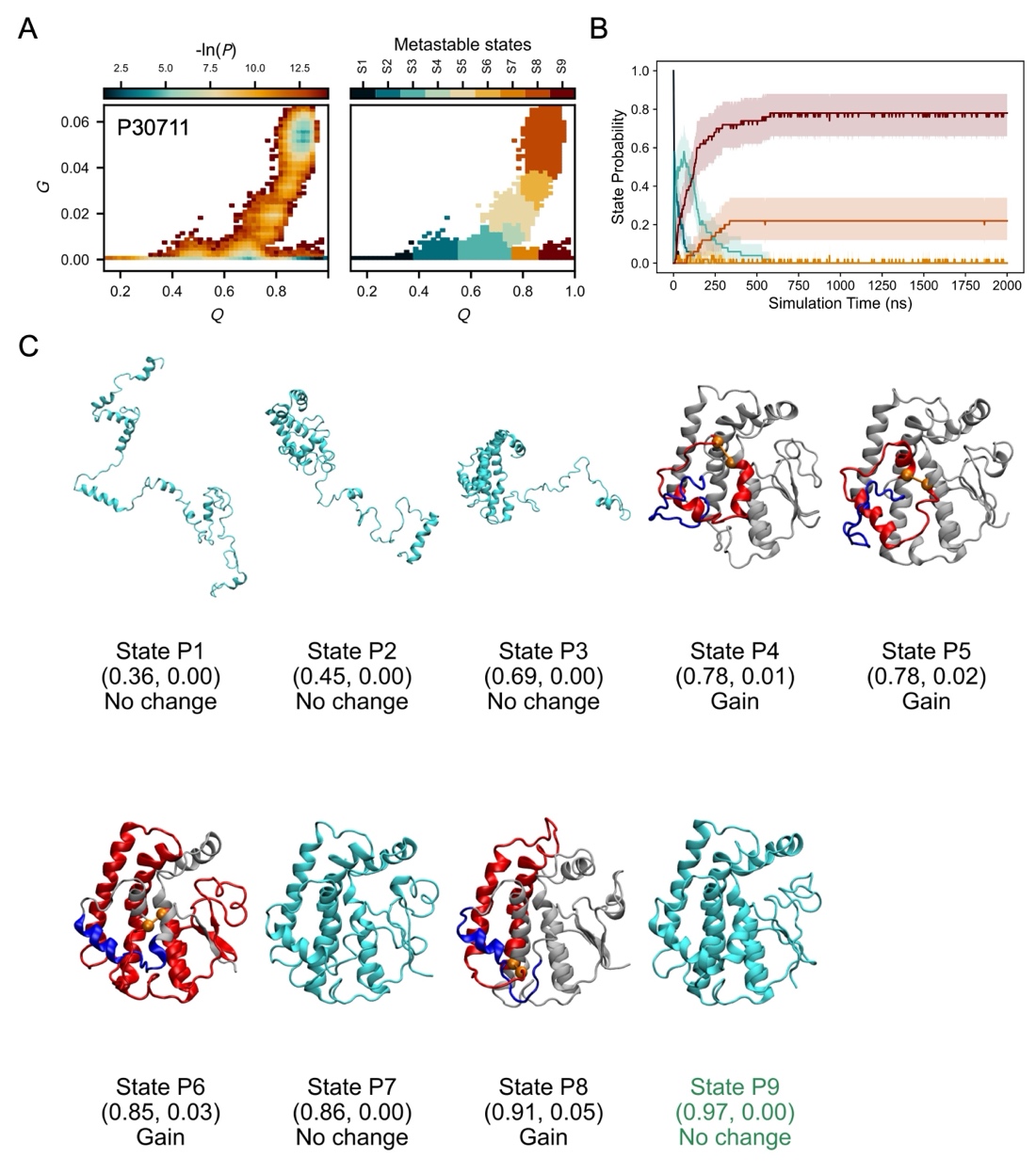

**Figure S13. Simulation structural ensemble of the NU-NE protein P30711.** A. The probability distribution (-ln*P*) in the *Q* – *G* space (left) and the corresponding metastable states mapped onto structures from temperature-quench refolding simulations (right). B. Time evolution of metastable-state probabilities. Colors match the state assignments shown in panel A. Shaded regions denote 95% confidence intervals, estimated using bootstrap resampling (10^5^ iterations). C. Representative structures of each metastable state predicted from simulations. *Q* and *G* values at the cluster center, as well as the type of entanglement change (gain, loss or no change), are shown below each structure image. Color scheme follows that used in Fig. 2B, 2D, and 2F.

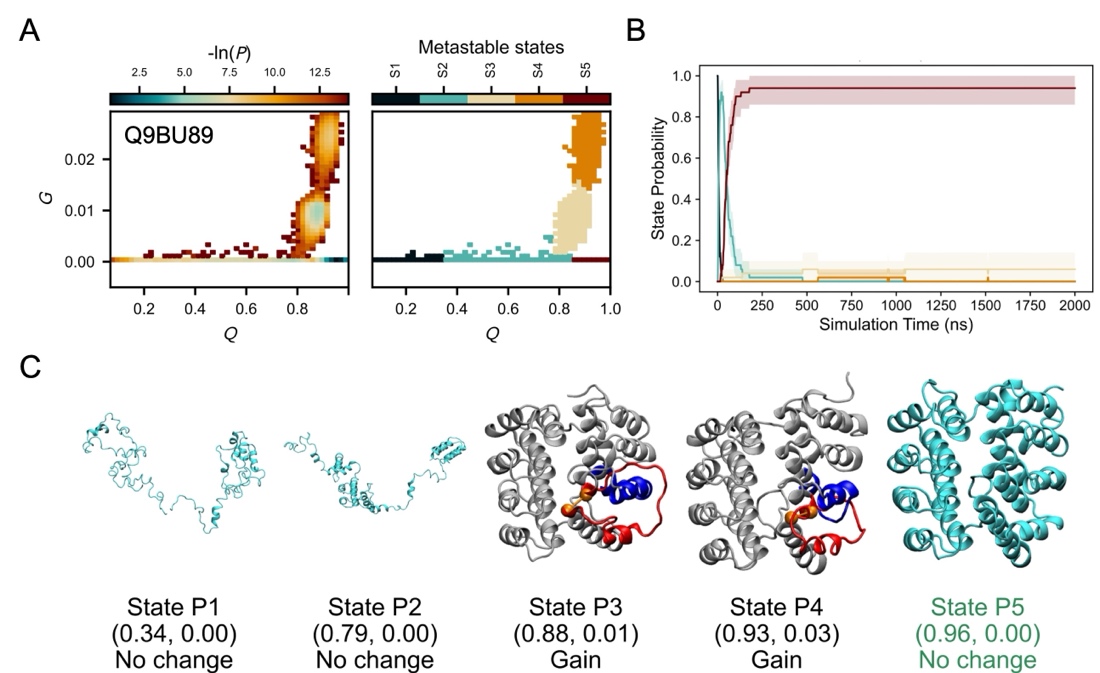

**Figure S14. Simulation structural ensemble of the NU-NE protein Q9BU89.** A. The probability distribution (-ln*P*) in the *Q* – *G* space (left) and the corresponding metastable states mapped onto structures from temperature-quench refolding simulations (right). B. Time evolution of metastable-state probabilities. Colors match the state assignments shown in panel A. Shaded regions denote 95% confidence intervals, estimated using bootstrap resampling (10^5^ iterations). C. Representative structures of each metastable state predicted from simulations. *Q* and *G* values at the cluster center, as well as the type of entanglement change (gain, loss or no change), are shown below each structure image. Color scheme follows that used in Fig. 2B, 2D, and 2F.

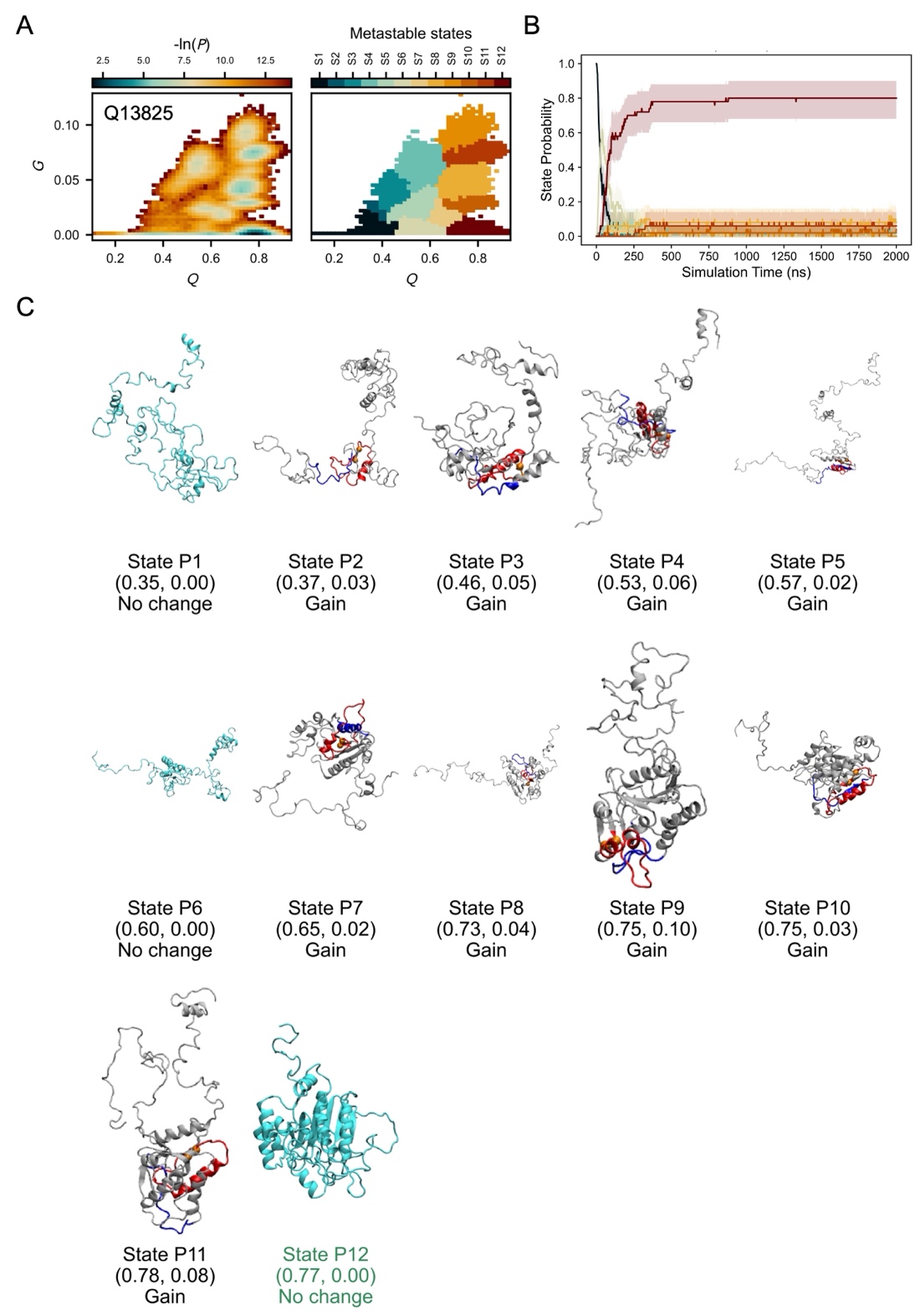

**Figure S15. Simulation structural ensemble of the NU-NE protein Q13825.** A. The probability distribution (-ln*P*) in the *Q* – *G* space (left) and the corresponding metastable states mapped onto structures from temperature-quench refolding simulations (right). B. Time evolution of metastable-state probabilities. Colors match the state assignments shown in panel A. Shaded regions denote 95% confidence intervals, estimated using bootstrap resampling (10^5^ iterations). C. Representative structures of each metastable state predicted from simulations. *Q* and *G* values at the cluster center, as well as the type of entanglement change (gain, loss or no change), are shown below each structure image. Color scheme follows that used in Fig. 2B, 2D, and 2F.

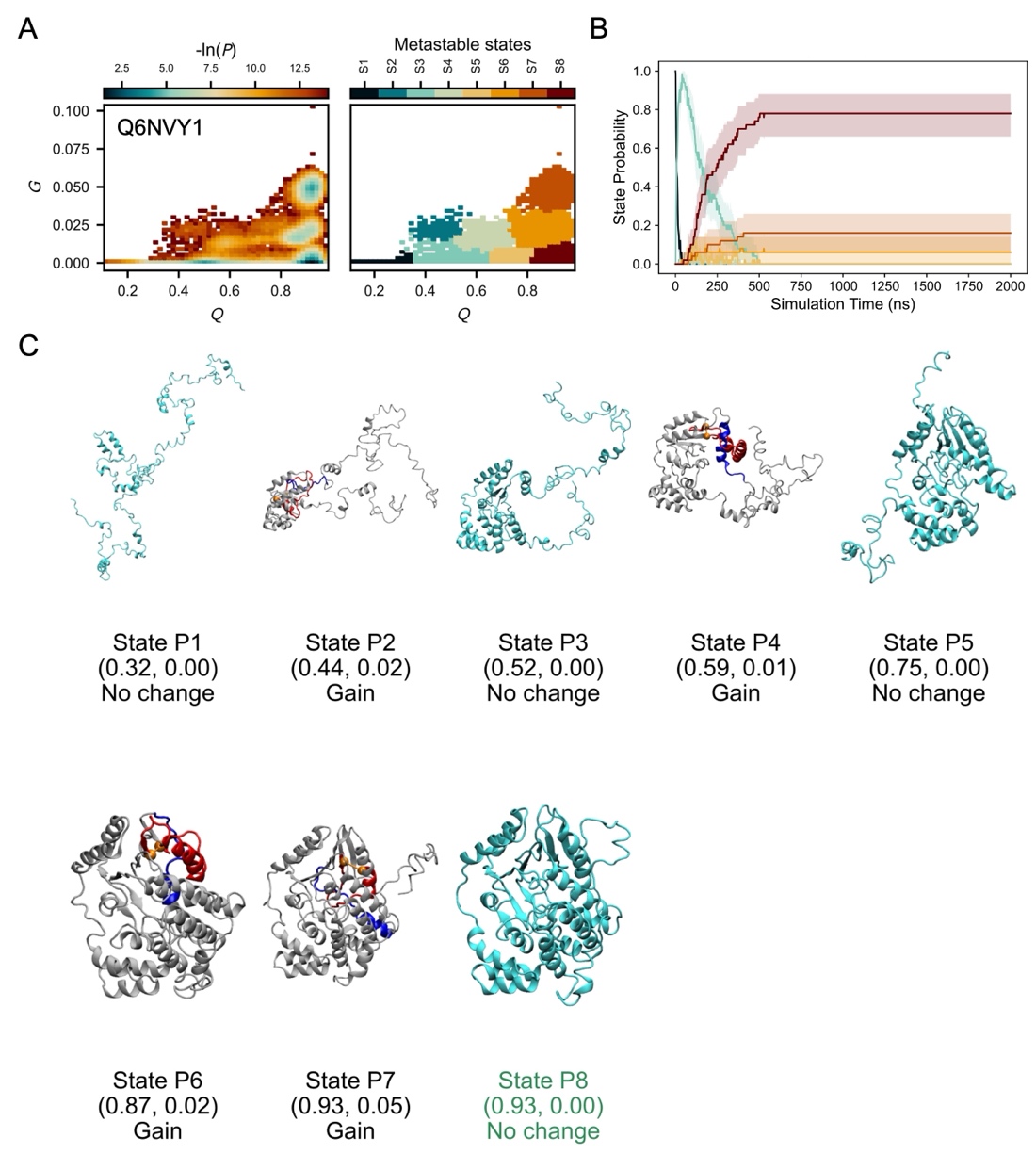

**Figure S16. Simulation structural ensemble of the NU-NE protein Q6NVY1.** A. The probability distribution (-ln*P*) in the *Q* – *G* space (left) and the corresponding metastable states mapped onto structures from temperature-quench refolding simulations (right). B. Time evolution of metastable-state probabilities. Colors match the state assignments shown in panel A. Shaded regions denote 95% confidence intervals, estimated using bootstrap resampling (10^5^ iterations). C. Representative structures of each metastable state predicted from simulations. *Q* and *G* values at the cluster center, as well as the type of entanglement change (gain, loss or no change), are shown below each structure image. Color scheme follows that used in Fig. 2B, 2D, and 2F.

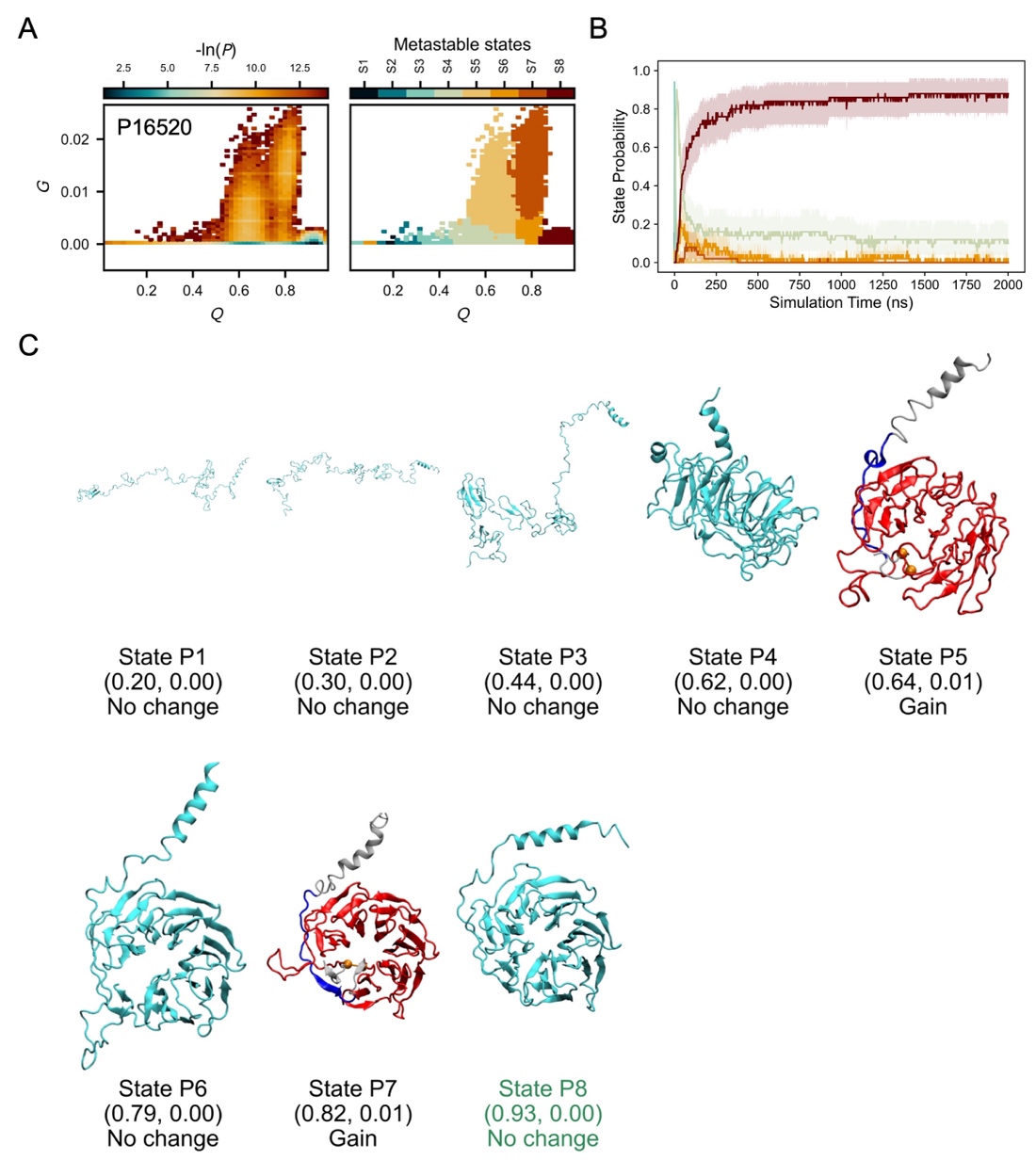

**Figure S17. Simulation structural ensemble of the NU-NE protein P16520.** A. The probability distribution (-ln*P*) in the *Q* – *G* space (left) and the corresponding metastable states mapped onto structures from temperature-quench refolding simulations (right). B. Time evolution of metastable-state probabilities. Colors match the state assignments shown in panel A. Shaded regions denote 95% confidence intervals, estimated using bootstrap resampling (10^5^ iterations). C. Representative structures of each metastable state predicted from simulations. *Q* and *G* values at the cluster center, as well as the type of entanglement change (gain, loss or no change), are shown below each structure image. Color scheme follows that used in Fig. 2B, 2D, and 2F.

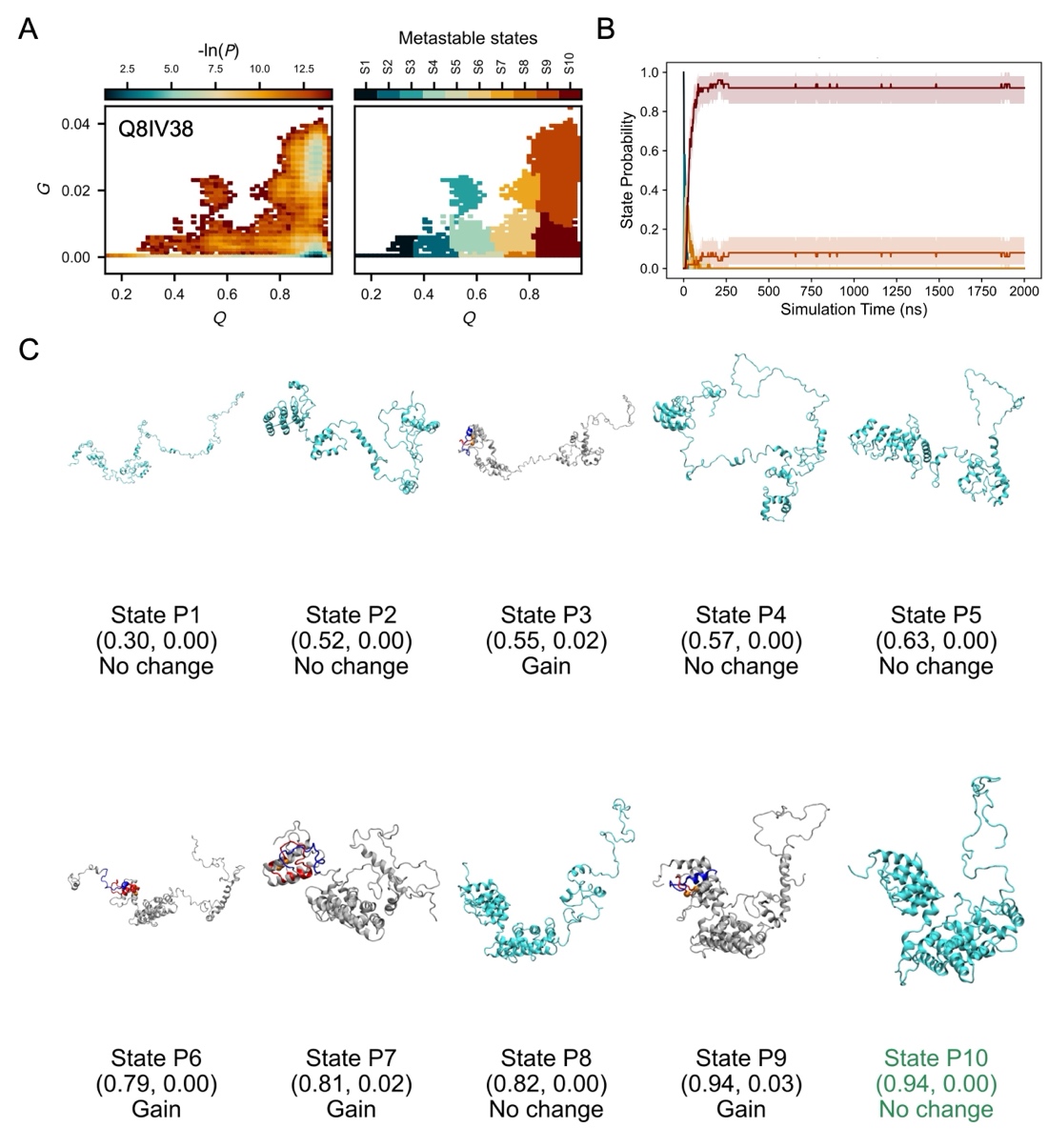

**Figure S18. Simulation structural ensemble of the NU-NE protein Q8IV38.** A. The probability distribution (-ln*P*) in the *Q* – *G* space (left) and the corresponding metastable states mapped onto structures from temperature-quench refolding simulations (right). B. Time evolution of metastable-state probabilities. Colors match the state assignments shown in panel A. Shaded regions denote 95% confidence intervals, estimated using bootstrap resampling (10^5^ iterations). C. Representative structures of each metastable state predicted from simulations. *Q* and *G* values at the cluster center, as well as the type of entanglement change (gain, loss or no change), are shown below each structure image. Color scheme follows that used in Fig. 2B, 2D, and 2F.

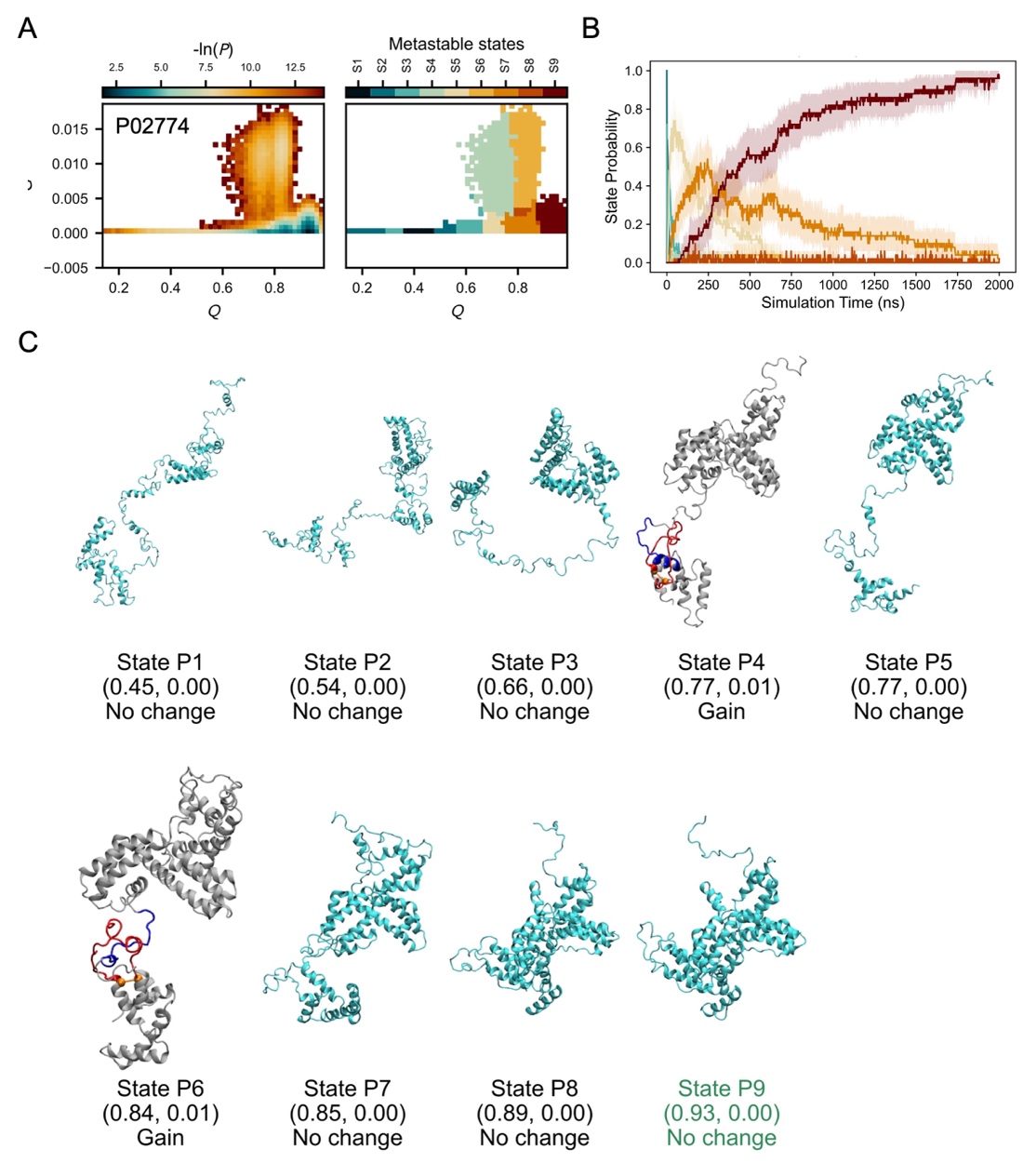

**Figure S19. Simulation structural ensemble of the NU-NE protein P02774.** A. The probability distribution (-ln*P*) in the *Q* – *G* space (left) and the corresponding metastable states mapped onto structures from temperature-quench refolding simulations (right). B. Time evolution of metastable-state probabilities. Colors match the state assignments shown in panel A. Shaded regions denote 95% confidence intervals, estimated using bootstrap resampling (10^5^ iterations). C. Representative structures of each metastable state predicted from simulations. *Q* and *G* values at the cluster center, as well as the type of entanglement change (gain, loss or no change), are shown below each structure image. Color scheme follows that used in Fig. 2B, 2D, and 2F.

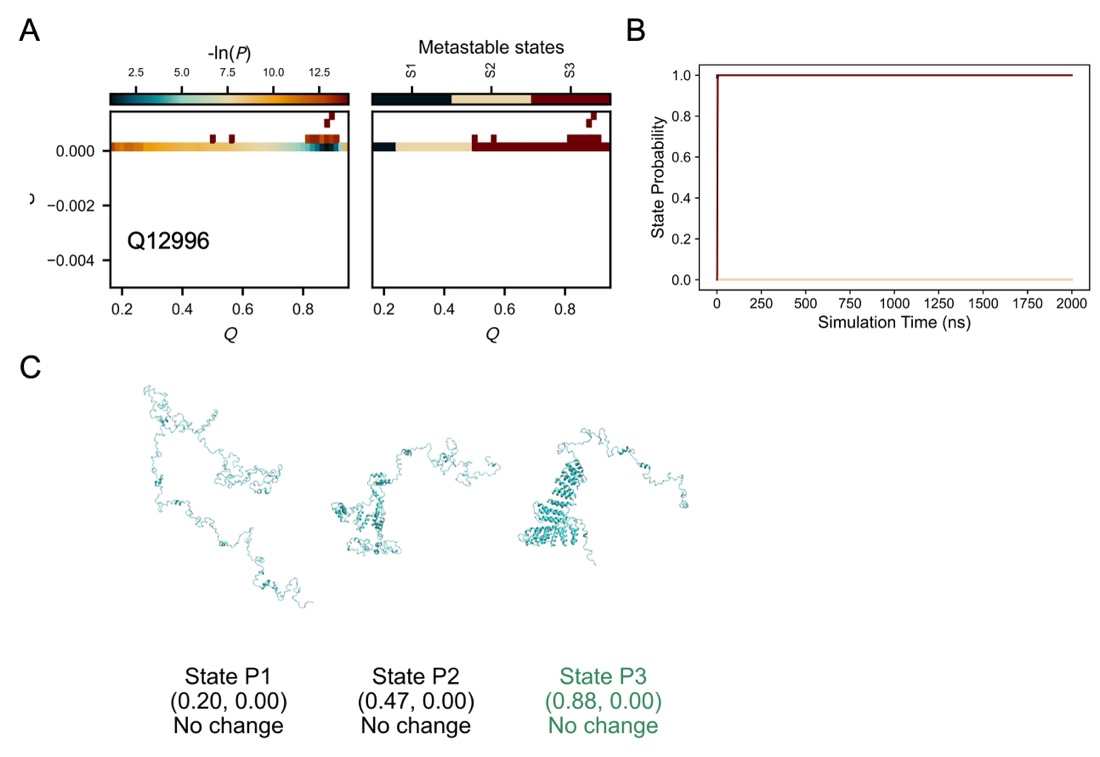

**Figure S20. Simulation structural ensemble of the NU-NE protein Q12996.** A. The probability distribution (-ln*P*) in the *Q* – *G* space (left) and the corresponding metastable states mapped onto structures from temperature-quench refolding simulations (right). B. Time evolution of metastable-state probabilities. Colors match the state assignments shown in panel A. Shaded regions denote 95% confidence intervals, estimated using bootstrap resampling (10^5^ iterations). C. Representative structures of each metastable state predicted from simulations. *Q* and *G* values at the cluster center, as well as the type of entanglement change (gain, loss or no change), are shown below each structure image. Color scheme follows that used in Fig. 2B, 2D, and 2F.

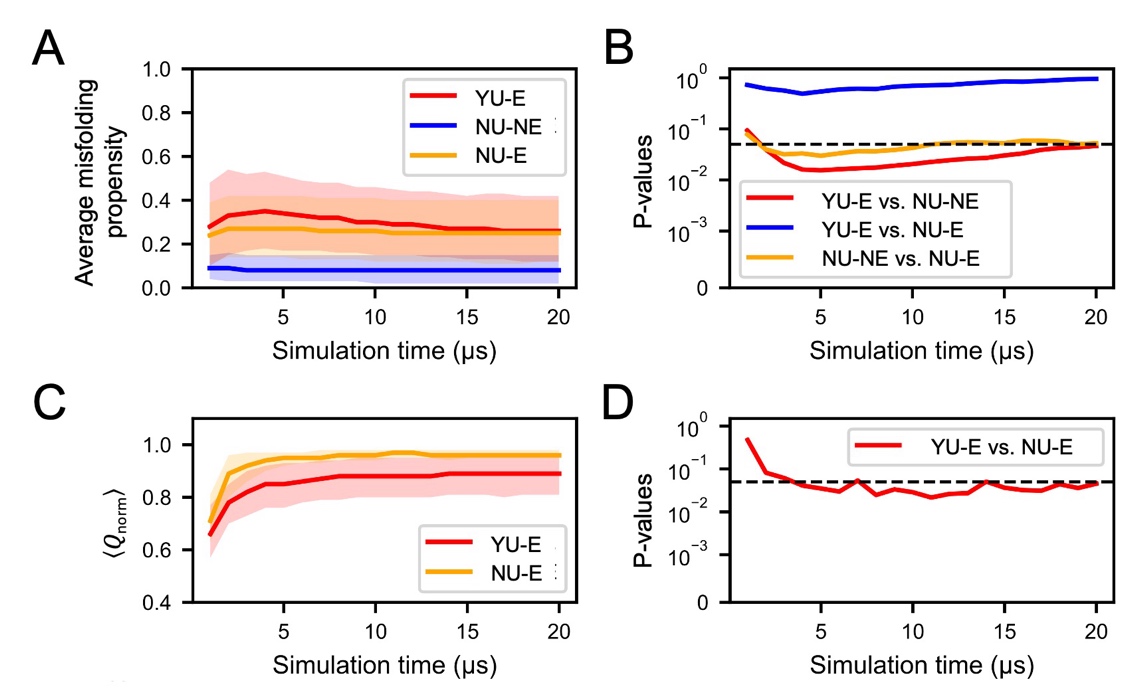

**Figure S21. Average misfolding propensity and** $\left\langle\boldsymbol{Q}_{\text{norm}} \right\rangle$ **along the extended simulation trajectories.** A. Average misfolding propensity of YU-E (red), NU-NE (blue) and NU-E (orange) proteins. Shaded regions denote 95% confidence intervals, estimated using hierarchical bootstrap resampling (10^5^ iterations). B. P-values for the differences in misfolding propensities between YU-E and NU-NE (red), YU-E and NU-E (blue) and NU-NE and NU-E (orange) proteins, estimated using two-sided hierarchical permutation test for 10^6^ iterations. The horizontal dashed line marks the significance level at 0.05. C. $\left\langle Q_{\text{norm}} \right\rangle$ of YU-E (red) and NU-E (orange) proteins. Shaded regions denote 95% confidence intervals, estimated using hierarchical bootstrap resampling (10^5^ iterations). D. P-values for the differences in $\left\langle Q_{\text{norm}} \right\rangle$ between YU-E and NU-E (red) proteins, estimated using two-sided hierarchical permutation test for 10^6^ iterations. The horizontal dashed line marks the significance level at 0.05.

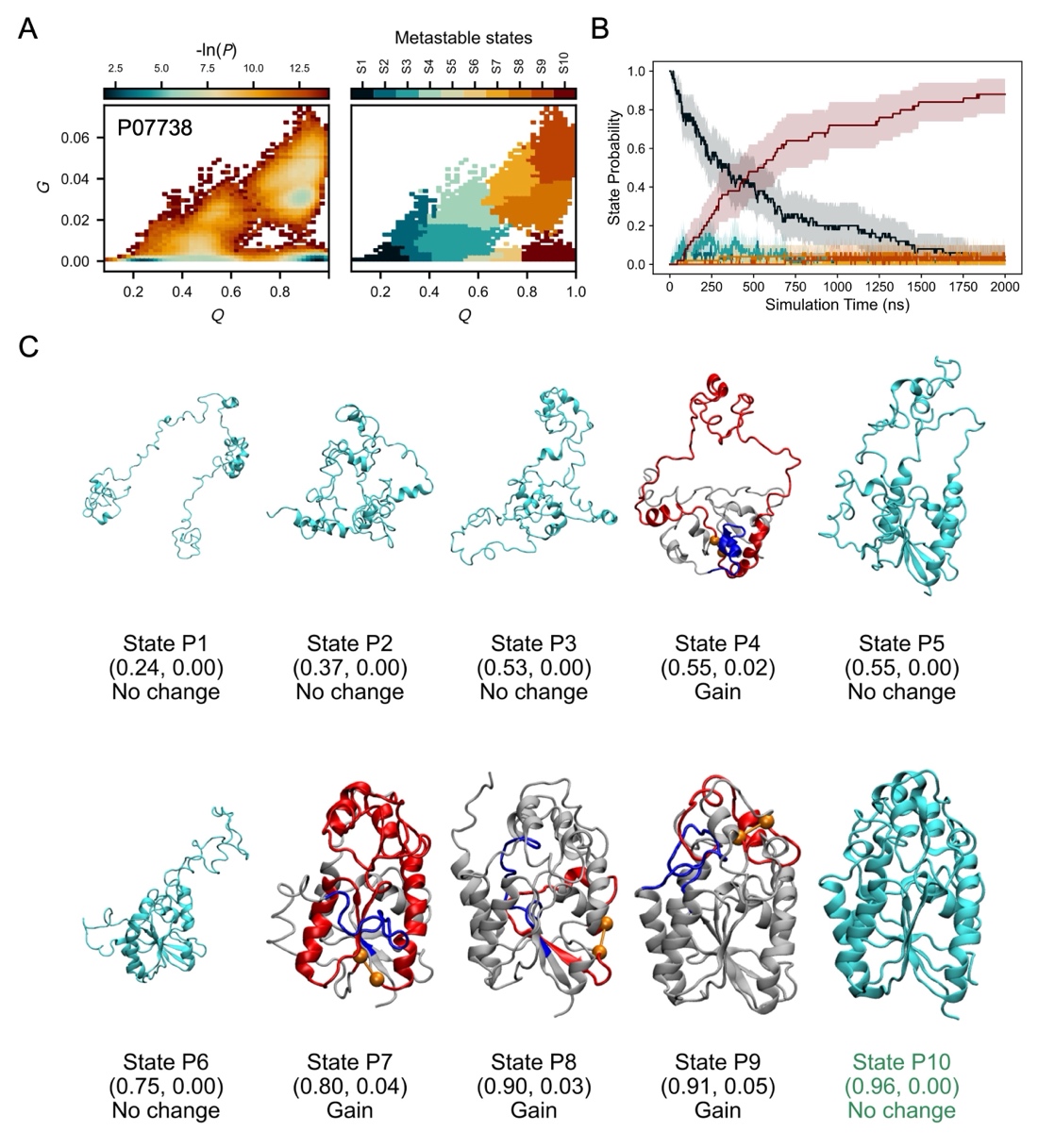

**Figure S22. Simulation structural ensemble of the NU-E protein P07738.** A. The probability distribution (-ln*P*) in the *Q* – *G* space (left) and the corresponding metastable states mapped onto structures from temperature-quench refolding simulations (right). B. Time evolution of metastable-state probabilities. Colors match the state assignments shown in panel A. Shaded regions denote 95% confidence intervals, estimated using bootstrap resampling (10^5^ iterations). C. Representative structures of each metastable state predicted from simulations. *Q* and *G* values at the cluster center, as well as the type of entanglement change (gain, loss or no change), are shown below each structure image. Color scheme follows that used in Fig. 2B, 2D, and 2F.

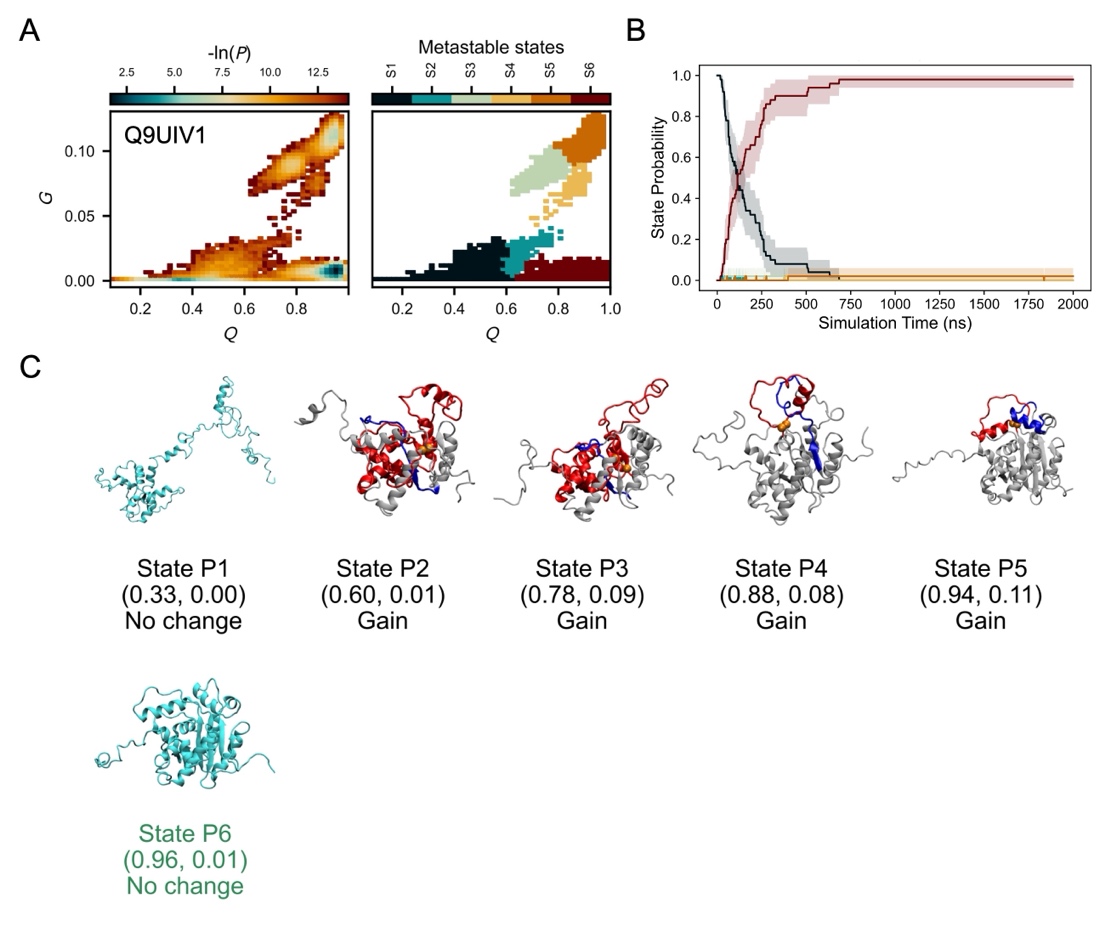

**Figure S23. Simulation structural ensemble of the NU-E protein Q9UIV1.** A. The probability distribution (-ln*P*) in the *Q* – *G* space (left) and the corresponding metastable states mapped onto structures from temperature-quench refolding simulations (right). B. Time evolution of metastable-state probabilities. Colors match the state assignments shown in panel A. Shaded regions denote 95% confidence intervals, estimated using bootstrap resampling (10^5^ iterations). C. Representative structures of each metastable state predicted from simulations. *Q* and *G* values at the cluster center, as well as the type of entanglement change (gain, loss or no change), are shown below each structure image. Color scheme follows that used in Fig. 2B, 2D, and 2F.

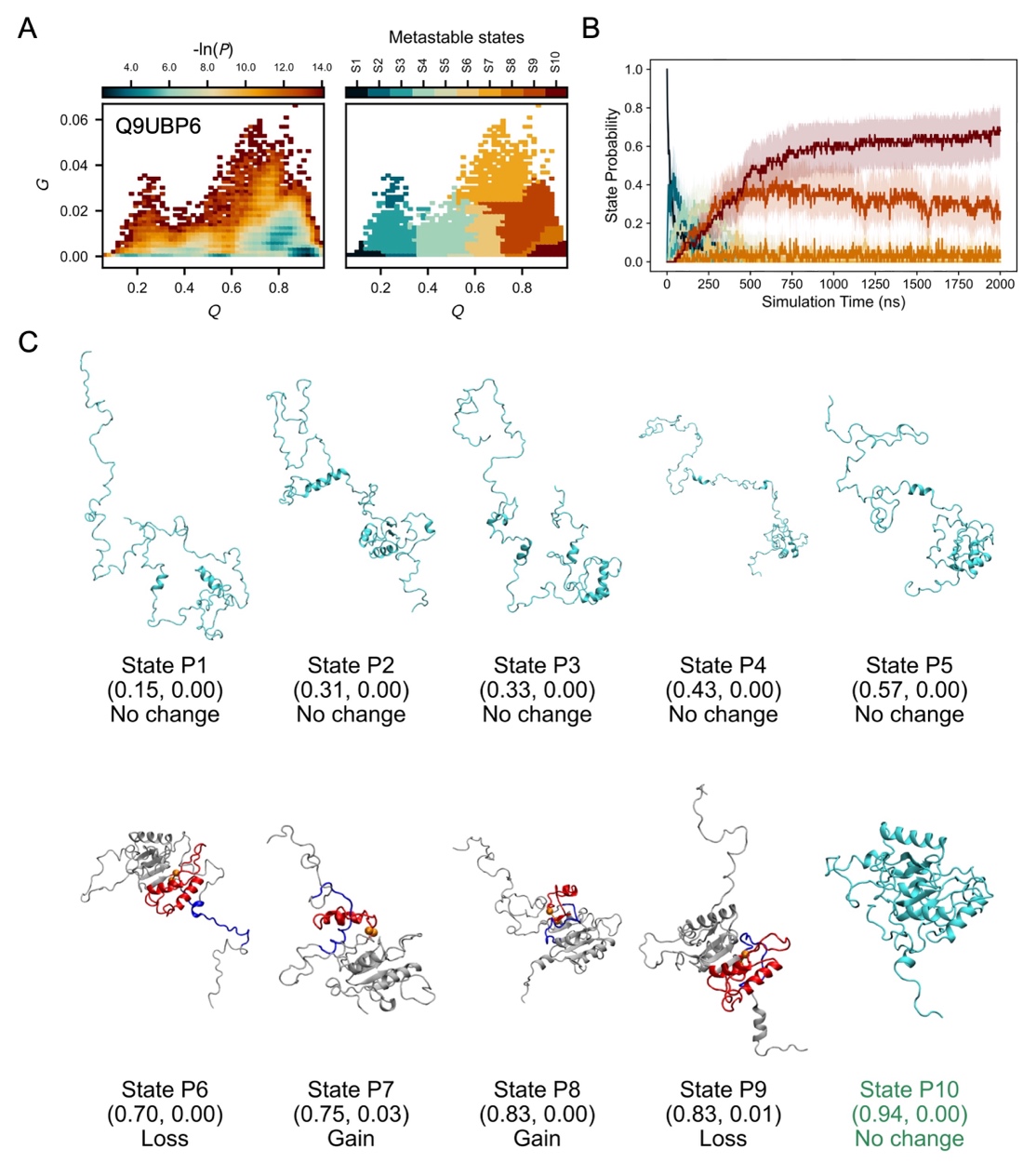

**Figure S24. Simulation structural ensemble of the NU-E protein Q9UBP6.** A. The probability distribution (-ln*P*) in the *Q* – *G* space (left) and the corresponding metastable states mapped onto structures from temperature-quench refolding simulations (right). B. Time evolution of metastable-state probabilities. Colors match the state assignments shown in panel A. Shaded regions denote 95% confidence intervals, estimated using bootstrap resampling (10^5^ iterations). C. Representative structures of each metastable state predicted from simulations. *Q* and *G* values at the cluster center, as well as the type of entanglement change (gain, loss or no change), are shown below each structure image. Color scheme follows that used in Fig. 2B, 2D, and 2F.

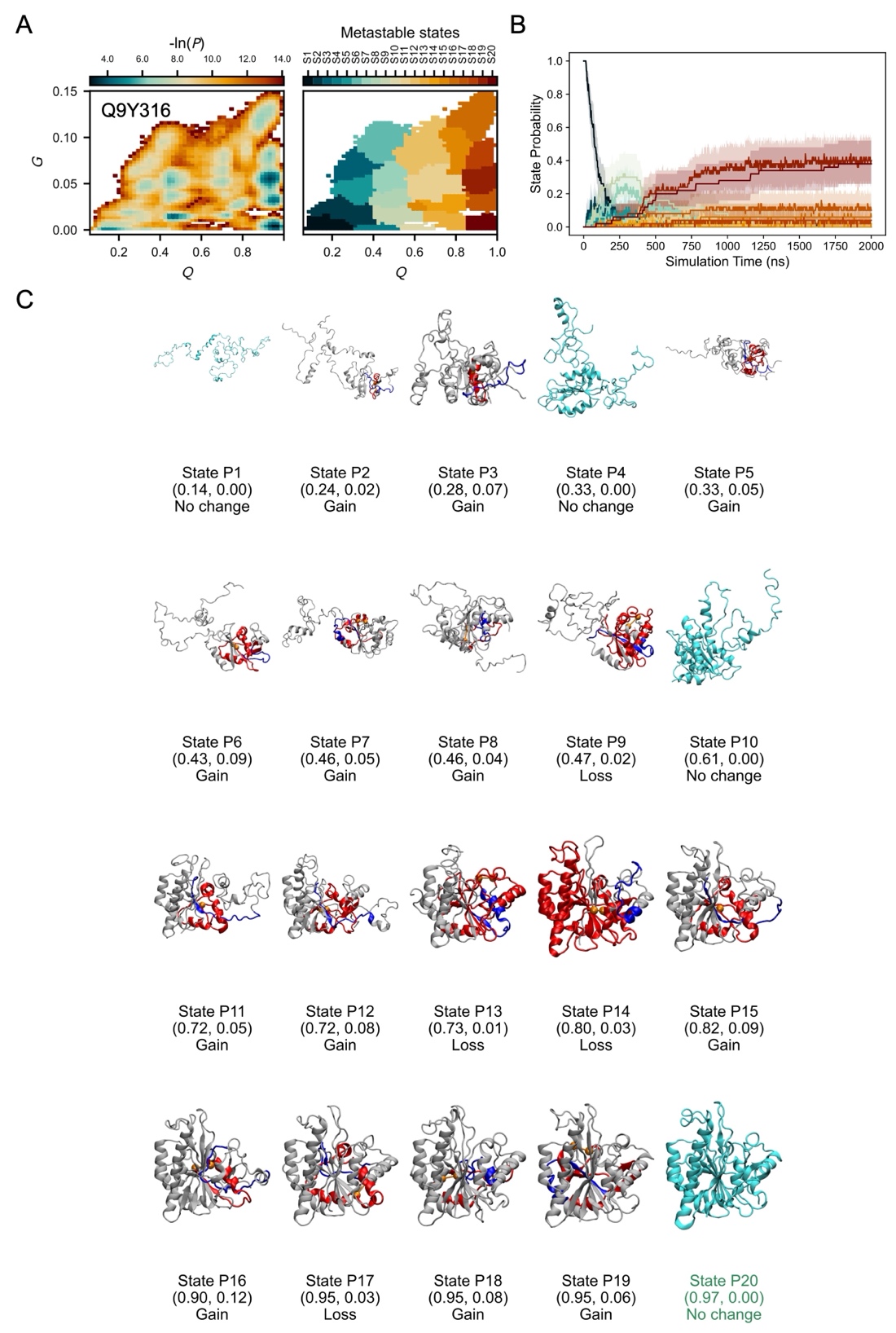

**Figure S25. Simulation structural ensemble of the NU-E protein Q9Y316.** A. The probability distribution (-ln*P*) in the *Q* – *G* space (left) and the corresponding metastable states mapped onto structures from temperature-quench refolding simulations (right). B. Time evolution of metastable-state probabilities. Colors match the state assignments shown in panel A. Shaded regions denote 95% confidence intervals, estimated using bootstrap resampling (10^5^ iterations). C. Representative structures of each metastable state predicted from simulations. *Q* and *G* values at the cluster center, as well as the type of entanglement change (gain, loss or no change), are shown below each structure image. Color scheme follows that used in Fig. 2B, 2D, and 2F.

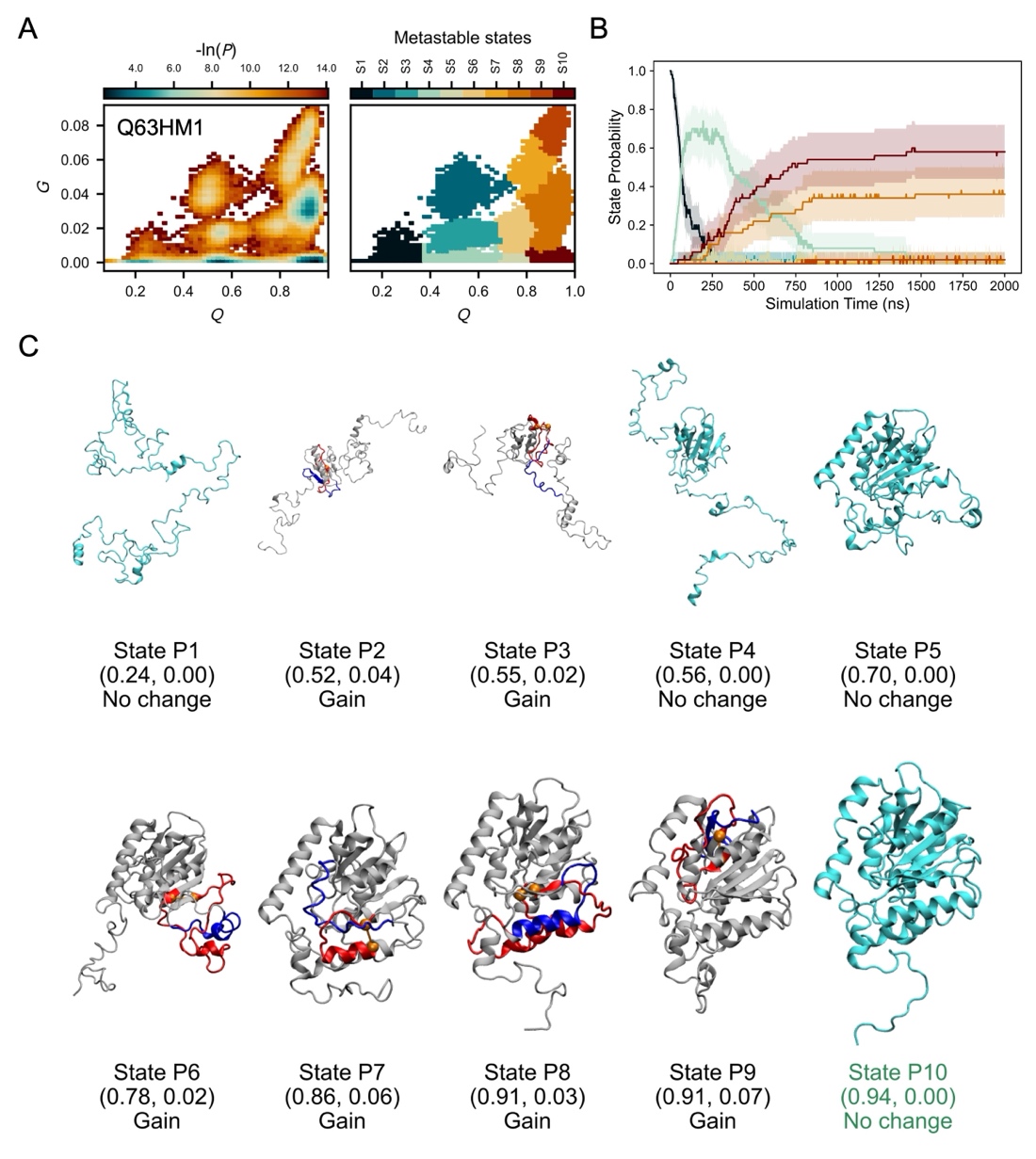

**Figure S26. Simulation structural ensemble of the NU-E protein Q63HM1.** A. The probability distribution (-ln*P*) in the *Q* – *G* space (left) and the corresponding metastable states mapped onto structures from temperature-quench refolding simulations (right). B. Time evolution of metastable-state probabilities. Colors match the state assignments shown in panel A. Shaded regions denote 95% confidence intervals, estimated using bootstrap resampling (10^5^ iterations). C. Representative structures of each metastable state predicted from simulations. *Q* and *G* values at the cluster center, as well as the type of entanglement change (gain, loss or no change), are shown below each structure image. Color scheme follows that used in Fig. 2B, 2D, and 2F.

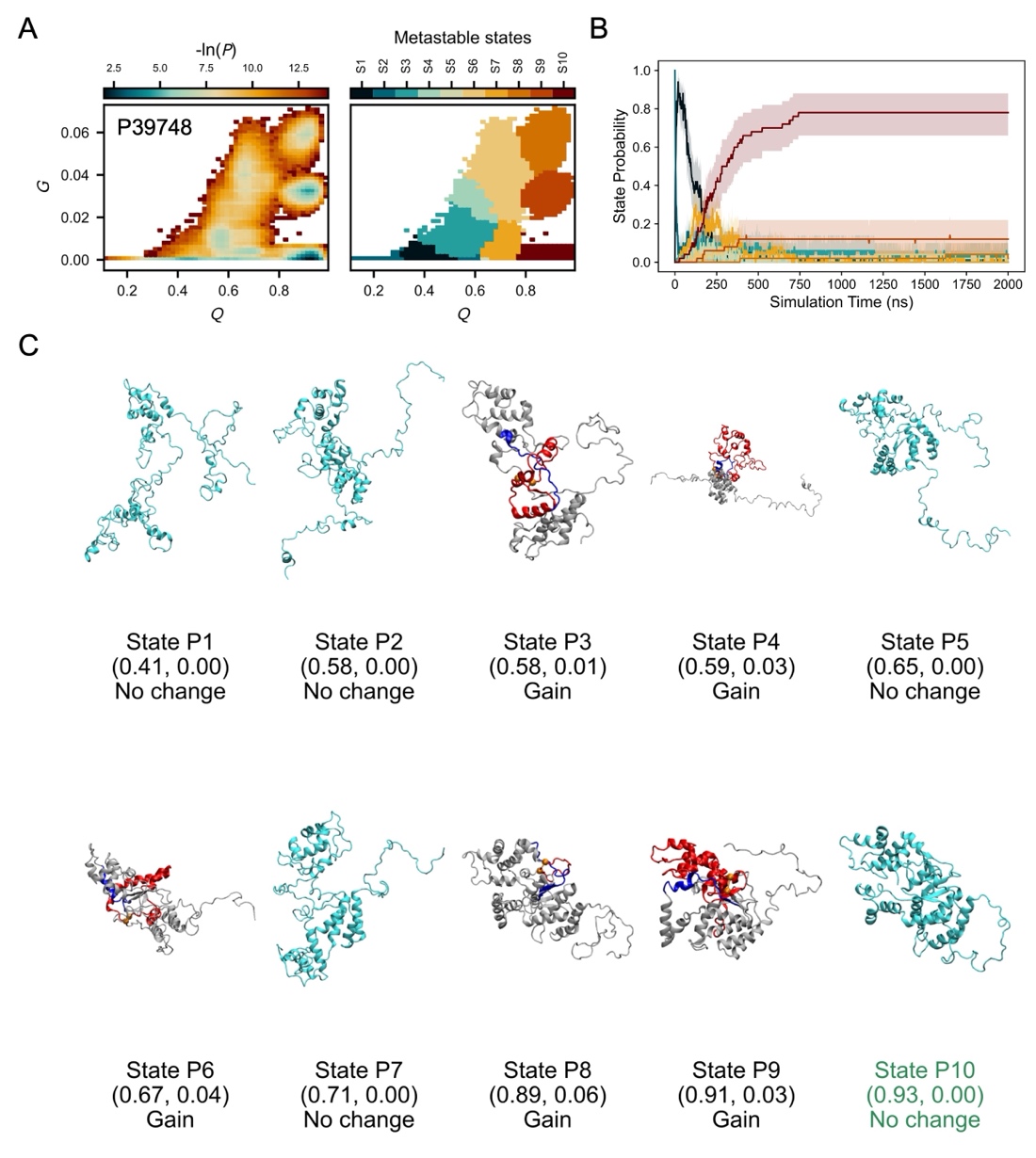

**Figure S27. Simulation structural ensemble of the NU-E protein P39748.** A. The probability distribution (-ln*P*) in the *Q* – *G* space (left) and the corresponding metastable states mapped onto structures from temperature-quench refolding simulations (right). B. Time evolution of metastable-state probabilities. Colors match the state assignments shown in panel A. Shaded regions denote 95% confidence intervals, estimated using bootstrap resampling (10^5^ iterations). C. Representative structures of each metastable state predicted from simulations. *Q* and *G* values at the cluster center, as well as the type of entanglement change (gain, loss or no change), are shown below each structure image. Color scheme follows that used in Fig. 2B, 2D, and 2F.

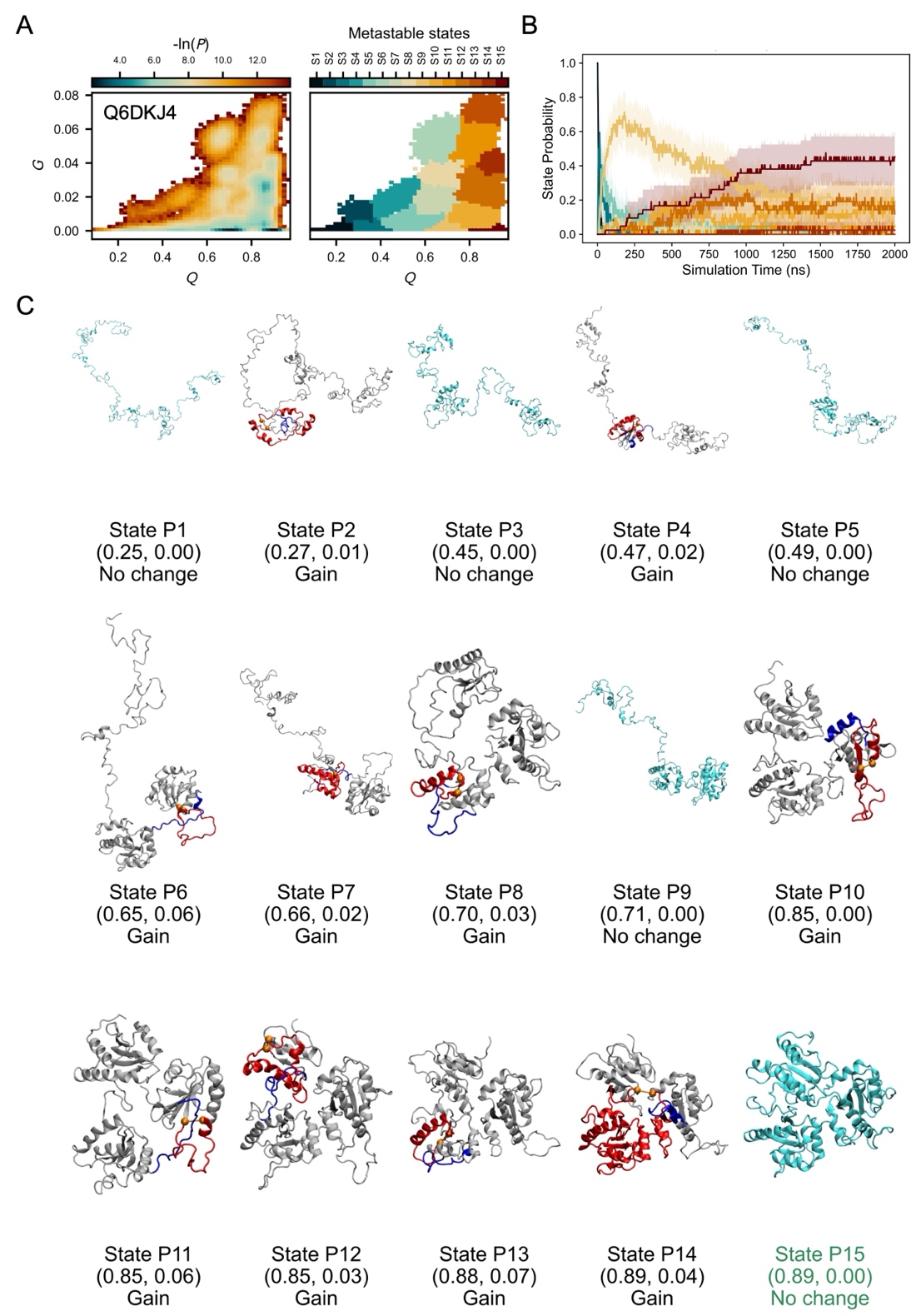

**Figure S28. Simulation structural ensemble of the NU-E protein Q6DKJ4.** A. The probability distribution (-ln*P*) in the *Q* – *G* space (left) and the corresponding metastable states mapped onto structures from temperature-quench refolding simulations (right). B. Time evolution of metastable-state probabilities. Colors match the state assignments shown in panel A. Shaded regions denote 95% confidence intervals, estimated using bootstrap resampling (10^5^ iterations). C. Representative structures of each metastable state predicted from simulations. *Q* and *G* values at the cluster center, as well as the type of entanglement change (gain, loss or no change), are shown below each structure image. Color scheme follows that used in Fig. 2B, 2D, and 2F.

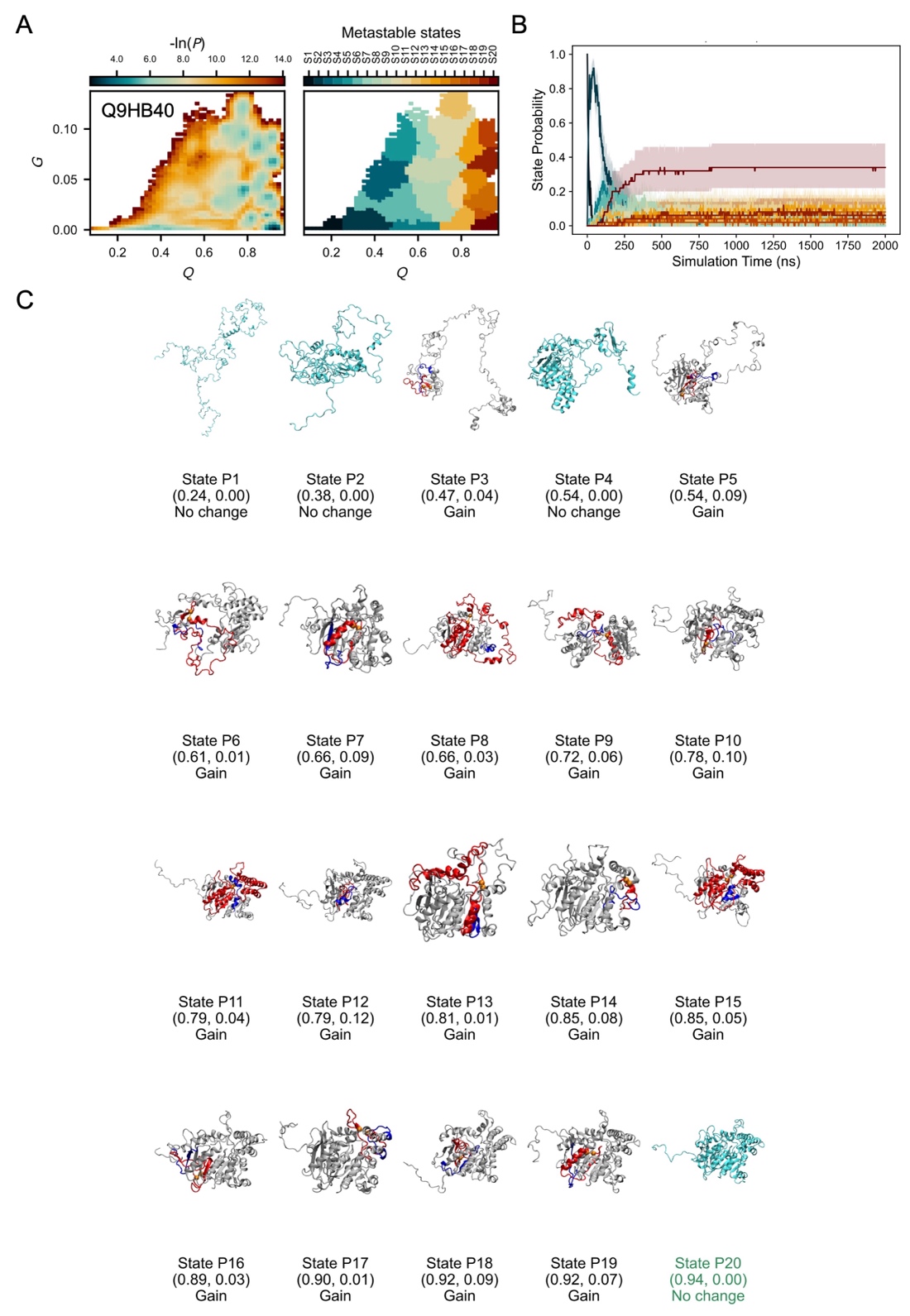

**Figure S29. Simulation structural ensemble of the NU-E protein Q9HB40.** A. The probability distribution (-ln*P*) in the *Q* – *G* space (left) and the corresponding metastable states mapped onto structures from temperature-quench refolding simulations (right). B. Time evolution of metastable-state probabilities. Colors match the state assignments shown in panel A. Shaded regions denote 95% confidence intervals, estimated using bootstrap resampling (10^5^ iterations). C. Representative structures of each metastable state predicted from simulations. *Q* and *G* values at the cluster center, as well as the type of entanglement change (gain, loss or no change), are shown below each structure image. Color scheme follows that used in Fig. 2B, 2D, and 2F.

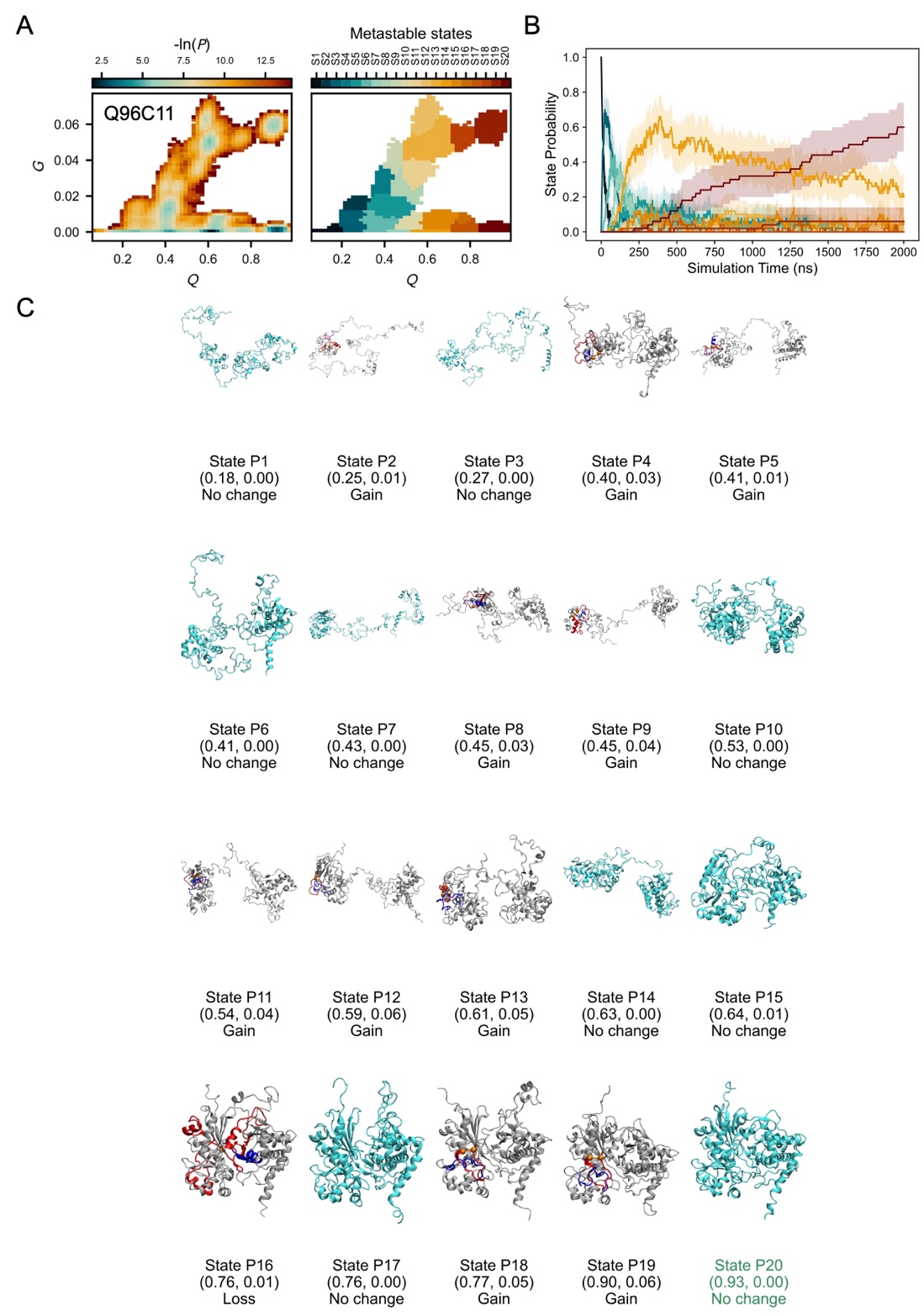

**Figure S30. Simulation structural ensemble of the NU-E protein Q96C11.** A. The probability distribution (-ln*P*) in the *Q* – *G* space (left) and the corresponding metastable states mapped onto structures from temperature-quench refolding simulations (right). B. Time evolution of metastable-state probabilities. Colors match the state assignments shown in panel A. Shaded regions denote 95% confidence intervals, estimated using bootstrap resampling (10^5^ iterations). C. Representative structures of each metastable state predicted from simulations. *Q* and *G* values at the cluster center, as well as the type of entanglement change (gain, loss or no change), are shown below each structure image. Color scheme follows that used in Fig. 2B, 2D, and 2F.

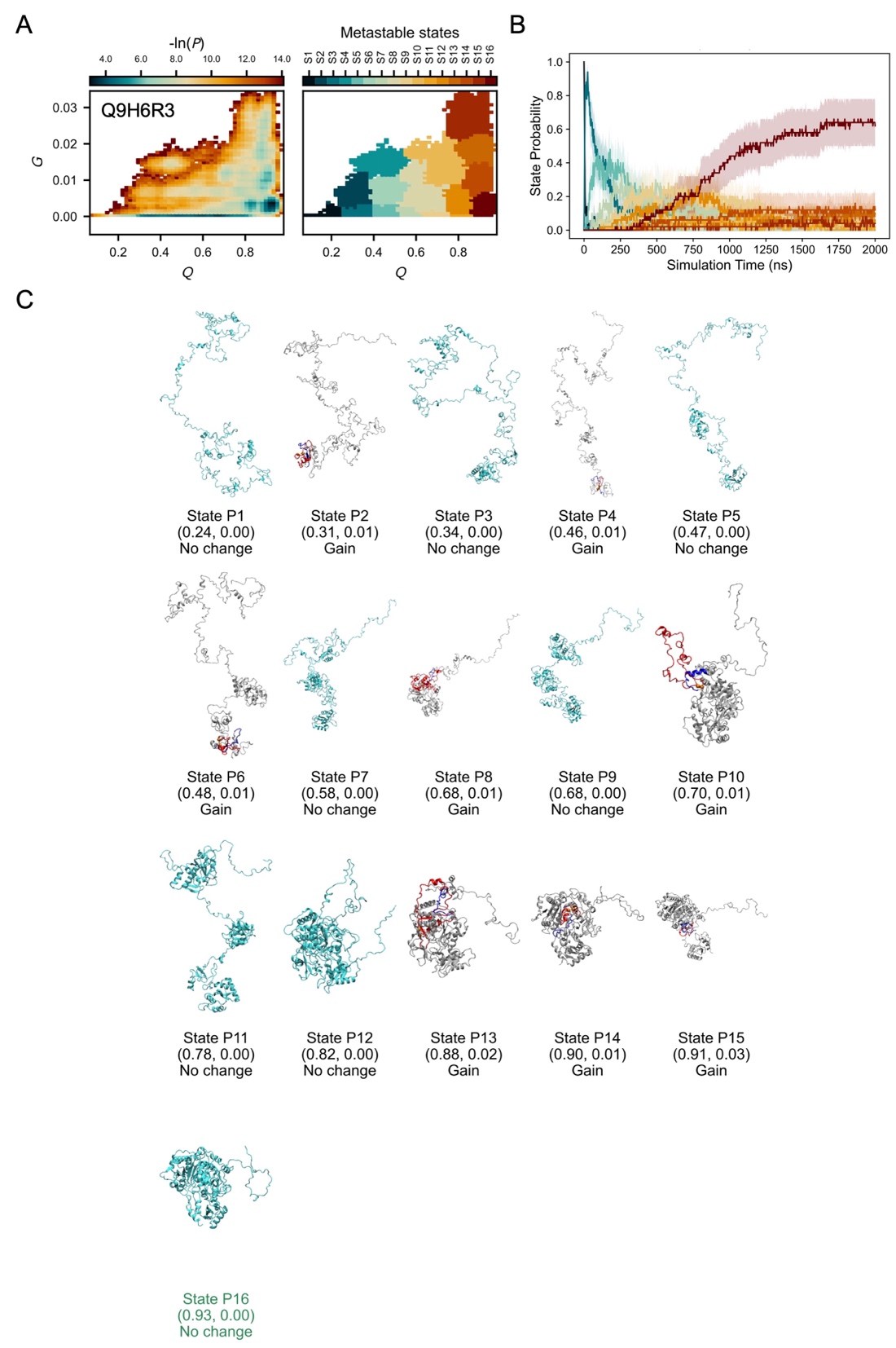

**Figure S31. Simulation structural ensemble of the NU-E protein Q9H6R3.** A. The probability distribution (-ln*P*) in the *Q* – *G* space (left) and the corresponding metastable states mapped onto structures from temperature-quench refolding simulations (right). B. Time evolution of metastable-state probabilities. Colors match the state assignments shown in panel A. Shaded regions denote 95% confidence intervals, estimated using bootstrap resampling (10^5^ iterations). C. Representative structures of each metastable state predicted from simulations. *Q* and *G* values at the cluster center, as well as the type of entanglement change (gain, loss or no change), are shown below each structure image. Color scheme follows that used in Fig. 2B, 2D, and 2F.

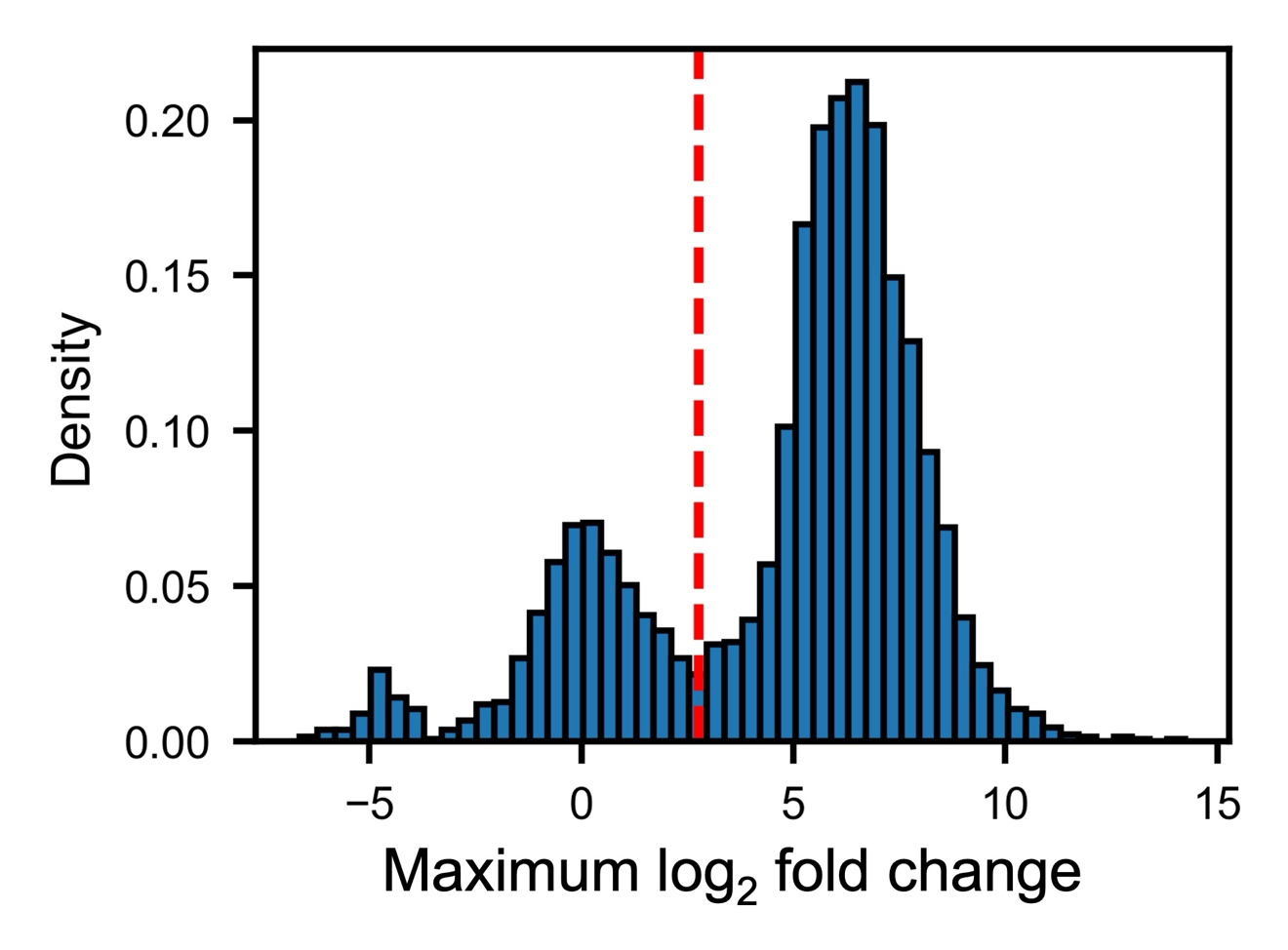

**Figure S32. Distribution of the maximum log_2_ fold change in Kε-GG peptide abundance (per protein) upon MG132-mediated proteasome inhibition in the birthdating experiments.** The red dashed line marks the threshold of 6.75, which separates the two peaks in the distribution.

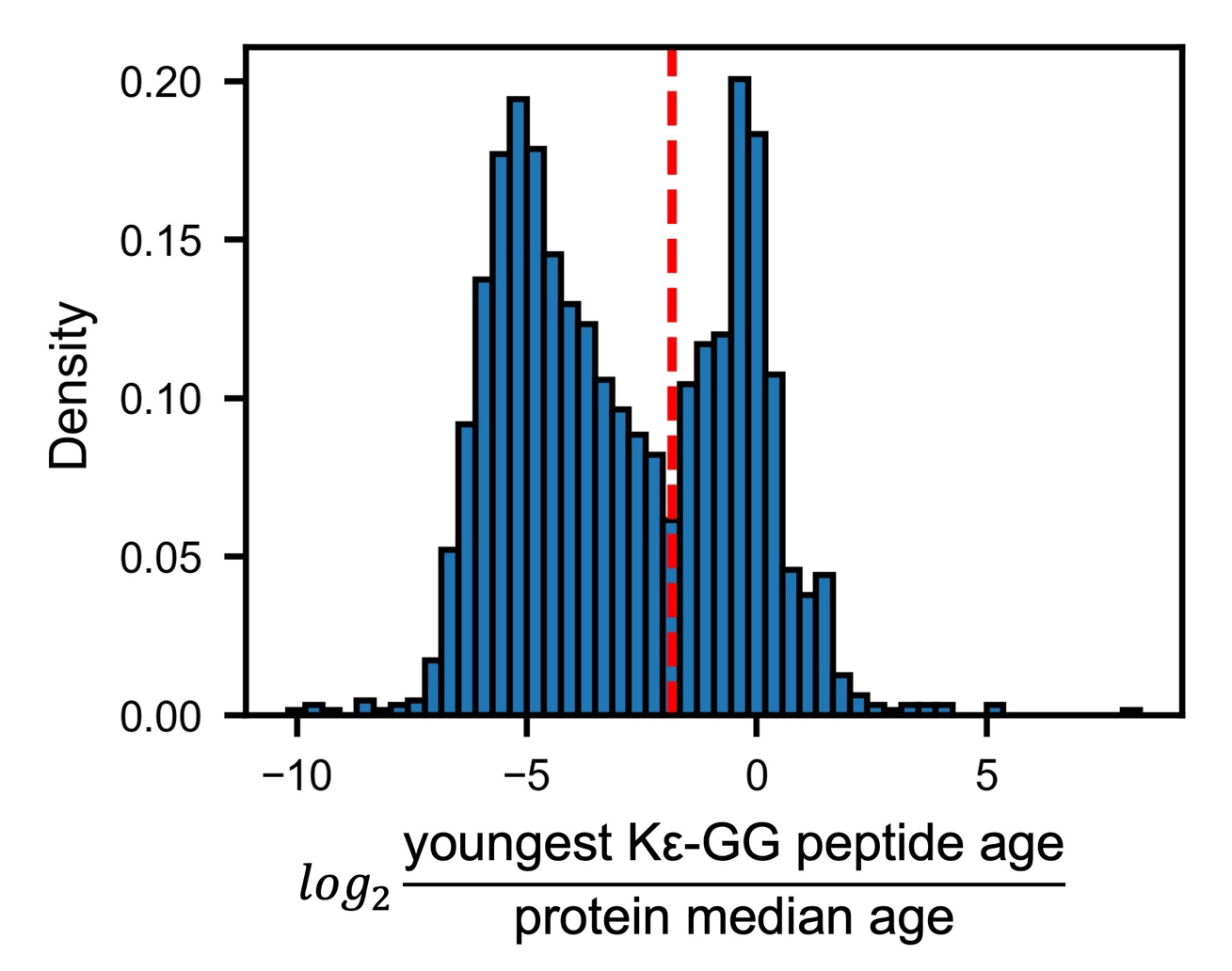

**Figure S33. Distribution of the log_2_ ratio between the age of the youngest Kε-GG peptide and the protein’s median age in the birthdating experiments.** The red dashed line marks the threshold of 0.2772, which separates the two peaks in the distribution.

**Table S1.** Contingency table for young-age ubiquitination within 6 hours (number of proteins).

|  | Young-age ubiquitination (< 6 hours) | Non-ubiquitination |
| --- | --- | --- |
| Entangled protein | 431 | 693 |
| Non-entangled protein | 72 | 294 |

**Table S2.** Contingency table for young-age ubiquitination within 1 day (number of proteins).

|  | Young-age ubiquitination (< 1 day) | Non-ubiquitination |
| --- | --- | --- |
| Entangled protein | 459 | 693 |
| Non-entangled protein | 82 | 294 |

**Table S3.** Logistic regression results for the Kim datasets.

| Dataset | IPI to Uniprot protein mapping | Sample size (number of proteins) | Predictor | O.R. | 95% CI | p-value |
| --- | --- | --- | --- | --- | --- | --- |
| Kim_btz | Legacy official mapping | 684 | NCLE | 1.49 | [1.02, 2.17] | 0.0392 |
|  |  |  | Protein Length | 1.39 | [1.11, 1.75] | 0.0042 |
|  | Gene name & motif search | 839 | NCLE | 1.85 | [1.33, 2.59] | 0.0003 |
|  |  |  | Protein Length | 1.47 | [1.19, 1.81] | 0.0003 |
| Kim_epox | Legacy official mapping | 499 | NCLE | 1.17 | [0.75, 1.82] | 0.4967 |
|  |  |  | Protein Length | 1.42 | [1.08, 1.87] | 0.0121 |
|  | Gene name & motif search | 633 | NCLE | 1.36 | [0.92, 2.00] | 0.1192 |
|  |  |  | Protein Length | 1.44 | [1.12, 1.84] | 0.0039 |

**Table S4.** *Q*_norm_ of the misfolded entanglement states of YU-E and NU-E proteins.

| YU-E proteins | | | NU-E proteins | | |
| --- | --- | --- | --- | --- | --- |
| Uniprot ID | Number of trajectories w/o mirror images | *Q*_norm_ [95% CI] | Uniprot ID | Number of trajectories w/o mirror images | *Q*_norm_ [95% CI] |
| P31150 | 50 | 0.80 [0.74, 0.85] | Q9HB40 | 50 | 0.91 [0.89, 0.94] |
| Q0PNE2 | 50 | 0.93 [0.91, 0.96] | P07738 | 50 | 0.93 [0.90, 0.96] |
| A8MXV4 | 50 | 0.79 [0.75, 0.82] | P39748 | 50 | 0.90 [0.81, 0.98] |
| P16152 | 50 | 0.57 [0.56, 0.59] | Q9UIV1 | 50 | 0.99 [0.99, 0.99] |
| P00491 | 50 | 0.69 [0.59, 0.81] | Q9Y316 | 50 | 0.97 [0.96, 0.98] |
| O95394 | 50 | 0.98 [0.96, 0.99] | Q96C11 | 50 | 0.73 [0.69, 0.78] |
| P04350 | 50 | 0.57 [0.57, 0.57] | Q6DKJ4 | 42 | 0.98 [0.96, 1.00] |
| P19623 | 44 | 0.94 [0.93, 0.95] | Q63HM1 | 50 | 0.98 [0.97, 0.99] |
| P29218 | 50 | 0.88 [0.83, 0.92] | Q9UBP6 | 50 | 0.91 [0.90, 0.92] |
| P52888 | 50 | 0.82 [0.72, 0.93] | Q9H6R3 | 50 | 0.96 [0.95, 0.98] |
| Average | 494 | 0.80 [0.71, 0.88] | Average | 492 | 0.93 [0.87, 0.97] |

**Table S5.** rSASA of the misfolded entanglement states of YU-E and NU-E proteins.

| YU-E proteins | | | NU-E proteins | | |
| --- | --- | --- | --- | --- | --- |
| Uniprot ID | Number of trajectories w/o mirror images | rSASA [95% CI] | Uniprot ID | Number of trajectories w/o mirror images | rSASA [95% CI] |
| P31150 | 50 | 1.67 [1.48, 1.86] | Q9HB40 | 50 | 1.11 [1.07, 1.15] |
| Q0PNE2 | 50 | 1.06 [1.02, 1.10] | P07738 | 50 | 1.23 [1.17, 1.31] |
| A8MXV4 | 50 | 1.23 [1.18, 1.29] | P39748 | 50 | 1.21 [1.02, 1.41] |
| P16152 | 50 | 2.07 [2.02, 2.11] | Q9UIV1 | 50 | 1.03 [1.03, 1.03] |
| P00491 | 50 | 1.63 [1.37, 1.87] | Q9Y316 | 50 | 1.09 [1.07, 1.11] |
| O95394 | 50 | 1.04 [1.01, 1.08] | Q96C11 | 50 | 1.57 [1.45, 1.66] |
| P04350 | 50 | 2.08 [2.07, 2.09] | Q6DKJ4 | 42 | 0.99 [0.95, 1.03] |
| P19623 | 44 | 1.03 [1.00, 1.05] | Q63HM1 | 50 | 1.04 [1.03, 1.04] |
| P29218 | 50 | 1.23 [1.14, 1.34] | Q9UBP6 | 50 | 1.02 [1.00, 1.04] |
| P52888 | 50 | 1.41 [1.10, 1.62] | Q9H6R3 | 50 | 1.05 [1.01, 1.08] |
| Average | 494 | 1.45 [1.21, 1.70] | Average | 492 | 1.13 [1.04, 1.25] |

**Table S6.** The structural domains and the force field parameters ($n_{\text{scal}}$) of the 10 YU-E proteins (* denotes the instable domain or interface that cannot be stabilized using the highest level of $n_{\text{scal}}$ and the median value was used.)

| Uniprot ID | Name | Length | Domain and Interfaces | Structural Class | $n_{\text{scal}}$ |
| --- | --- | --- | --- | --- | --- |
| Q0PNE2 | Elongator complex protein 6 | 266 | Domain 1: 1 – 266 | α/β | 1.1556 |
| P16152 | Carbonyl reductase [NADPH] 1 | 277 | Domain 1: 1 – 277 | α/β | 1.1556 |
| P29218 | Inositol monophosphatase 1 | 277 | Domain 1: 1 – 148 | α/β | 1.1556 |
|  |  |  | Domain 2: 149 – 277 | α/β | 1.4213 |
|  |  |  | 1 \| 2 Interface | – | 1.2747 |
| P00491 | Purine nucleoside phosphorylase | 289 | Domain 1: 1 – 289 | α/β | 1.1556 |
| P19623 | Spermidine synthase | 302 | Domain 1: 1 – 73 | β | 1.4732 |
|  |  |  | Domain 2: 74 – 302 | α/β | 1.1556 |
|  |  |  | 1 \| 2 Interface | – | 2.1670 |
| A8MXV4 | Acyl-coenzyme A diphosphatase NUDT19 | 375 | Domain 1: 1 – 292 | α/β | 1.1556 |
|  |  |  | Domain 2: 293 – 375 | α/β | 1.1556 |
|  |  |  | 1 \| 2 Interface | – | 1.8611 |
| P04350 | Tubulin beta-4A chain | 444 | Domain 1: 1 – 260 | α/β | 1.1556 |
|  |  |  | Domain 2: 261 – 371 | α/β | 1.1556 |
|  |  |  | Domain 3: 372 – 444 | α | 1.1954 |
|  |  |  | 1 \| 2 Interface | – | 1.2747 |
|  |  |  | 1 \| 3 Interface | – | 1.2747 |
|  |  |  | 2 \| 3 Interface | – | 2.5044 |
| P31150 | Rab GDP dissociation inhibitor alpha | 447 | Domain 1: 1 – 41, 231 – 288, 392 – 447 | α/β | 1.1556 |
|  |  |  | Domain 2: 42 – 118, 221 – 230, 289 – 391 | α/β | 1.1556 |
|  |  |  | Domain 3: 119 – 220 | α | 1.1954 |
|  |  |  | 1 \| 2 Interface | – | 1.2747 |
|  |  |  | 1 \| 3 Interface | – | 1.2747 |
|  |  |  | 2 \| 3 Interface | – | 1.2747 |
| O95394 | Phosphoacetylglucosamine mutase | 542 | Domain 1: 1 – 175 | α/β | 1.1556 |
|  |  |  | Domain 2: 176 – 297 | α/β | 1.1556 |
|  |  |  | Domain 3: 298 – 443 | α/β | 1.1556 |
|  |  |  | Domain 4: 444 – 542 | α/β | 1.4213 |
|  |  |  | 1 \| 2 Interface | – | 1.2747 |
|  |  |  | 1 \| 3 Interface | – | 1.2747 |
|  |  |  | 1 \| 4 Interface | – | 1.2747 |
|  |  |  | 2 \| 3 Interface | – | 1.2747 |
|  |  |  | 2 \| 4 Interface | – | 1.2747 |
|  |  |  | 3 \| 4 Interface | – | 1.8611* |
| P52888 | Thimet oligopeptidase | 689 | Domain 1: 1 – 154 | α | 1.1954 |
|  |  |  | Domain 2: 187 – 246 | α/β | 1.1556 |
|  |  |  | Domain 3: 361 – 545, 608 – 689 | α/β | 1.1556 |
|  |  |  | Domain 4: 155 – 186, 247 – 360, 546 – 607 | α | 1.1954 |
|  |  |  | 1 \| 2 Interface | – | 1.2747 |
|  |  |  | 1 \| 3 Interface | – | 1.2747 |
|  |  |  | 1 \| 4 Interface | – | 1.2747 |
|  |  |  | 2 \| 3 Interface | – | 1.2747 |
|  |  |  | 2 \| 4 Interface | – | 1.2747 |
|  |  |  | 3 \| 4 Interface | – | 1.2747 |

**Table S7.** The structural domains and the force field parameters ($n_{\text{scal}}$) of the 10 NU-NE proteins (* denotes the instable domain or interface that cannot be stabilized using the highest level of $n_{\text{scal}}$ and the median value was used.)

| Uniprot ID | Name | Length | Domain and Interfaces | Structural Class | $n_{\text{scal}}$ |
| --- | --- | --- | --- | --- | --- |
| Q8WV22 | Non-structural maintenance of chromosomes element 1 homolog | 266 | Domain 1: 1 – 90 | α/β | 1.4213 |
|  |  |  | Domain 2: 91 – 185 | α | 1.1954 |
|  |  |  | Domain 3: 186 – 266 | α/β | 1.6871* |
|  |  |  | 1 \| 2 Interface | – | 1.8611* |
|  |  |  | 1 \| 3 Interface | – | 1.2747 |
|  |  |  | 2 \| 3 Interface | – | 1.2747 |
| A6NDU8 | RAB7A-interacting MON1-CCZ1 complex subunit 1 | 294 | Domain 1: 1 – 122 | α | 1.1954 |
|  |  |  | Domain 2: 123 – 294 | α | 1.1954 |
|  |  |  | 1 \| 2 Interface | – | 1.2747 |
| P30711 | Glutathione S-transferase theta-1 | 240 | Domain 1: 1 – 81 | α/β | 1.1556 |
|  |  |  | Domain 2: 82 – 240 | α | 1.1954 |
|  |  |  | 1 \| 2 Interface | – | 1.2747 |
| Q9BU89 | Deoxyhypusine hydroxylase | 302 | Domain 1: 1 – 139 | α | 1.1954 |
|  |  |  | Domain 2: 140 – 302 | α | 1.1954 |
|  |  |  | 1 \| 2 Interface | – | 1.5679 |
| Q13825 | Methylglutaconyl-CoA hydratase, mitochondrial | 339 | Domain 1: 1 – 67 | α/β | 1.1556 |
|  |  |  | Domain 2: 68 – 279 | α/β | 1.4213 |
|  |  |  | Domain 3: 280 – 339 | α | 1.7453* |
|  |  |  | 1 \| 2 Interface | – | 1.8611* |
|  |  |  | 1 \| 3 Interface | – | 1.2747 |
|  |  |  | 2 \| 3 Interface | – | 1.2747 |
| Q6NVY1 | 3-hydroxyisobutyryl-CoA hydrolase, mitochondrial | 386 | Domain 1: 1 – 30 | α | 1.7453* |
|  |  |  | Domain 2: 31 – 386 | α/β | 1.1556 |
|  |  |  | 1 \| 2 Interface | – | 1.2747 |
| P16520 | Guanine nucleotide-binding protein G(I)/G(S)/G(T) subunit beta-3 | 340 | Domain 1: 1 – 340 | β | 1.4732 |
| Q8IV38 | Ankyrin repeat and MYND domain-containing protein 2 | 441 | Domain 1: 1 – 135 | α | 1.4704 |
|  |  |  | Domain 2: 136 – 311 | α | 1.1954 |
|  |  |  | Domain 3: 312 – 441 | α/β | 1.6871* |
|  |  |  | 1 \| 2 Interface | – | 1.8611* |
|  |  |  | 1 \| 3 Interface | – | 1.2747 |
|  |  |  | 2 \| 3 Interface | – | 2.5044 |
| P02774 | Vitamin D-binding protein | 474 | Domain 1: 1 – 126 | α | 1.4704 |
|  |  |  | Domain 2: 127 – 226 | α | 1.1954 |
|  |  |  | Domain 3: 227 – 409 | α | 1.1954 |
|  |  |  | Domain 4: 410 – 474 | α | 1.1954 |
|  |  |  | 1 \| 2 Interface | – | 1.2747 |
|  |  |  | 1 \| 3 Interface | – | 1.2747 |
|  |  |  | 1 \| 4 Interface | – | 1.2747 |
|  |  |  | 2 \| 3 Interface | – | 1.2747 |
|  |  |  | 2 \| 4 Interface | – | 1.5679 |
|  |  |  | 3 \| 4 Interface | – | 1.2747 |
| Q12996 | Cleavage stimulation factor subunit 3 | 717 | Domain 1: 1 – 233 | α | 1.1954 |
|  |  |  | Domain 2: 234 – 337 | α | 1.1954 |
|  |  |  | Domain 3: 338 – 558 | α | 1.1954 |
|  |  |  | Domain 4: 559 – 717 | α/β | 1.6871* |
|  |  |  | 1 \| 2 Interface | – | 2.1670 |
|  |  |  | 1 \| 3 Interface | – | 1.2747 |
|  |  |  | 1 \| 4 Interface | – | 1.2747 |
|  |  |  | 2 \| 3 Interface | – | 1.2747 |
|  |  |  | 2 \| 4 Interface | – | 1.5679 |
|  |  |  | 3 \| 4 Interface | – | 1.2747 |

**Table S8.** The structural domains and the force field parameters ($n_{\text{scal}}$) of the 10 NU-NE proteins (* denotes the instable domain or interface that cannot be stabilized using the highest level of $n_{\text{scal}}$ and the median value was used.)

| Uniprot ID | Name | Length | Domain and Interfaces | Structural Class | $n_{\text{scal}}$ |
| --- | --- | --- | --- | --- | --- |
| P07738 | Bisphosphoglycerate mutase | 259 | Domain 1: 1 – 259 | α/β | 1.1556 |
| Q9UIV1 | CCR4-NOT transcription complex subunit 7 | 285 | Domain 1: 1 – 285 | α/β | 1.1556 |
| Q9UBP6 | tRNA (guanine-N(7)-)-methyltransferase | 276 | Domain 1: 1 – 276 | α/β | 1.1556 |
| Q9Y316 | Protein MEMO1 | 297 | Domain 1: 1 – 297 | α/β | 1.1556 |
| Q63HM1 | Kynurenine formamidase | 303 | Domain 1: 1 – 303 | α/β | 1.1556 |
| P39748 | Flap endonuclease 1 | 380 | Domain 1: 1 – 216 | α/β | 1.1556 |
|  |  |  | Domain 2: 217 – 285 | α | 1.1954 |
|  |  |  | Domain 3: 286 – 380 | α | 1.1954 |
|  |  |  | 1 \| 2 Interface | – | 1.2747 |
|  |  |  | 1 \| 3 Interface | – | 1.2747 |
|  |  |  | 2 \| 3 Interface | – | 1.2747 |
| Q6DKJ4 | Nucleoredoxin | 435 | Domain 1: 1 – 169 | α/β | 1.1556 |
|  |  |  | Domain 2: 170 – 311 | α/β | 1.1556 |
|  |  |  | Domain 3: 312 – 435 | α/β | 1.1556 |
|  |  |  | 1 \| 2 Interface | – | 1.8611* |
|  |  |  | 1 \| 3 Interface | – | 1.2747 |
|  |  |  | 2 \| 3 Interface | – | 1.2747 |
| Q9HB40 | Retinoid-inducible serine carboxypeptidase | 452 | Domain 1: 1 – 204, 347 – 452 | α/β | 1.1556 |
|  |  |  | Domain 2: 205 – 346 | α | 1.1954 |
|  |  |  | 1 \| 2 Interface | – | 1.2747 |
| Q96C11 | FGGY carbohydrate kinase domain-containing protein | 551 | Domain 1: 1 – 257 | α/β | 1.1556 |
|  |  |  | Domain 2: 258 – 333, 438 – 551 | α/β | 1.1556 |
|  |  |  | Domain 3: 334 – 437 | α/β | 1.1556 |
|  |  |  | 1 \| 2 Interface | – | 1.2747 |
|  |  |  | 1 \| 3 Interface | – | 1.2747 |
|  |  |  | 2 \| 3 Interface | – | 1.2747 |
| Q9H6R3 | Acyl-CoA synthetase short-chain family member 3, mitochondrial | 686 | Domain 1: 1 – 556 | α/β | 1.1556 |
|  |  |  | Domain 2: 557 – 686 | α/β | 1.4213 |
|  |  |  | 1 \| 2 Interface | – | 1.8611* |
